# A general strategy to develop fluorogenic polymethine dyes for bioimaging

**DOI:** 10.1101/2023.01.31.526423

**Authors:** Annabell Martin, Pablo Rivera-Fuentes

## Abstract

Fluorescence imaging is an invaluable tool to study biological processes and further progress depends on the development of advanced probes. Fluorogenic dyes are crucial to reach intracellular targets and label them with high specificity. Excellent fluorogenic rhodamine dyes have been reported, but they often require a long and low-yielding synthesis and are spectrally limited to the visible range. Here, we present a general strategy to transform polymethine compounds into fluorogenic dyes using an intramolecular ring closure approach. We illustrate the generality of this method by creating both spontaneously blinking and no-wash, turn-on polymethine dyes with emissions across the visible and near-infrared spectrum. These probes are compatible with self-labeling proteins and small-molecule targeting ligands and can be combined with rhodamine-based dyes for multicolor and fluorescence lifetime multiplexing imaging. This strategy provides access to bright, fluorogenic dyes that emit at wavelengths that are significantly more red-shifted than those of existing rhodamine-based dyes.

## Main

Fluorescence microscopy is crucial to study the structure and function of cells. Fluorescent protein tags allow for the dynamic observation of proteins in living cells, but their brightness and photostability are often inferior compared to those of small-molecule fluorophores^1^. Fluorogenic dyes conjugated to self-labeling protein tags such as HaloTag^2^ or SNAP-tag^3^ combine the excellent photophysical properties of smallmolecule dyes with the precise labeling of genetically encoded tags and have been widely utilized for fluorescence microscopy and nanoscopy.

The development of fluorogenic probes has predominantly focused on rhodamine-based scaffolds^4,5^. Rhodamine dyes exist in an equilibrium between a fluorescent zwitterionic (open) and a non-fluorescent spirocyclic (closed) form. This equilibrium is sensitive to the microenvironment of the probe. In media of low polarity and at high pH, rhodamines are primarily present in the closed form, whereas polar media and low pH favor the open form^6–8^. Rhodamine dyes have been systematically optimized for fluorogenicity by varying the electron density of their conjugated core and by modulating the electronic character of the intramolecular nucleophile that closes the spirocycle^9,10^. These dyes, which span the visible range, display significant fluorescence increase upon target binding and have been used for no-wash, multicolor, live-cell fluorescence imaging experiments. In contrast, the development of novel rhodamines with excitation and emission wavelengths in the near-infrared region (>700 nm) has been hampered by their complicated, long, and low-yielding syntheses, as well as by their tendency to remain in the non-fluorescent form even upon binding to their target^11^.

Polymethine dyes are some of the most used fluorophores in cell, tissue, and whole-organism imaging due to their simple and highly modular synthesis, high extinction coefficients, biocompatibility, and tunable emission wavelengths spanning from the green to the shortwave infrared wavelengths. Members of this class of fluorophores include indoleninium-based dyes such as carbocyanines (Cy)^12–15^ and squaraines^16–18^, flavylium-based dyes^19,20^, and coumarin-hemicyanine hybrid scaffolds^21–24^, among others. The variety of applications of these dyes is illustrated by two classic examples: Indocyanine Green^25^, a clinically-approved, near-infrared dye for optical imaging of the vasculature, and Cy5 derivatives (e. g., AlexaFluor^®^ 647), which are the most widely used fluorophores in single-molecule localization microscopy (SMLM)^26^. Unfortunately, there is no general strategy to impart binding-induced fluorogenicity to polymethine dyes and thus, their applicability in live-cell imaging of specific protein targets remains very limited.

Unlike rhodamine-based dyes, polymethine fluorophores do not possess a built-in intramolecular cyclization equilibrium that could be leveraged to induce fluorogenicity. Through the addition of nucleophilic side chains, however, several research groups have attempted to create fluorogenic polymethine dyes. The Ohe group decorated Cy5 and Cy7 dyes with alcohols, amines, and thiols as nucleophiles, thereby forming oxazines, diazines, and thiazines, respectively^27–30^. The Raymo group developed coumarin-hemicyanine hybrid fluorophores bearing a *p*-nitrophenol group that exists mainly in the spirocyclic form but can undergo a photoinduced and reversible interconversion into its open fluorescent form^24^. Furthermore, joint efforts from the Johnsson and Urano groups utilized coumarin-hemicyanine hybrid fluorophores as well, but with a hydroxyethyl ring-closing moiety to generate esterase-activatable fluorescent probes^21^. Despite these efforts, cyclization reactions in polymethine dyes have not been as efficient as in rhodamines, and the creation of robustly fluorogenic polymethine dyes has been elusive. In this work, we applied classic organic chemistry heuristics to design favorable cyclization reactions that led to a general strategy for the creation of fluorogenic polymethine dyes for live-cell imaging.

## Results and Discussion

### Probe design

An efficient fluorogenic dye must exist exclusively in the non-fluorescent form unless it is bound to its intended target. In the case of cyclization-based fluorogenicity, it means that the barrier of ring opening must be high, and the barrier of ring closure must be low (Extended Data Fig. 1). The favorability of polar cyclization reactions can be estimated using heuristic guidelines known as the Baldwin rules (Extended Data Fig. 1)^31^. We noticed that in previous attempts to create fluorogenic polymethine dyes, the intramolecular cyclization was of the 5-*endo-trig* type, which is unfavorable according to the Baldwin rules (Extended Data Fig. 1). This cyclization (Fig. 1a) is classified as 5-*endo-trig* because it involves the formation of a 5-membered ring that contains (*endo*) the single bond formed from the double bond of the trigonal (*trig*) carbon attacked by the nucleophile (Fig. 1a). Although 6-*endo-trig* cyclizations should be more favorable than 5-*endo-trig*, previous attempts at using 6-*endo-trig* cyclizations to impart fluorogenicity did not lead to more stable cyclic isomers^27–30^. Based on these observations, we hypothesized that a 5-*exo-trig* cyclization would be a more efficient alternative. 5-*exo-trig* cyclizations are those in which the single bond that is formed upon attack of the double bond is not contained (*exo*) in the newly formed ring (Fig. 1b).

**Fig. 1:**
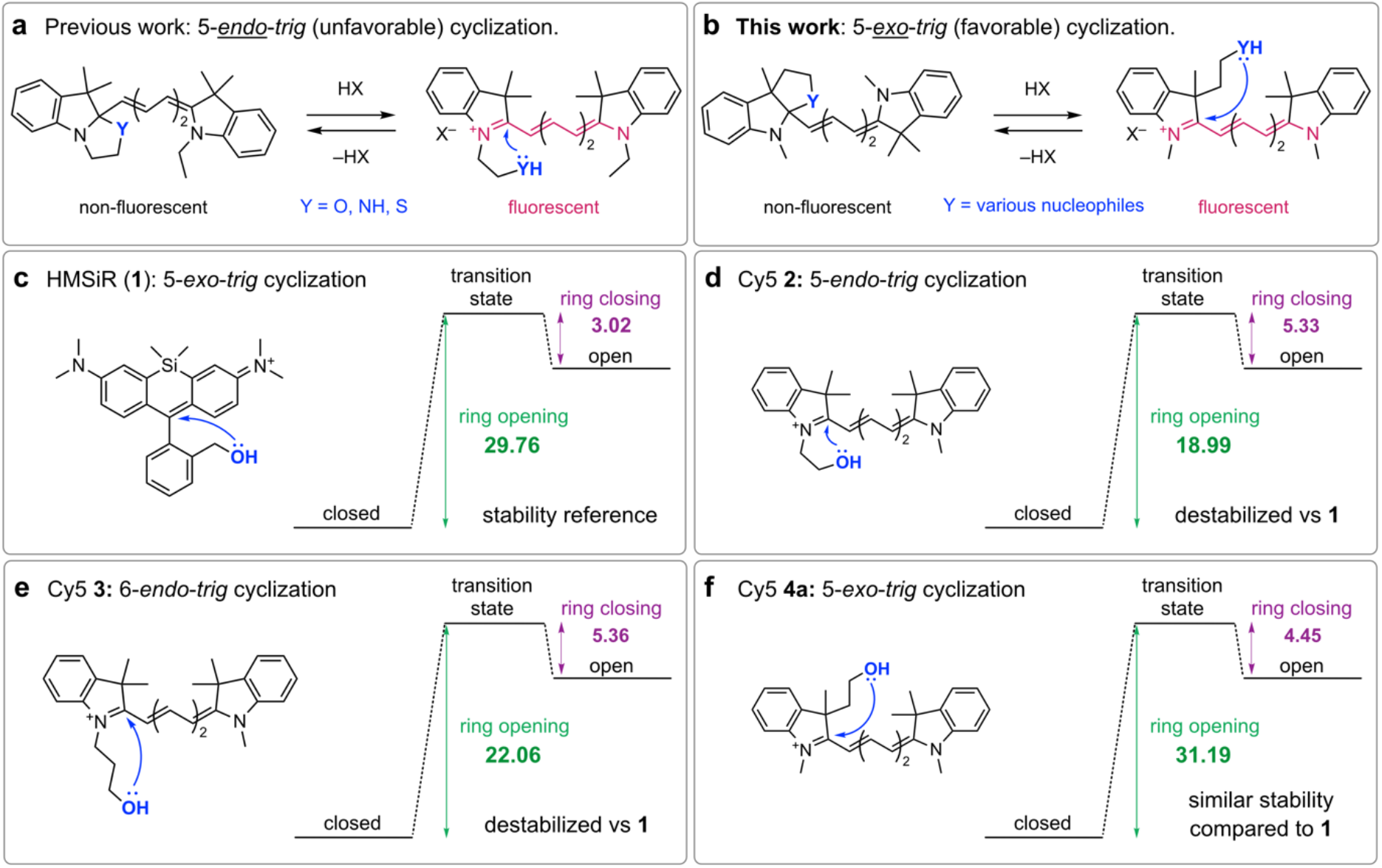
Intramolecular cyclization reactions of carbocyanines and HMSiR (1). a) Example of previous work using a 5-*endo-trig* cyclization to create fluorogenic carbocyanine dyes. b) 5-*exo-trig* cyclization as a robust and general strategy towards fluorogenicity in polymethine dyes. Structures and calculated energies of ring opening and closing of rhodamine HMSiR (**1**) (c), and Cy5 derivatives **2** (d), **3** (e), and **4a** (f). Energies were calculated at the B3LYP/DGTZVP level of theory and are reported in kcal mol^-1^. A summary of calculated energies using the B3LYP or M06-2X functional can be found in Supplementary Table 1.

To test our hypothesis, we used density functional theory (DFT, Methods) to calculate the energies of ring opening and closure for HMSiR (**1**), a silicon rhodamine derivative with an efficient 5-*exo-trig* intramolecular cyclization reaction (Fig. 1c)^32^. We then compared these values to those calculated for Cy5 derivatives **2**, **3**, and **4a**, which undergo *5-endo-trig, 6-endo-trig*, or 5-*exo-trig* cyclizations (Fig. 1d-f). HMSiR (**1**) has a larger ring-opening and smaller ring-closing energies than Cy5 derivatives **2** and **3**, which undergo 5-*endo-trig* and *6-endo-trig* cyclizations, respectively (Fig. 1c-d). In contrast, the energies of ring opening and closure in Cy5 derivative **4a**, which undergoes a 5-*exo-trig* cyclization, compare favorably with the computed values for HMSiR (**1**)(Fig. 1c,f). Furthermore, we observed that judicious addition of electron-withdrawing groups to the Cy5 dye further increased the stability of the closed isomer (e. g., derivative **4b,** Extended Data Fig. 1), allowing for additional modulation of the energies of ring opening and closure. These predictions suggested that simply by employing a 5-*exo-trig* cyclization instead of a 5-*endo-trig* one, more robust fluorogenic polymethine dyes could be obtained.

To test these computational predictions, we prepared the indoleninium building blocks **5**, **6**, **7a-d**, and linker **8** (Fig. 2a and Supplementary Information) according to published procedures^33–36^. Next, we synthesized the 5-*endo-trig* Cy5 probes **2a** and **2b** in a two-step, microwave-assisted procedure (Supplementary Information). The 5-*exo-trig* Cy5 probes **4a** and **4b** were prepared in two steps through the methyl ester intermediates **9a**-**d** followed by reduction with LiAlH4 (Fig. 2a). We performed pH titrations of these Cy5 derivatives to evaluate the stability of the closed isomer (Fig. 2b). The pH titrations of 5-*endo-trig* probes revealed ring-opening *pK_a_* values above physiological pH (11.1 for **2a** and 8.5 for **2b**), indicating that these probes would be highly fluorescent as free small molecules in cells. In contrast, the ring-opening *pK_a_* values of 5-*exo-trig* probes were below 7 (6.4 for **4a** and 5.7 for **4b**), suggesting that they might only show some fluorescence in acidic compartments. In live-cell imaging experiments (Fig. 2c), the 5-*endo-trig* probes **2a** and **2b** exhibited bright fluorescence in mitochondria (Fig. 2c and Extended Data Fig. 2). This subcellular localization is likely a consequence of the delocalized positive charge of the open Cy5 scaffold. In contrast, 5-*exo-trig* probes **4a** and **4b** showed only very faint fluorescence, mostly localized in lysosomes (Fig. 2c and Extended Data Fig. 2). These results confirm that the closed form of probes that undergo 5-*exo-trig* cyclizations (**4a** and **4b**) is very stable under physiological conditions and greatly minimizes non-specific background in live-cell imaging.

**Fig. 2:**
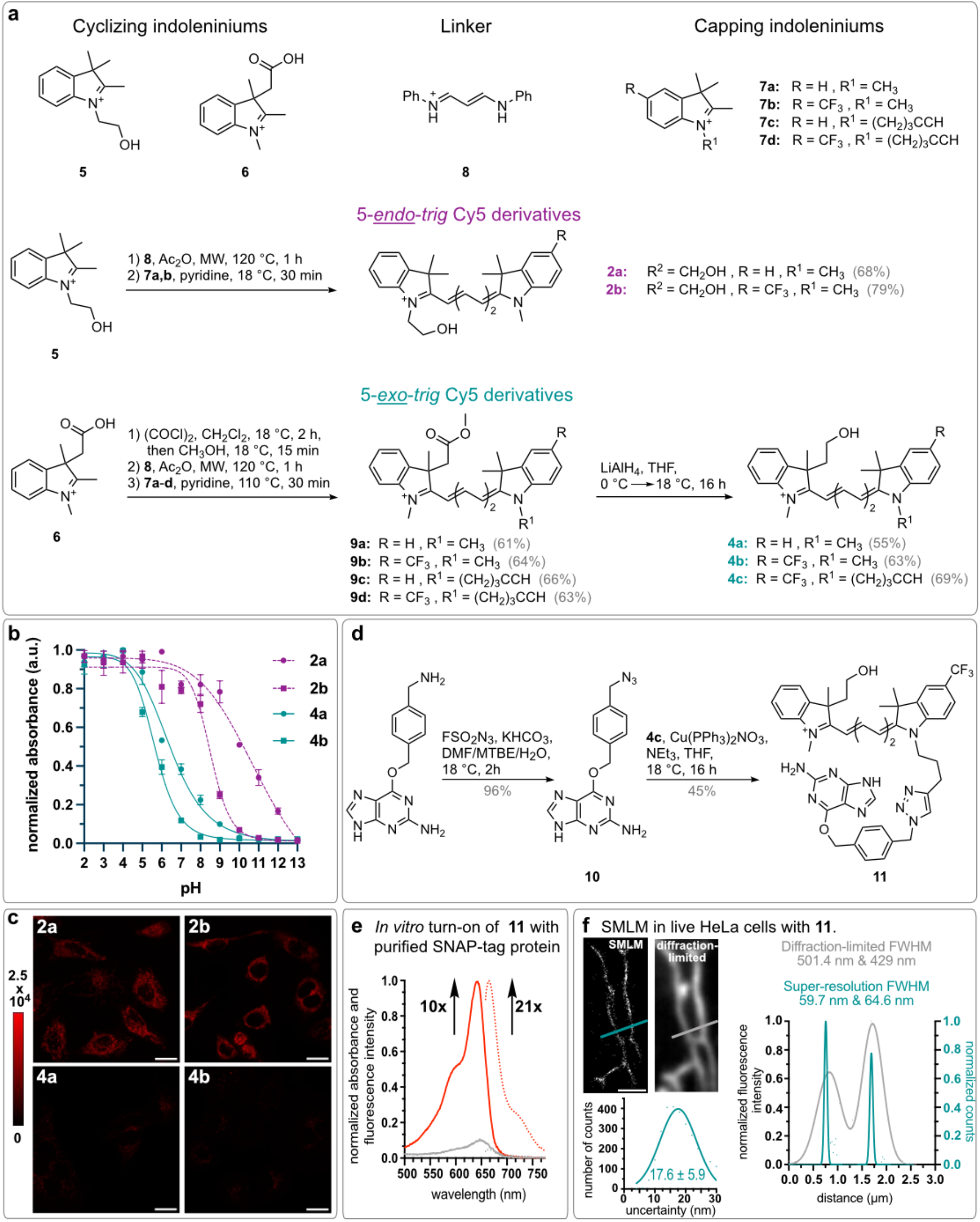
Synthesis and imaging properties of 5-*endo-trig* and 5-*exo-trig* Cy5 derivatives. a) Synthesis of 5-*endo-trig* and 5-*exo-trig* Cy5 dyes with a hydroxy group as a ring-closing moiety. b) pH profiles of dyes **2a**, **2b**, **4a**, and **4b**. c) Fluorescence of untargeted 5-*endo-trig* (**2a**,**b**) and 5-*exo-trig* Cy5 derivatives (**4a,b**). The calibration bar indicates raw pixel values. d) SNAP-tag functionalization of **4c** to yield SNAP-tag-reactive probe **11**. e) Protein-binding turn-on of **11** (2.5 μM) incubated with purified SNAP-tag protein (5 μM) in PBS for 1.5 h. f) SMLM imaging with **11** (100 nM) in live HeLa cells transfected with β-tubulin-SNAP-tag. Comparison between a superresolved and a diffraction-limited image, and localization precision of single-molecules of probe **11** in live cells. Scale bar: 20 μm (panel c) and 2 μm (panel f). FWHM = full width at half-maximum.

We next investigated whether we could induce fluorescence turn-on upon conjugation of dye **4b** to a self-labeling protein tag. Given that HaloTag has been thoroughly optimized for rhodamine-based dyes^2,37^, we chose to work with SNAP-tag^38^, thereby providing an orthogonal system that could be used in multiplexing studies with rhodamines and HaloTag. We synthesized the azide-modified SNAP-tag ligand **10** using the diazotizing reagent FSO_2_N_3_^39^ and combined it with alkyne-modified Cy5 dye **4c** in a “click” reaction to generate probe **11** (Fig. 2d and Methods). This dye displayed a 10-fold turn-on in absorbance and a 21-fold turn-on in fluorescence when incubated with purified SNAP-tag protein (Fig. 2e and Methods), demonstrating its fluorogenicity upon binding to a self-labeling tag.

Importantly, probe **11** was designed to mimic the cyclization equilibrium in HMSiR (**1**), which is a spontaneously blinking fluorophore useful for SMLM. Thus, to assess the spontaneous blinking properties of probe **11**, we expressed SNAP-tag fused to β-tubulin in HeLa cells and treated these cells with compound **11** (Methods and Supplementary Tables 2 and 3). We performed live-cell SMLM imaging in growth medium without any blinking or anti-fading agents (Methods). These experiments revealed that probe **11** indeed displays spontaneous blinking (Supporting Movie 1), allowing for the acquisition of super-resolved fluorescence images (Fig. 2f). Single-molecules of probe **11** could be localized with a precision of 18 ±6 (mean ± standard deviation) nm, and widths of about 60 nm were measured for microtubules in living cells (Fig. 2f). These results are comparable to those obtained with HMSiR (**1**)^32^.

### A fluorogenic Cy5 derivative

Next, we applied the 5-*exo-trig* ring-closing strategy to develop a no-wash, fluorogenic Cy5 derivative. We transformed the methyl ester group of **9** into *N*-methyl amide **12** and prepared the corresponding 5-*endo-trig* indoleninium **13** and Cy5 probe **14** (Fig. 3a). These alkyne-bearing derivatives were conjugated to benzyl guanine **10** to generate probes **15** and **16** (Fig. 3b). Incubation of these dyes with purified SNAP-tag protein (Methods) led to a 6-fold increase in absorbance and 19-fold increase in fluorescence for compound **15**. The larger turn-on in fluorescence indicates that binding to SNAP-tag not only induces ring-opening but also increases the quantum yield of emission of compound **15** compared to the free dye in solution (Supplementary Table 4). In comparison, compound **16** underwent only a 1.4-fold increase in absorbance and 2.5-fold increase in fluorescence upon binding to SNAPtag (Fig. 3c). The *in vitro* fluorogenicity of probe **15** is comparable to that of the popular **JF646** dye functionalized with a benzyl guanine (**JF646**-BG) for SNAP-tag labeling (Fig. 3c). This observation confirms that the 5-*exo-trig* cyclization of polymethine dyes can be reversed upon binding to protein targets, analogously to the turn-on mechanism of rhodamines.

**Fig. 3:**
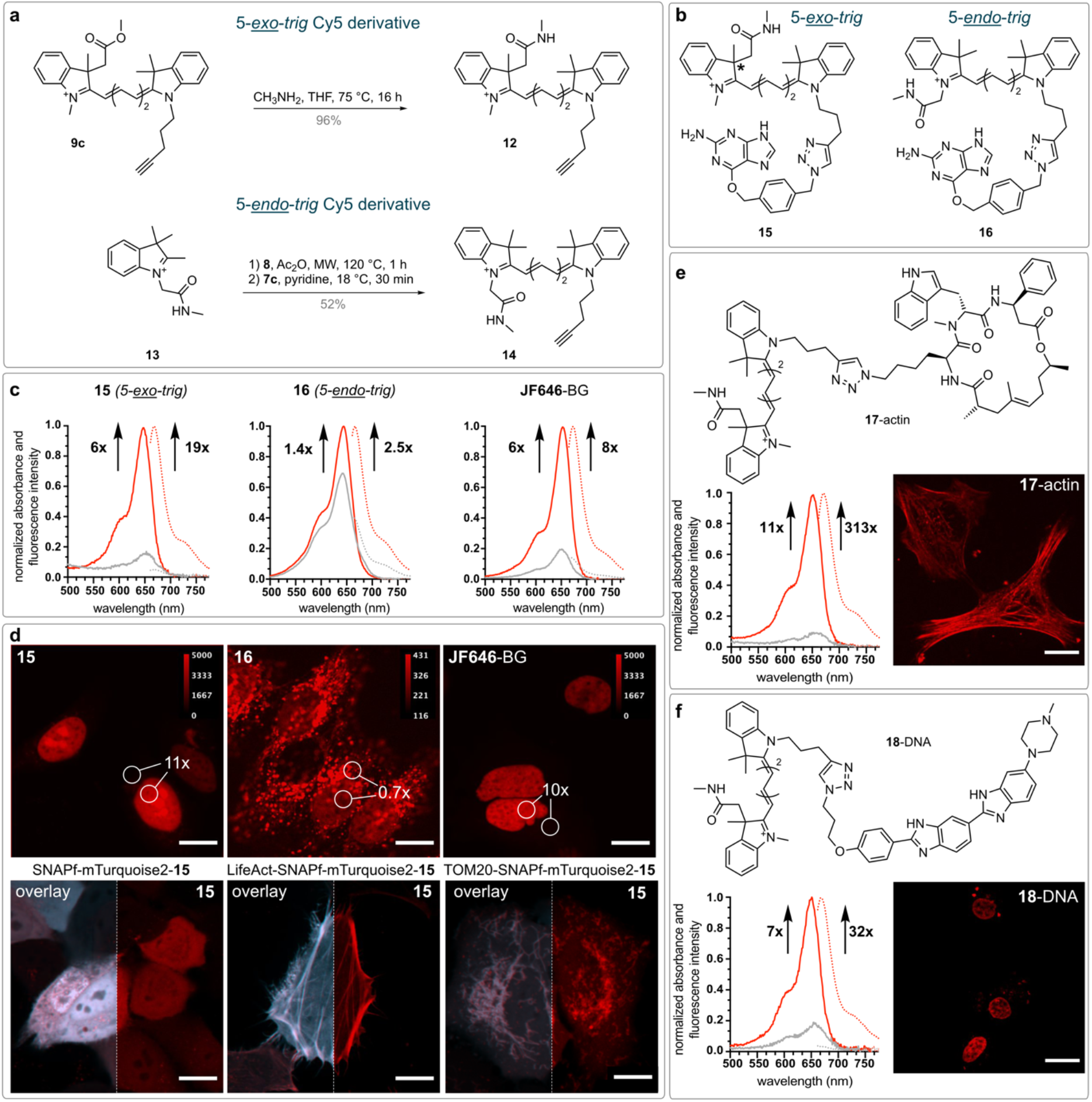
Fluorogenic Cy5 derivatives. a) Synthesis of 5-*endo-trig* and 5-*exo-trig* Cy5 derivatives with an *N*-methyl amide as a ring-closing moiety. b) Structures of compounds **15** and **16** functionalized with benzyl guanine for SNAP-tag labeling. *Chiral carbon. c) Protein-binding turn-on and no-wash imaging of live HeLa cells employing **15** (50 nM), **16** (50 nM), or **JF646**-BG (50 nM). For protein-binding studies, probes (2.5 μM) were incubated with 5 μM purified SNAP-tag protein in PBS, pH 7.4, for 1.5 h. d) No-wash, live-cell imaging of HeLa cells that were transfected with H2B-SNAPf-mTurquoise2 (upper images) or with the indicated plasmid (lower images). e) Protein-binding turn-on and no-wash live-cell imaging in HeLa cells employing **17**-actin. For protein-binding studies, the concentration of **17**-actin was 2.5 μM, for cell imaging, it was 250 nM. f) DNA-binding turn-on and live-cell imaging in HeLa cells employing **18**-DNA. For DNA-binding studies, the concentration of **18**-DNA was 1 μM, for cell imaging, it was 500 nM. Scale bars: 15 μm.

Next, we carried out no-wash, live-cell imaging to compare the performance of probes **15**, **16**, and **JF646** (Methods). We transfected HeLa cells with a plasmid encoding the fusion protein H2B-SNAPf-mTurquoise2 and incubated them with dyes **15**, **16**, or **JF646**-BG (Fig. 3d). Probe **15** exhibited bright fluorescence signal in the nucleus and only very faint unspecific background signals. This performance was comparable to that of **JF646**-BG (Fig. 3d). Probe **16**, on the other hand, displayed more than 10-fold weaker fluorescence signal, indicating that it might be less membrane-permeant (Fig. 3d). In addition, probe **16** exhibited most of its fluorescent signal in vesicles, confirming that 5-*endo-trig* Cy5 derivatives are not suitable as no-wash turn-on dyes. Finally, we tested the generality of labeling with probe **15** by transfecting HeLa cells with plasmids encoding for untargeted SNAPf-mTurquoise2 fusion protein (whole cell), LifeAct-SNAPf-mTurquoise2 (actin), or TOM20-SNAPf-mTurquoise2 (mitochondria). In all cases, labeling was specific, as judged by the excellent overlay between the signal of the reference fluorescent protein mTurquoise2 and that of **15** linked to SNAP-tag (Fig. 3d).

The 5-*exo*-*trig* ring-closure design leads to an indolenine fragment that contains a chiral carbon (Extended Data Fig. 3). To investigate the effect of stereochemistry on the ring opening of the dye upon binding to SNAP-tag, we separated the enantiomers of **6**, (+)-**6** and (−)-**6**, using a chiral stationary phase (Methods). We employed electronic circular dichroism and time-dependent DFT^40^ to assign the absolute configuration of the two enantiomers (Extended Data Fig. 3 and Methods) and found that (−)-**6** has *S* absolute configuration ((+)-(*R*)-**6** and (−)-(*S*)-**6**). We used each enantiomer separately to prepare enantiomerically pure probes (*R*)-**15** and (*S*)-**15**. We did not observe a substantial difference in fluorogenicity upon binding of each enantiomer to either purified SNAP-tag or in live-cell imaging (Extended Data Fig. 3). We hypothesize that the relatively long distance between the chiral center of the dye and the protein surface alleviates any potential enantioselective binding interactions. We argue, however, that such interactions could be leveraged to further increase the fluorogenicity of 5-*exo-trig* polymethine probes upon binding to chiral targets.

We next tested whether binding to other macromolecular targets also induced fluorescence turn-on in probe **15**. For this purpose, we prepared compounds **17**-actin and **17**-DNA. These probes were composed of a 5-*exo-trig* Cy5 derivative linked through a triazole-containing alkane to either the actin-binding cyclo-depsipeptide jasplakinolide (Fig. 3e)^41^ or to Hoechst 33342 (Fig. 3f), a dye that binds to the minor groove of double-stranded DNA (dsDNA) and has been used to target fluorogenic rhodamine dyes to the cell nucleus^42^. Probe **17**-actin showed fluorogenic behavior with an 11-fold turn-on in absorbance and 313-fold turn-on in fluorescence upon binding to actin filaments *in vitro* (Fig. 3e). The much larger fluorescence turn-on compared to absorbance increase indicates that binding to actin enhances the quantum yield of carbocyanine **17** significantly. In live-cell imaging experiments, **17**-actin labeled actin fibers selectively in live, unmodified HeLa cells. Similar results were obtained for **18**-DNA, which displayed a 7-fold increase in absorbance and 32-fold increase in fluorescence upon binding to double-stranded DNA (Fig. 3f). Live-cell imaging also confirmed specific staining of the cell nucleus (Fig. 3f). These results demonstrate that the fluorogenicity of 5-*exo-trig* Cy5 derivatives is not limited to SNAP-tag, and other macromolecules can also induce fluorescence turn-on.

### Fluorogenic Cy3 and Cy7 derivatives

Cy dyes can cover a large spectral range by varying the number of conjugated carbon atoms in between the two indoleninium moieties. Thus, we next explored whether the 5-*exo-trig* fluorogenic strategy could be extended to Cy3 and Cy7 dyes, providing fluorophores for two additional imaging channels. We suspected that, compared to Cy5, the Cy3 derivative would have a higher LUMO energy, whereas the opposite would be true for the Cy7 derivative. Thus, we expected the Cy3 dyes to have a higher tendency to adopt the open form than Cy5 dyes, whereas Cy7 dyes would tend to adopt the closed form. To balance these trends, we added a CF3 moiety on the capping indoleninium to favor the closed form of Cy3. Similarly, we replaced the nucleophilic *N*-methyl amide with an electron-deficient amide to facilitate ring-opening in our Cy7 design. From a synthetic point of view, compared to the preparation of Cy5 derivatives, we only changed the commercially available linker and adjusted the temperatures during the microwave-assisted protocol (Supplementary Information).

We prepared Cy3 derivative **19** and Cy7 derivative **20** (Fig. 4a) for SNAP-tag labeling. Both probes displayed fluorogenic behavior upon binding to purified SNAP-tag protein, with increases in absorbance of 6-fold for **19** and 11-fold for **20**, and increases in fluorescence of 28-fold for **19** and 124-fold for **20** (Fig. 4b). Similar to compound **15**, the large turn-on in fluorescence indicates that binding to SNAP-tag increases the quantum yield of emission of these probes significantly compared to the free dyes in solution (Supplementary Table 4). Next, probes **19** and **20** were utilized in no-wash live-cell imaging experiments with HeLa cells transiently transfected with the H2B-SNAPf-mTurquoise2 plasmid described before. We observed excellent co-localization of the mTurquoise2 reference signal with the yellow and near-infrared fluorescence signals of **19** and **20**, respectively, confirming the specificity of the probes (Fig. 4b). Labeling of other cellular structures was performed with similar results (Extended Data Fig. 4). Probes **15**, **19**, and **20** cover a significant portion of the visible spectrum and reach into the near-infrared region. Probes **15** and **20** display excellent brightness and good photostability when bound to SNAP-tag (Fig. 4c-d and Supplementary Table 4). Cy3 derivative **19** is not as bright or photostable, but these properties could be further tuned by introducing substituents or by making the polymethine chain more rigid^43^.

**Fig. 4:**
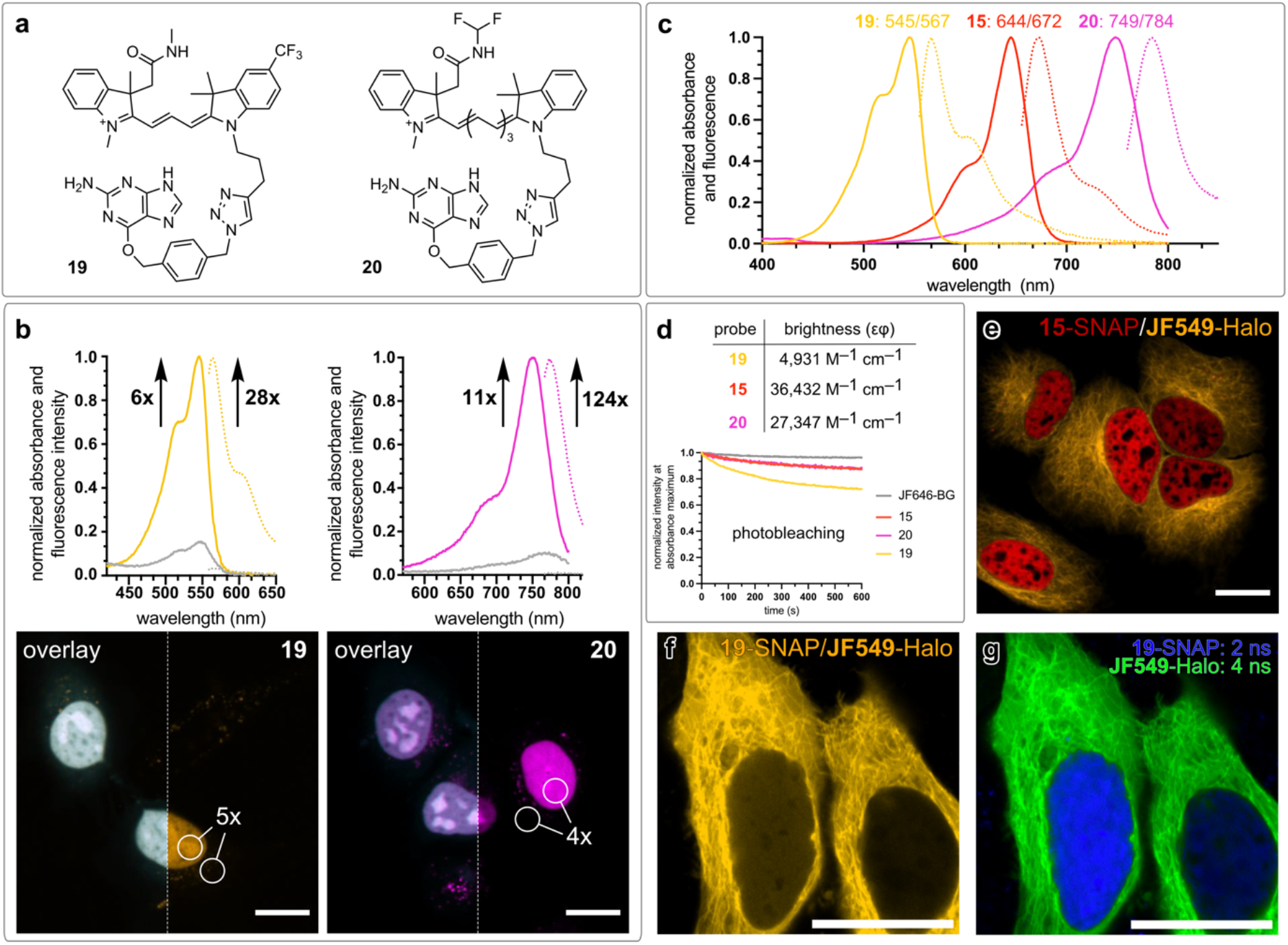
Fluorogenic Cy3 and Cy7 derivatives. a) Structures of Cy3 and Cy7 derivatives **19** and **20**. b) SNAP-tag-binding turn-on and no-wash live-cell imaging of HeLa cells transfected with H2B-SNAPf-mTurquoise2 and treated with **19** (50 nM) or **20** (250 nM). Scale bar: 15 μm. c) Normalized absorbance and fluorescence spectra of Cy3 **19**, Cy5 **15**, and Cy7 **20** measured in ethanol + 0.1% TFA. d) Brightness of SNAP-tag adducts in PBS, pH 7.4, and photobleaching curves measured for SNAPtag adducts at a concentration of 50 nM in PBS upon irradiation at the maximum excitation wavelength with monochromatic light (1.2 mW power). e) Multicolor nowash imaging in live HeLa cells co-transfected with H2B-SNAPf-mTurquoise2 and TUBB5-Halo, and incubated with **15** (50 nM) and **JF549**-Halo (50 nM). f) Total intensity measured in a cell co-transfected with H2B-SNAPf-mTurquoise2 and TUBB5-Halo and incubated with **19** (50 nM) and **JF549**-Halo (10 nM). Scale bar: 20 μm. g) Fluorescence lifetime multiplexing of the cell shown in panel (f). Scale bar: 20 μm.

Finally, we applied our probes to multiplexing experiments. First, we carried out multicolor imaging experiments using the fluorogenic Cy5 dye **15** in combination with the widely used **JF549**-Halo dye. We confirmed the orthogonality of our fluorogenic probe with rhodamine-HaloTag conjugates by imaging the **15**/**JF549**-Halo pair in live HeLa cells co-transfected with H2B-SNAPf-mTurquoise2 and TUBB5-HaloTag (Fig. 4e). Next, we explored whether the fluorescence signals of the Cy3 derivative **19** and **JF549**-Halo, which have nearly identical excitation and emission wavelengths, could be separated by their excited-state lifetimes. We co-transfected HeLa cells with H2B-SNAPf-mTurquoise2 and TUBB5-HaloTag and performed fluorescence lifetime imaging (FLIM). Whereas the signals could not be separated by their wavelength (Fig. 4f), they could be easily unmixed by their average excited-state lifetimes (2 ns for **19** and 4 ns for **JF549**-Halo, Fig. 4g) using phasor plot analysis (Methods).

## Conclusions

We presented a general strategy to impart fluorogenicity to polymethine dyes *via* a 5-*exo-trig* ring-closure approach. These dyes, regardless of their excitation wavelength, can be readily synthesized in two high-yielding steps and are easy to derivatize by varying the indoleninium building blocks or the ring-closing moiety. We have illustrated the potential of our fluorogenic polymethine scaffold by generating the first spontaneously blinking Cy5 dye, fluorogenic Cy3 and Cy5 dyes, and a bright and photostable near-infrared fluorogenic Cy7. Cy7 derivative probe **20** is particularly interesting because of its high brightness and long emission wavelength (εϕ = 27,300 M^-1^ cm^-1^ and λ_em_ = 784 nm), in particular in comparison with fluorescent proteins in the same spectral range (e. g., miRFP718nano, εϕ = 4,500 M^-1^ cm^-1^ and λ_em_ = 718 nm^44^).

We demonstrated the versatility of our turn-on strategy by utilizing the self-labeling protein tag SNAP-tag, as well as jasplakinolide or Hoechst 33342 dye, to drive fluorescence turn-on. Furthermore, we showed that these probes can be used for no-wash confocal live-cell microscopy, SMLM, and in combination with the widely used rhodamine-HaloTag conjugates in multicolor and multiplexed lifetime imaging experiments.

Given the high modularity of polymethine dyes, the spectral range can be further extended into the green (e. g., Cy1 dyes^45^) as well as into the shortwave infrared (e. g., nonamethine dyes^46^) wavelengths. Furthermore, the photophysical properties and fluorogenicity of polymethine dyes could be further tuned by varying the substituents on the indolenines or the linker^13^. We envision that this simple, yet general, method will be used to develop improved fluorogenic probes, facilitating new bioimaging experiments.

## Methods

### General remarks

All reagents were purchased from commercial sources and used as received. Anhydrous solvents were procured from Acros Organics and used as received. NMR spectra were acquired on a Bruker 400 or 600 instrument. ^1^H NMR chemical shifts are reported in ppm relative to SiMe_4_ (δ = 0) and were referenced internally with respect to residual protons in the solvent (δ = 7.26 for CDCl_3_, δ = 1.94 for CD_3_CN, δ = 3.31 for CD_3_OD and δ = 2.50 for (CD_3_)_2_SO). Coupling constants are reported in Hz. ^13^C NMR chemical shifts are reported in ppm relative to SiMe_4_ (δ = 0) and were referenced internally with respect to solvent signal (δ = 77.16 for CDCl_3_, δ = 1.32 for CD_3_CN, δ = 49.00 for CD_3_OD and δ = 39.52 for (CD_3_)_2_SO). High-resolution mass spectra (HRMS) were acquired on a *timsTOF Pro* TIMS-QTOF LC/MS spectrometer (Bruker Daltonics GmbH, Bremen, Germany) by using electrospray ionization (ESI). IUPAC names of all compounds are provided and were determined using CS ChemBioDrawUltra 16.0.

### Computational Modeling

All calculations were carried out using Gaussian 09 at the B3LYP/DGTZVP level of theory as well as at the M06-2X/DGTZVP level. An implicit solvation model (IEFPCM) was used to simulate the effect of an aqueous environment. All stationary states were characterized by harmonic analysis at the same level of theory. All minima displayed zero imaginary frequencies and all transition states gave one imaginary frequency along the C–O bond elongation coordinate. Energies were corrected by zero-point energy.

### Optical Spectroscopy

Stock solutions were prepared in DMSO (spectrophotometric grade >99.9%) at concentrations of 5 mM, 1 mM, and 50 μM and stored at –20 °C. Spectroscopic measurements were conducted in phosphate-buffered saline (PBS). UV-Vis spectra were acquired using a Multiskan SkyHigh Microplate Spectrophotometer (ThermoFisher Scientific) and quartz cuvettes from ThorLabs (10 mm path length).

Buffered aqueous solutions in the pH range of 2 to 8 were prepared by mixing citric acid (0.1 M) and sodium dihydrogen phosphate NaH_2_PO_4_ (0.2 M) in ultrapure water. Buffered aqueous solutions in the pH range of 9 to 11 were prepared by mixing sodium bicarbonate (0.1 M) and sodium carbonate (0.1 M) in ultrapure water. Buffered aqueous solutions in the pH range of 12 to 13 were prepared by mixing potassium chloride KCl (0.2 M) and sodium hydroxide (0.2 M). 5 μM solutions of the dyes in the buffered aqueous solutions were prepared and the absorbance spectra were recorded after 1.5 h in triplicates using 96-well plates (Corning) on the MultiSkan SkyHigh microplate reader. The obtained UV-Vis spectra were background corrected and the absorbance maxima of each pH value were normalized. Normalized absorbance values were plotted against the pH value and fitted (non-linear fit, sigmoidal, 4PL) using Prism 8.4.

Extinction coefficients were obtained by measuring the absorbance spectra at concentrations of 15 μM, 12.5 μM, 10 μM, 7.5 μM, 5 μM, 2.5 and 1.25 μM. The absorbance maxima were plotted against the corresponding concentration and fitted (simple linear regression) using Prism 8.4.

Fluorescence spectra were acquired using an FS5 Spectrofluorometer (Edinburgh Instruments) equipped with an SC-25 cuvette holder or SC-40 plate reader. Absolute fluorescence quantum yields were determined using 1 μM solutions employing an integrating sphere (SC-30, Edinburgh instruments). All spectroscopic measurements were carried out in triplicates and at room temperature.

Photobleaching experiments were carried out using a FS5 Spectrofluorometer equipped with an SC-25 cuvette holder. Protein-adducts were measured at a 50 nM concentration and the emission slit width was adjusted individually for each dye to achieve a power of 1.2 mW (547 nm: 12 nm, 646 nm and 654 nm: 22.5 nm, 751 nm: 29.9 nm). The emission slit width was set to 0.75 nm and the fluorescence was measured every second for ten minutes while keeping the shutter always open. The power was measured at the cuvette holder using a PMD100D compact power and energy meter console equipped with a S120VC standard photodiode power sensor (UV-Extended Si, 200 - 1100 nm, 50 mW, Thorlabs GmbH).

### Cloning

All plasmids were cloned by Gibson assembly. DNA encoding SNAPf or H2B-SNAPf was amplified from pSNAPf-H2B control plasmid (Addgene #101124) to generate an insert. To generate a backbone containing an organelle-specific targeting group and a fluorescent protein, the plasmids pmTurquoise2-ER (Addgene #36204) and pmTurquoise2-Golgi (Addgene #36205) were used. Primers for amplification (minimum 15 overlapping base pairs) were generated using SnapGene^®^ and modified manually to minimize secondary structures, self-dimers, and repeated motifs. The designed primers were supplied by Microsynth AG (Switzerland) and are reported in Supplementary Tables 2 and 3. Vector and insert fragments were amplified by PCR using Phusion High-Fidelity PCR Master Mix with HF buffer from New England Biolabs (NEB). The fragments were analyzed by agarose gel electrophoresis and template DNA was digested using DpnI. PCR fragments were purified using a QIAquick PCR purification kit (Qiagen) according to the manufacturer’s instructions. The Gibson assembly reaction was performed using the Gibson Assembly Master Mix (NEB) according to the manufacturer’s protocol. NEB DH5α competent *E. coli* cells were transformed with the assembly product by heat shock following the manufacturer’s instructions, streaked onto lysogeny broth (LB) agar plates containing kanamycin (50 μg mL^-1^), and incubated at 37 °C for 24 h. Single colonies were selected and grown in LB liquid medium containing 50 μg mL^-1^ kanamycin at 37 °C for 16 h. Plasmid DNA was isolated using the QIAprep spin miniprep kit (Qiagen) according to the manufacturer’s instructions. The correct sequence of the gene of interest (GOI) was verified by the Sanger sequencing service of Microsynth using the Standard sequencing primers CMV-F, SV40pA-R, and EGFP-C. All plasmids produced in this paper are available on Addgene (#197494–98).

### Expression and purification of fSNAP protein

pET24b-6His-fSNAP (Addgene #106999) was transformed into *E. coli* strain Rosetta2(DE3) and streaked on an agar plate with kanamycin and chloramphenicol resistance. A single colony was picked, inoculated in 80 mL LB cultures containing 50 μg mL^-1^ kanamycin and 25 μg mL^-1^ chloramphenicol, and grown at 37 °C for 24 h. 40 mL of the starter culture were inoculated into 4 L of medium and shaken at 37 °C to an optical density at 600 nm (OD_600_) of 0.8. Expression was induced by the addition of 0.5 mM isopropyl β-D-thiogalactopyranoside (IPTG), and cells were grown at 18 °C for 16 h. Cells were harvested by centrifugation, resuspended in HEPES buffer (300 mM NaCl, 20 mM HEPES, pH = 7.5), and supplemented with turbonuclease (20 μL) and a protease inhibitor cocktail tablet (Roche). Cells were lysed by sonication (70% power, 10-sec pulse/10-sec pulse off for 2 min 30 sec) and the lysate was cleared by centrifugation. The protein was purified by Ni-His-affinity column chromatography and the fractions were analyzed by SDS-PAGE. The fractions containing the protein were concentrated using Amicon Ultra-4 and Ultra-15 centrifugal filter units (MWCO 10 kDa) and further purified by size-exclusion chromatography. The correct size and purity of the protein were verified by SDS-PAGE analysis. Purified SNAP protein was stored in aliquots in a buffer containing 150 mM NaCl, 20 mM HEPES, pH 7.4, 1 mM DTT at −80 °C.

### Protein-binding turn-on assays

Prior to biological assays with purified SNAP-tag protein, the buffer was exchanged to PBS using Zebra™ Spin Desalting Columns (Thermo Scientific Inc.) according to the manufacturer’s protocol. SNAP-dyes (1 mM in DMSO) were either added to PBS alone or to a 5 μM SNAP-protein solution in PBS. The final concentration of SNAP-dyes was 2.5 μM and the resulting mixtures were incubated at 37 °C for 1.5 h.

The turn-on assay with actin was carried out according to a published procedure^47^. Actin ligand **17**-actin (1 mM in DMSO) was added to either supplemented actin buffer alone or in the presence of 0.4 mg mL^-1^ G-actin (cat. AKL99, Cytoskeleton Inc.). The final ligand concentration was 2.5 μM. The supplemented actin buffer contained 5 mM Tris-HCl pH 8.0, 0.2 mM CaCl_2_ (from General Actin Buffer, cat. BSA01, Cytoskeleton Inc.), 50 mM KCl, 2 mM MgCl_2_, 5 mM guanidine carbonate and 1.2 mM ATP (from Actin Polymerization Buffer, cat. BSA02, and ATP, cat. BSA04, Cytoskeleton Inc.). Samples were incubated at 37 °C for 2 h.

The DNA-binding assay was carried out according to a published procedure^42^. The hairpin-forming 28-bp DNA oligonucleotide was purchased from Microsynth (5’-CGCGAATTCGCGTTTTCGCGAATTCGCG-3’) and dissolved in Tris-buffered saline (TBS, 50 mM Tris-HCl, 150 mM NaCl, pH 7.4) at 1 mM concentration. The folding of the DNA into the secondary hairpin structure was achieved by heating the DNA at 75 °C for 2 min followed by slowly cooling to 25 °C. Nucleus ligand **18**-DNA (1 mM in DMSO) was incubated with either TBS (50 mM Tris-HCl, 150 mM NaCl, pH 7.4) alone or in the presence of 50 μM hairpin DNA at room temperature for 1 h. The final ligand concentration was 1 μM.

Absorbance spectra were recorded in 96-well plates (Corning) on the Multiskan SkyHigh microplate reader and fluorescence measurements were carried out on the FS5 Spectrofluorometer (Edinburgh Instruments) equipped with a SC-40 plate holder. All spectroscopic measurements were carried out in triplicates and at room temperature.

### Cell Culture and Fluorescence Imaging

HeLa cells were grown in Dulbecco’s Modified Eagle Medium (DMEM) supplemented with fetal bovine serum (FBS, 10%) and penicillin-streptomycin (1%) at 37 °C in a 95% humidity atmosphere under a 5% CO2 environment. The cells were grown to 90% confluency before seeding at a density of 15-20 000 cells mL^-1^ onto Ibidi μ-Slide 8-well glass-bottom plates 48 h before the imaging experiment. Cells were transfected with the plasmids H2B-SNAPf-mTurquoise2, LifeAct-SNAPf-mTurquoise2, TOM20-SNAPf-mTurquoise2, SNAPf-mTurquoise2-KDEL or SNAPf-mTurquoise2 using jetPrime^®^ according to the manufacturer’s protocol 24 h prior to imaging. Cells were incubated with the respective probes in FluoroBrite DMEM for 1.5 h and imaged directly.

### Confocal microscopy

Confocal imaging was performed with a Nikon W1 spinning disk microscope equipped with a CMOS camera (Photometrix). Brightfield imaging was performed with a white LED. Laser lines and filters were set up for the appropriate channel as described in Supplementary Table 5.

Images were collected using a CFI Plan Apochromat Lambda D oil immersion objective (60x, NA = 1.4). Channels were imaged sequentially. The microscope was operated using the NIS elements software. Imaging experiments were performed at 37 °C in a 5% CO_2_ environment. Images were analyzed by Fiji/ImageJ.

### Determination of the absolute configuration of (+)-6 and (−)-6

Circular dichroism (CD) spectra were measured with a Chirascan™ V100 CD spectrometer (Applied Photophysics) and operated using the software Chirascan. For CD measurements, 60 μM solutions of (+)-**6** and (–)-**6** in PBS were used. Optical rotation measurements were carried out with a Jasco P-2000 polarimeter and operated using the Spectra Manager software.

For TD-DFT calculations, we first performed a systematic conformational search at the B3LYP/DGTZVP level of theory with an implicit solvation model (IEFPCM) by varying all rotatable bonds in 60° steps. Next, we applied the Boltzmann distribution to the set of low-energy minima obtained by using the free energy differences and considered the structures above the 0.1% population threshold for the TD-DFT calculation. TD-DFT calculations were carried out at the CAM-B3LYP/DGTZVP level of theory with IEFPCM as the solvation model.

### Fluorescence lifetime imaging (FLIM)

FLIM was carried out on a Leica SP8 inverse FALCON confocal laser scanning microscope and operated using the Leica LAS X Navigator. Images were collected using an HC PL APO corr CS2 oil objective (63x, NA = 2.4) and the RHOD fluorescence filter set (excitation 546/10, emission 585/40). Phasor analysis of the FLIM data was performed with the Leica LAS X Phasor License.

### SMLM

HeLa cells were transfected with β-tubulin-SNAP plasmid using jetPrime^®^ according to the manufacturer’s protocol 48 h before imaging. Cells were incubated with probe **11** (100 nM) for 14 h, then detached and transferred to a new Ibidi slide. Cells were left for 10 h to adhere to the glass surface, and the medium was exchanged to FluoroBrite prior to imaging. Single-molecule imaging was performed on a Ti2 eclipse inverted microscope (Nikon Ltd.) equipped with a water-cooled iXon 888 Ultra EMCCD camera (Andor) at 37 °C in a 5% CO_2_ environment. Acquisitions were carried out at 638 nm (90 mW, 30 ms) and a single band 708/75 emission filter was used. 2000 frames were acquired. Acquisitions were collected using a NIKON 100x TIRF Apo Plan SR oil objective (NA = 1.49). All laser and camera shutters were controlled by a NIDAQ oscilloscope (National Instruments) controller unit. The single-molecule signal was fitted with 2D Gaussian point spread functions using ThunderSTORM. The localization threshold was set to 1.1*std(Wave.F1).

## Data and materials availability

All data supporting this paper are available through Zenodo (DOI: 10.5281/zenodo.7588846). Plasmids generated in this work are available through Addgene (#197494-98).

## Acknowledgments

This work was supported by EPFL (SViPhD internal grant) and by the European Research Council (ERC Starting Grant HDPROBES, 801572). We thank Sarah Hübner for assistance with FLIM, Léa Blatti for the synthesis of some indoleninium building blocks, Karl Gademann for access to a polarimeter, and Pierre Gönczy and Luc Reymond for valuable discussions. This work made use of infrastructure services provided by SCITAS, the Scientific IT and Application Support of EPFL, and S^3^IT, the Service and Support for Science IT at the University of Zurich. Samples of jasplakinolide and Hoechst 33342 ligands were kindly donated by Spirochrome AG (https://spirochrome.com/). FLIM was carried out at the Center for Microscopy and Image Analysis of the University of Zurich.

## Author contributions

A. M. and P. R.-F. conceived the method. A. M. carried out all experiments. A. M. and P. R.-F. carried out computational modeling and analyzed the results. A. M. and P. R.-F. wrote the manuscript. P. R.-F. acquired funds and supervised the project.

## Competing Interests

EPFL and University of Zurich jointly filed a patent application (EP23153834.9) protecting the invention disclosed in this paper.

## Extended Data

**Extended Data Fig. 1:**
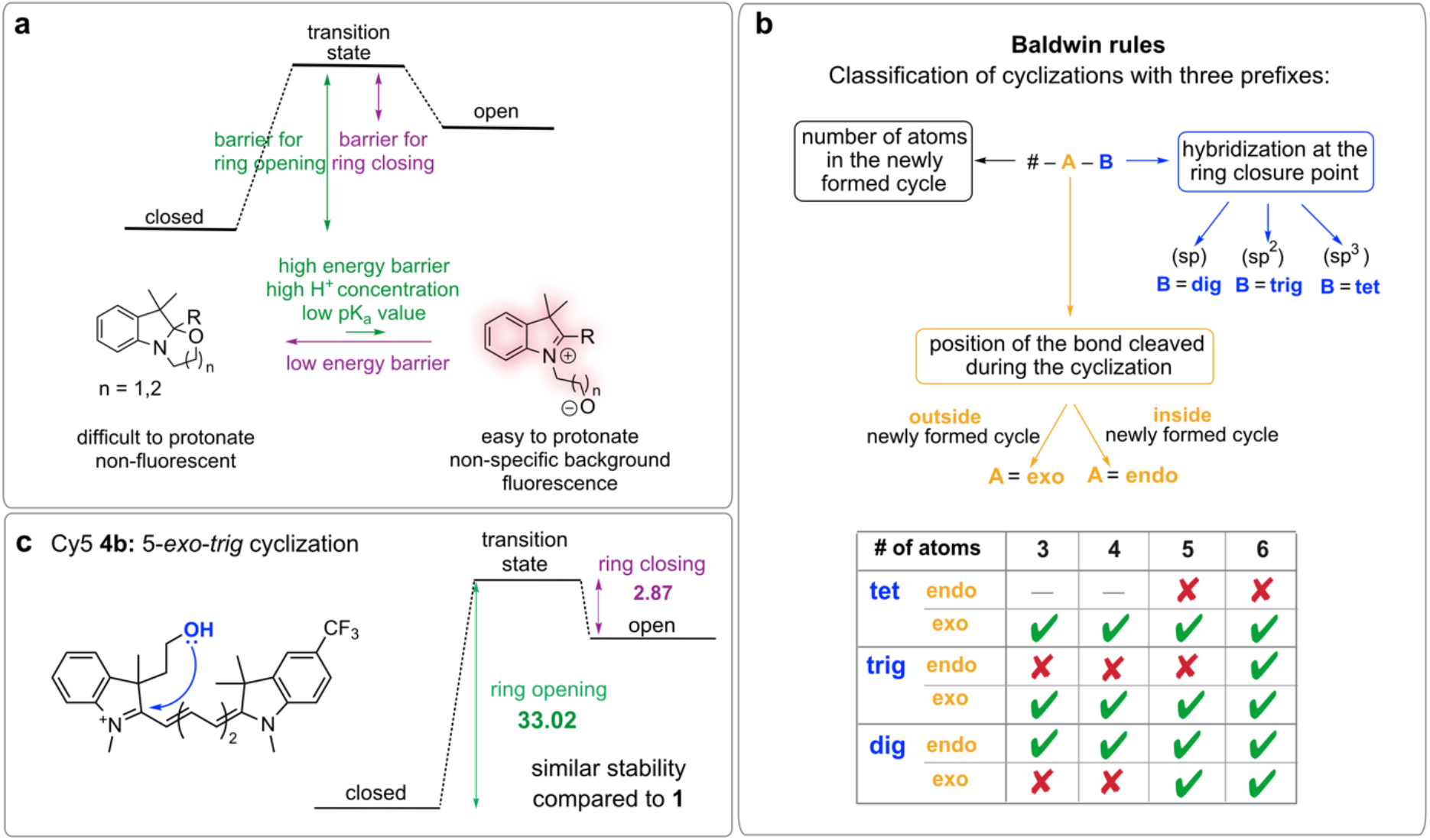
Probe design and Baldwin rules. a) Cyclization energy landscape that favors fluorogenicity: A large barrier of ring opening and a small barrier of ring closure. b) Explanation of the Baldwin rules. c) Structure and calculated energy of ring opening and closing of Cy5 derivatives **4b**. Calculations were carried out at the B3LYP/DGTZVP level of theory and energy values are reported in kcal mol^-1^.

**Extended Data Fig. 2:**
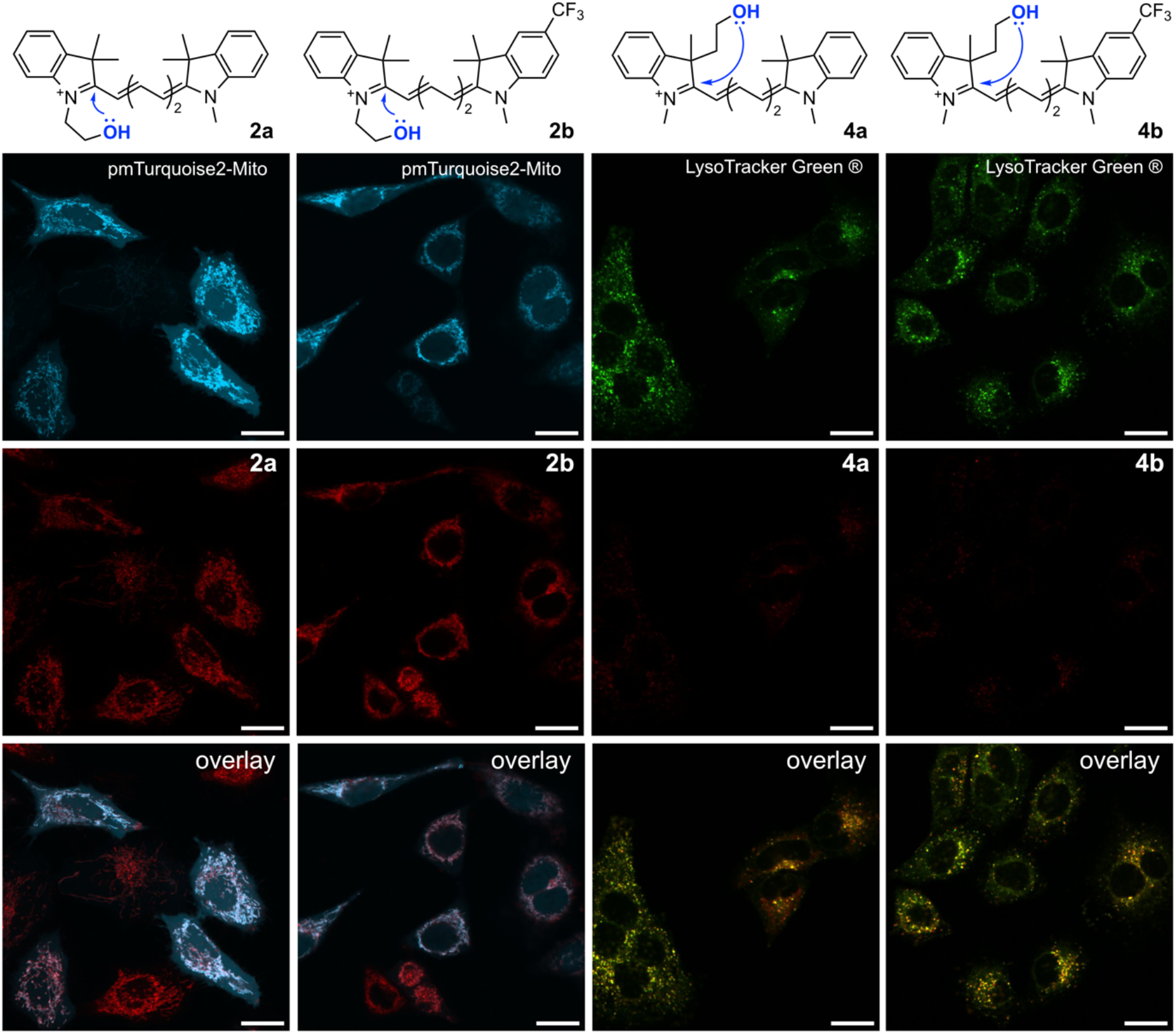
Subcellular localization of untargeted dyes 2a, 2b, 4a, and 4b. For 5-*endo-trig* probes **2a** and **2b**, HeLa cells were transfected with a plasmid encoding for COX8A-mTurquoise2 and incubated with 250 nM **2a** or **2b** for 2 h. For 5-*exo-trig* probes **4a** and **4b**, HeLa cells were incubated with 250 nM **4a** or **4b** for 2 h. LysoTracker Green^®^ (100 nM) was added 15 min prior to imaging.

**Extended Data Fig. 3:**
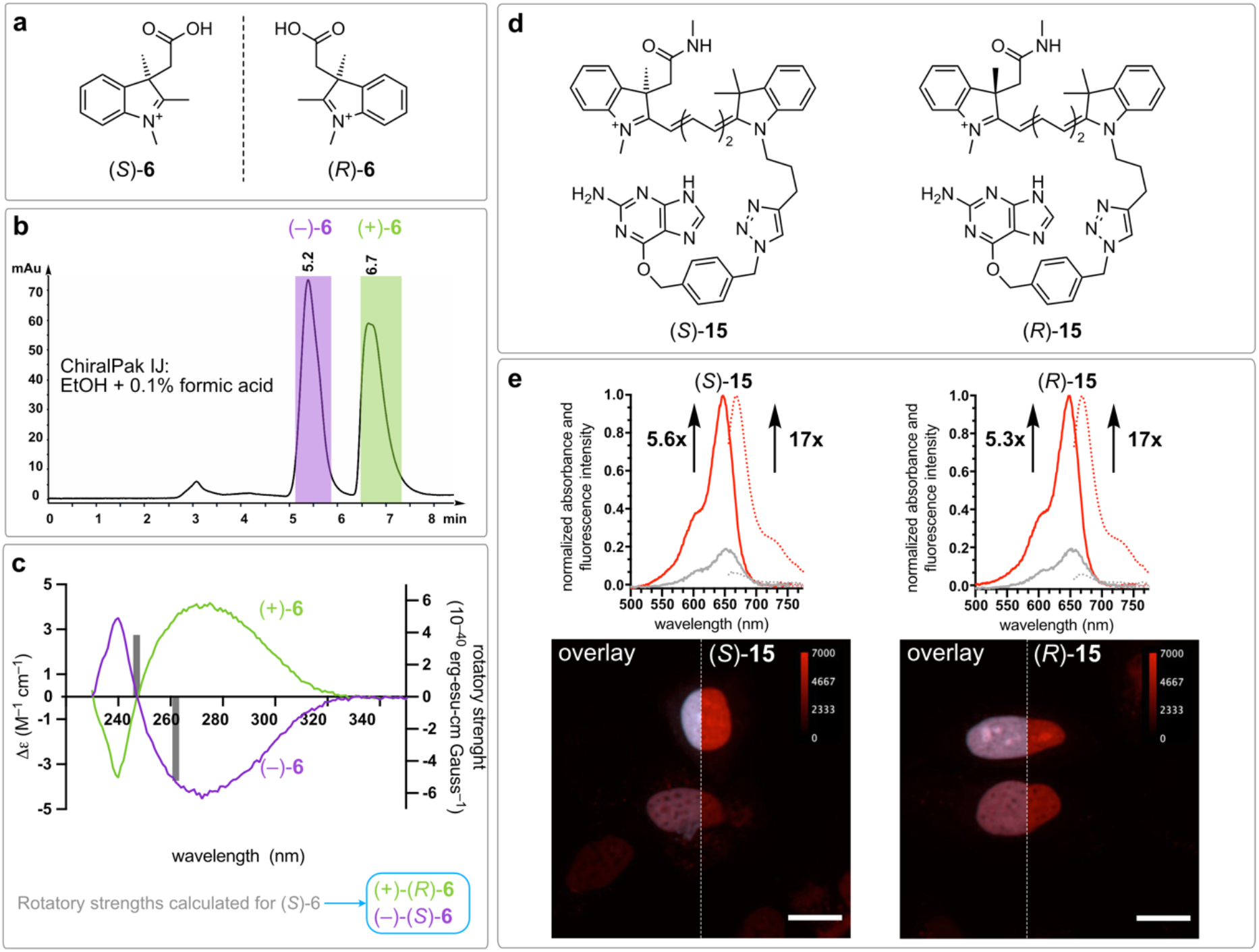
Separation and evaluation of the enantiomers (*S*)-15 and (*R*)-15. a) Structure of the enantiomers (*S*)-**6** and (*R*)-**6**. b) Separation of enantiomers (*S*)-**6** and (*R*)-**6** by HPLC under the conditions indicated. c) Determination of the absolute configuration of the enantiomers of **6** by electronic circular dichroism and comparison with the calculated rotatory strengths (grey bars) of *(S)*-**6** using TD-DFT (CAM-B3LYP/DGTZVP). d) Structures of the enantiomeric dyes. (*S*)-**15** and (*R*)-**15**. e) Protein-binding turn-on of probes (*S*)-**15** and (*R*)-**15** (2.5 μM) incubated with 5 μM purified SNAP-tag protein in PBS, pH 7.4, for 1.5 h. No-wash confocal microscopy of live HeLa cells transfected with H2B-SNAPf-mTurq2 and incubated with (*S*)-**15** (50 nM) and (*R*)-**15** (50 nM). Scale bars: 15 μm.

**Extended Data Fig. 4:**
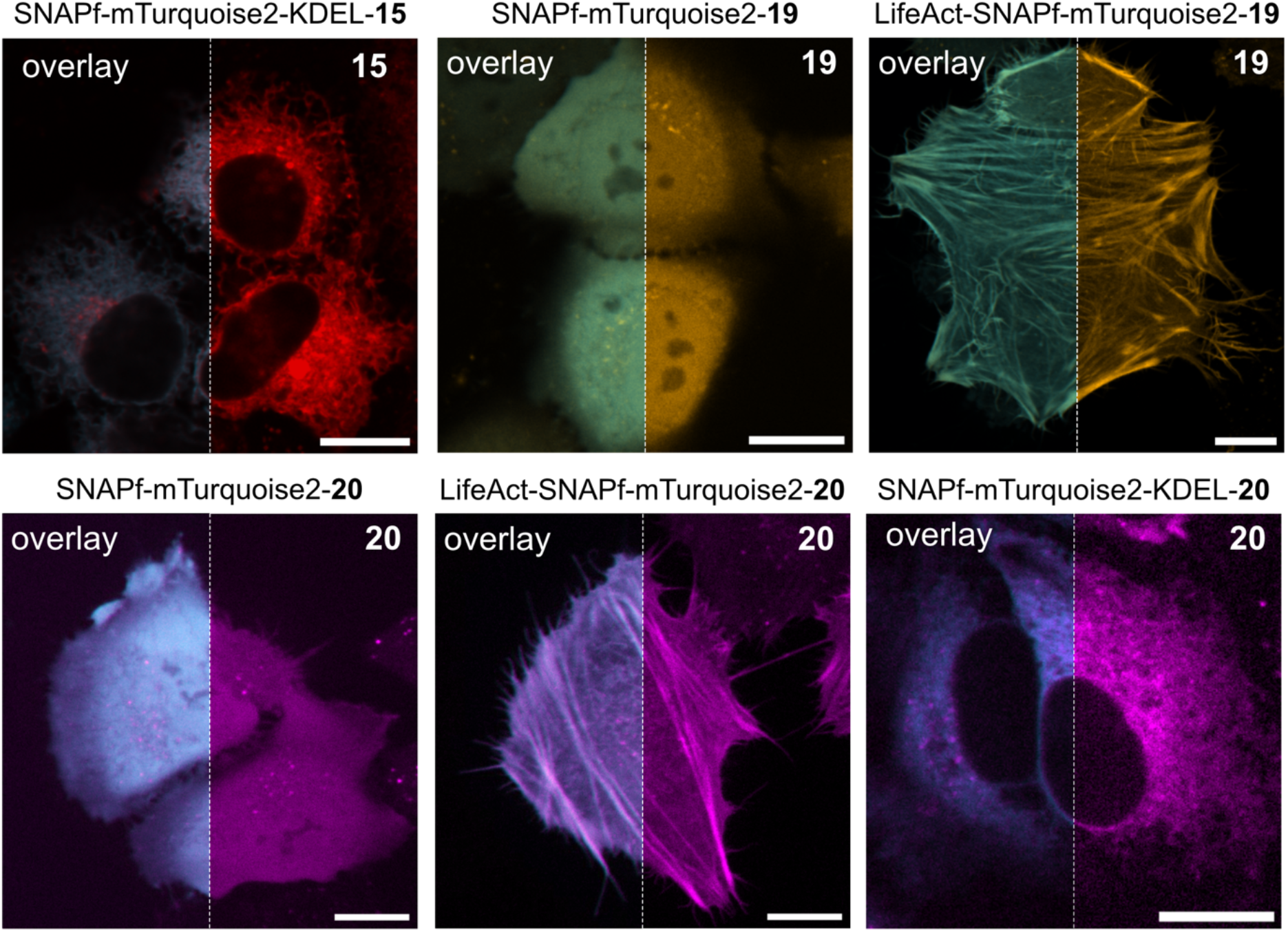
Additional cellular targeting of probes 15, 19 and 20. No-wash confocal microscopy of live HeLa cells transfected with SNAPf-mTurquoise2-KDEL, SNAPf-mTurquoise2 or LifeAct-SNAPf-mTurquoise2 and incubated with **15** (50 nM), **19** (50 nM) or **20** (250 nM). The overlay shows the signal of the reference fluorescent protein mTurquoise2 and that of the indicated compound linked to SNAP-tag. Scale bars: 15 μm.

## Notes

https://doi.org/10.5281/zenodo.7588846

