## Supporting Information for "A general strategy to develop fluorogenic polymethine dyes for bioimaging"

#### Supplementary Tables

**Supplementary Table 1.** Calculated energies of the open and closed form of **1**, **2**, **3, 4a**, **4b** and of the transition state of ring closure using DFT theory, B3LYP or M06-2X functional and the DGTZVP basis set. Energy values are reported in kcal $mol^{-1}$.

| Probe | closed form | open form | transition state |
|---|---|---|---|
| 1 (B3LYP) | -930591.73 | -930566.39 | -930563.39 |
| 2 (B3LYP) | -797468.7184 | -797455.0657 | -797449.7312 |
| 3 (B3LYP) | -822124.75 | -822108.05 | -822102.69 |
| 4a (B3LYP) | -797472.89 | -797446.15 | -797441.7 |
| 4b (B3LYP) | -1009053.449 | -1009023.302 | -1009020.434 |
| 1 (M06-2X) | -930256.9573 | -930219.9555 | -930216.4001 |
| 2 (M06-2X) | -797134.25 | -797104.38 | -797103.34 |
| 3 (M06-2X) | -821779.50 | -821747.81 | -821745.36* |
| 4a (M06-2X) | -797137.69 | -797096.93 | -797094.28 |
| 4b (M06-2X) | -1008650.22 | -1008608.32 | -1008605.57 |

*For the calculation of this transition state, the following gaussian command was used: opt=(calcfc,ts,noeigentest)

**Supplementary Table 2.** Plasmid sources.

| Plasmid | Source | Vector precursor | Insert precursor |
|---|---|---|---|
| pET24b-6His-fSNAP | Addgene #106999 | | |
| pSNAPf-H2B | Addgene #101124 | | |
| pEBTet-β-tubulin_SNAP | Addgene #136831 | | |
| pmTurquoise2-Golgi | Addgene #36205 | | |
| pmTurquoise2-ER | Addgene #36204 | | |
| pmTurquoise2-Mito | Addgene #36208 | | |
| LifeAct-Halo-T2A-eGFP | Addgene #135445 | | |
| TOM20-Halo-T2A-eGFP | Addgene #135443 | | |
| TUBB5-Halo | Addgene #64691 | | |
| SNAPf-mTurquoise2 | Gibson assembly[1] | pmTurquoise2-Golgi | pSNAPf-H2B |
| H2B-SNAPf-mTurquoise2 | Gibson assembly[1] | pmTurquoise2-Golgi | pSNAPf-H2B |
| SNAPf-mTurquoise2-KDEL | Gibson assembly[1] | pmTurquoise2-ER | pSNAPf-H2B |
| LifeAct-SNAPf-mTurquoise2 | Gibson assembly[1] | H2B-SNAPf-mTurquoise2 | LifeAct-HT7-T2A-eGFP |
| TOM20-SNAPf-mTurquoise2 | Gibson assembly[1] | H2B-SNAPf-mTurquoise2 | TOM20-HT7-T2A-eGFP |

**Supplementary Table 3.** List of primers for Gibson Assembly.

| Plasmid | **backbone forward primer** | **backbone reverse primer** | **fragment forward primer** | **fragment reverse primer** |
|---|---|---|---|---|
| SNAPf-mTurquoise2 | GAGGT TAATG TGAGC AAGGGC | TTTGT CCATT GA GTC CGGTA GCG | CGGAC TCAAT GGACA AAGAC TGC | CCTTG CTCAC ATTAA CCTCG AG |
| H2B-SNAPf-mTurquoise2 | GAGGT TAATG TGAGC AAGGGC | TCTGG CATTG AGTCC GGTAGC | CGGAC TCAAT GCCAG AGCCA | CCTTG CTCAC A TTAA CCTCGAG |
| SNAPf-mTurquoise2-KDEL | GAGGT TAATG TGAGC AAGGGC | GTCTT TGTCC ATGGT GGCGAC | CCACC ATGGA CAAAG ACTGC | CCTTG CTCAC ATTAA CCTCG AG |
| LifeAct-SNAPf-mTurquoise2 | CGGTA CTGCG CCAGC ATTT | CCACG CCCAT TGAGT CCGG | CTCAA TGGGC GTGGC CGAC | GCTGG CGCAG TACCG ATTTCG |
| TOM20-SNAPf-mTurquoise2 | TGTGG AAGCG CCAGC ATTT | CCCAC CATTG AGTCC GGTAG | GGACT CAATG GTGGG TCGG | GCTGG CGCTT CCACA TCATCT |

**Supplementary Table 4.** Characterization of SNAP-conjugated fluorogenic dyes.

| SNAP-dye | $\lambda_{ex}/\lambda_{em}$ | $\varepsilon$ [$M^{-1}cm^{-1}$] | $\phi$ |
|---|---|---|---|
| Cy3-SNAP **19** | 545/567 | 93,550[a]/30,820[b] | 0.04[a]/0.16[b] |
| Cy5-SNAP **15** | 644/672 | 195,500[a]/79,200[b] | 0.28[a]/0.46[b] |
| Cy7-SNAP **20** | 749/784 | 233,500[a]/82,870[b] | 0.16[a]/0.33[b] |

a: ethanol + 0.1% TFA, b: bound to SNAP-tag protein in PBS

**Supplementary Table 5.** Microscope settings for imaging channels.

| Channel | $\lambda_{ex}$ | Emission filter | Laser intensity | Exposure time |
|---|---|---|---|---|
| mTurquoise2 | 445 nm | 472/30 | 30 mW | 300 ms |
| Cy3, JF549 | 561 nm | 600/52 | 23 mW | 300 ms |
| Cy5, JF646 | 638 nm | 708/75 | 25 mW | 400 ms |
| Cy7 | 638 nm | 647LP | 25 mW | 400 ms |

#### Synthetic procedures

##### Synthesis of building blocks

Building blocks **5**, **7a**, **7b**, **8** and **13** were prepared according to previously reported methods.[2–4]

##### 3-(Carboxymethyl)-1,2,3-trimethyl-3*H*-indol-1-ium bromide[2] (6)

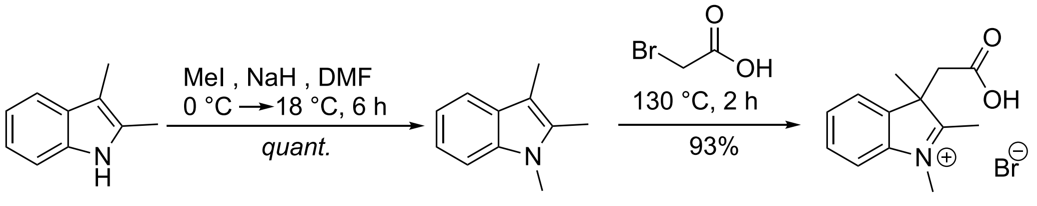

**6**

Sodium hydride (60% dispersion in mineral oil, 7.3 g, 183 mmol, 2.6 equiv.) was added to a flame-dried round-bottom flask, cooled to 0 °C and carefully dissolved in DMF (50 mL). 2,3-dimethylindole (10 g, 68.9 mmol, 1.0 equiv.) was added portion-wise at 0 °C followed by the addition of iodomethane (5.6 mL, 89.5 mmol, 1.3 equiv.). After stirring at 0 °C for 1 h, the suspension was warmed to room temperature and stirred for 4 h. The mixture was cooled to 0 °C and cold water (25 mL) was added dropwise. The mixture was extracted twice with ethyl acetate and the organic phase was washed with brine, dried over $Na_2SO_4$ and filtered. Solvents were removed under reduced pressure to afford 1,2,3-trimethyl-1*H*-indole as a dark orange oil, which was used directly without further purification.

Bromoacetic acid (28.8 g, 207 mmol, 3.0 equiv.) was slowly melted in a flame-dried flask (45-50 °C) under argon atmosphere. 1,2,3-trimethyl-1*H*-indole (11 g, 69.1 mmol, 1.0 equiv.) was added and the mixture was heated to 130 °C for 2 h. The mixture was slowly cooled to 18 °C and the residue was washed twice with diethyl ether and $CH_2Cl_2$. Acetonitrile was added and the suspension was left in the fridge for 1 h. The beige precipitate was collected by filtration and washed with diethyl ether to afford the product as a beige solid (19.1 g, 93%). The racemic mixture was then separated on a ChiralPak IJ column using 100% EtOH + 0.1% FA as mobile phase.

(*R*)-(+)-6: $[\alpha]_{589}^{296K}$ = +74° ± 2°, (c = 6.0 mg $mL^{-1}$, $CH_3OH$).
(*S*)-(–)-6: $[\alpha]_{589}^{297K}$ = –64° ± 3°, (c = 6.0 mg $mL^{-1}$, $CH_3OH$).
$^1$H NMR (400 MHz, $CD_3OD$): δ = 7.83 (dd, *J* = 6.6, 2.5 Hz, 1H), 7.79 – 7.77 (m, 1H), 7.68 – 7.61 (m, 2H), 4.07 (s, 3H), 3.58 (d, *J* = 17.5 Hz, 1H), 3.50 (d, *J* = 17.5 Hz, 1H), 1.56 (s, 3H) ppm.
$^{13}$C{$^1$H} NMR (101 MHz, $CD_3OD$): δ = 196.1, 171.2, 143.0, 139.5, 129.5, 129.4, 122.9, 114.67, 55.6, 40.9, 33.6, 20.7 ppm.
HRMS (ESI) calcd for $[C_{13}H_{16}NO_2]^+$: 218.11810, found 218.11770.

**2,3,3-Trimethyl-1-(pent-4-yn-1-yl)-3*H*-indol-1-ium iodide (7c)**

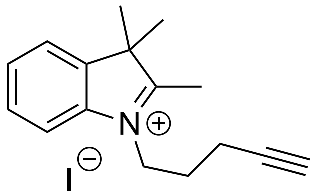

5-iodo-pent-1-yne was prepared according to previously reported methods.[5] 2,3,3-trimethylindolenine (200 μL, 1.26 mmol, 1.0 equiv.) and 5-iodo-pent-1-yne (731 mg, 3.77 mmol, 3.0 equiv.) were dissolved in $CH_3CN$ (5 mL) and heated under reflux for 16 h. Solvents were removed under reduced pressure and the residue was washed with diethyl ether multiple times to yield a pale pink solid (389 mg, 88%).

$^1$H NMR (400 MHz, $(CD_3)_2SO$): δ = 7.99 – 7.96 (m, 1H), 7.87 – 7.84 (m, 1H), 7.64 – 7.62 (m, 2H), 4.51 (t, *J* = 7.7 Hz, 2H), 2.95 (t, *J* = 2.7 Hz, 1H), 2.86 (s, 3H), 2.43 (td, *J* = 7.2, 2.7 Hz, 2H), 2.07 (p, *J* = 7.2 Hz, 2H), 1.55 (s, 6H) ppm.
$^{13}$C{$^1$H} NMR (101 MHz, $(CD_3)_2SO$): δ = 197.0, 141.8, 141.1, 129.4, 128.9, 123.5, 115.2, 83.1, 72.4, 54.3, 46.8, 26.0, 22.0, 15.2, 14.2 ppm.
HRMS (ESI) calcd for $[C_{16}H_{20}N]^+$: 226.15958, found 226.15904.

**2,3,3-Trimethyl-1-(pent-4-yn-1-yl)-5-(trifluoromethyl)-3*H*-indol-1-ium iodide (7d)**

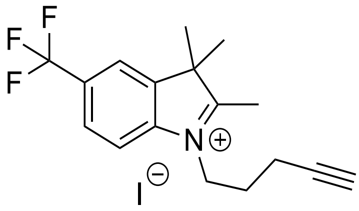

2,3,3-trimethyl-5-(trifluoromethyl)-3*H*-indole was prepared according to previously reported methods.[6] 2,3,3-trimethyl-5-(trifluoromethyl)-3*H*-indole (850 mg, 3.74 mmol, 1.0 equiv.) and 5-iodo-pent-1-yne (1.81 g, 9.35 mmol, 2.5 equiv.) were dissolved in $CH_3CN$ (10 mL) and heated under reflux for 36 h. Solvents were removed under reduced pressure and the residue was washed with diethyl ether multiple times to yield the product as a pale purple solid (580 mg, 37%).

$^1$H NMR (400 MHz, $CD_3CN$): δ = 8.08 (s, 1H), 7.98 – 7.93 (m, 2H), 4.48 (t, *J* = 7.8 Hz, 2H), 2.81 (s, 3H), 2.42 (td, *J* = 6.9, 2.7 Hz, 2H), 2.35 (t, *J* = 2.7 Hz, 1H), 2.14 (p, *J* = 7.3 Hz, 2H), 1.59 (s, 6H).
$^{13}$C{$^1$H} NMR (101 MHz, $CD_3CN$): δ = 160.3 (q, *J* = 37.1 Hz), 145.0, 143.8, 127.9 (q, 3.8 Hz), 126.2, 123.5, 122.1 (q, 3.8 Hz), 117.2, 115,6, 83.2, 71.8, 56.3, 48.5, 27.0, 22.5, 16.3.
HRMS (ESI) calcd for $[C_{17}H_{19}F_3N]^+$: 294.14696, found 294.14606.

**6-((4-(Aminomethyl)benzyl)oxy)-9*H*-purin-2-amine-6-((4-(azidomethyl)benzyl)-oxy)-9*H*-purin-2-amine (10)**

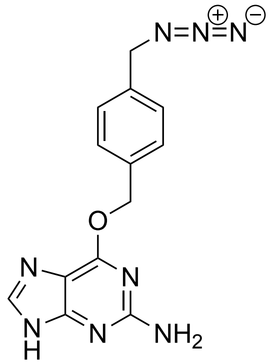

SNAP-amine[7] and $FSO_2N_3$[8] were prepared according to previously reported methods. SNAP-amine (30 mg, 0.11 mmol, 1.0 equiv.) was dissolved in DMF/MTBE/$H_2O$ (94:5:1, 1.5 mL) and $FSO_2N_3$ solution (1.35 equiv., 307 μL of a 488 mM solution) and aqueous potassium hydrogen carbonate solution (3 M, 1.5 mL) were added. The mixture was stirred for 3 h at 18 °C. Ethyl acetate was added and the organic phase was washed trice with brine, once with water and again with brine. The extracted organic phase was dried over $Na_2SO_4$, filtered, and solvents were removed under reduced pressure to yield the product as an off-white solid (32 mg, 97%).

$^1$H NMR (400 MHz, $(CD_3)_2SO$): δ = 12.43 (s, 1H), 7.82 (s, 1H), 7.54 (d, *J* = 8.1 Hz, 2H), 7.40 (d, *J* = 8.2 Hz, 2H), 6.29 (s, 2H), 5.49 (s, 2H), 4.46 (s, 2H) ppm.
$^{13}$C{$^1$H} NMR (101 MHz, $(CD_3)_2SO$): δ = 159.6, 136.7, 135.4, 129.0, 128.7, 128.54, 128.50, 128.4, 66.3, 53.3 ppm.
HRMS (ESI) calcd for $[C_{13}H_{12}N_8O+H]^+$: 297.12068, found 297.12043.

**Synthesis of 5-endo *trig* carbocyanines**

**General procedure to synthesize 5-endo *trig* hydroxyethyl probes 2a and 2b**

Compound **5** (50 mg, 0.15 mmol, 1.0 equiv.) and **8** (45 mg, 0.18 mmol, 1.2 equiv.) were dissolved in $Ac_2O$ (1 mL) in a microwave vial. The solution was heated to 120 °C for 1 h in the microwave. The mixture was cooled to room temperature and a solution of **7** (0.17 mmol, 1.1 equiv.) in pyridine (1 mL) was added dropwise. The mixture was stirred for 30 min at 18 °C. The color of the solution turned from green to blue. Solvents were removed under reduced pressure. The residue was washed with diethyl ether and then stirred in a mixture of $CH_3OH$ (2 mL) and saturated $K_2CO_3$ (3 mL) for 2 h at 18 °C to deprotect the acetylated alcohol. Solvents were removed under reduced pressure and the residue was dissolved in $CH_2Cl_2$ and washed three times with water and twice with brine. The organic layer was dried over $Na_2SO_4$, filtered, and purified via reverse-phase column chromatography (10 → 70% $CH_3CN$ in dd$H_2O$ + 0.1% TFA over 35 min) to afford the product as a fine blue powder.

**1-(2-Hydroxyethyl)-3,3-dimethyl-2-((1*E*,3*E*)-5-((*E*)-1,3,3-trimethylindolin-2-ylidene)-penta-1,3-dien-1-yl)-3*H*-indol-1-ium 2,2,2-trifluoroacetate (2a)**

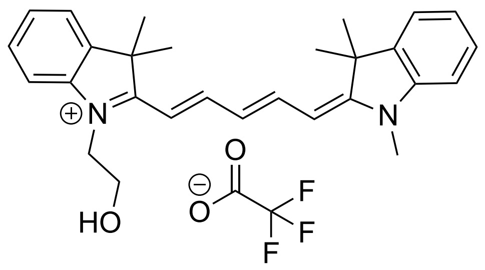

Yield: 54 mg, 68%

$^1$H NMR (400 MHz, $CD_3OD$): δ = 8.23 (td, *J* = 13.2 Hz, 5.4 Hz, 2H), 7.47 (d, *J* = 7.4 Hz, 2H), 7.40 (q, *J* = 7.6 Hz, 7.1 Hz, 2H), 7.33 – 7.22 (m, 4H), 6.59 (t, *J* = 12.4 Hz, 1H), 6.35 (d, *J* = 13.7 Hz, 1H), 6.25 (d, *J* = 13.7 Hz, 1H), 4.23 (t, *J* = 5.4 Hz, 2H), 3.94 (t, *J* = 5.4 Hz, 2H), 3.61 (s, 3H), 1.74 (s, 6H), 1.72 (s, 6H) ppm.

$^{13}$C{$^1$H} NMR (101 MHz, $CD_3OD$): δ = 175.7, 175.2, 155.5, 155.4, 155.3, 155.2, 144.3, 144.1, 142.6, 142.5, 129.7, 129.6, 126.2, 126.2, 126.2, 123.3, 123.3, 112.4, 111.8, 60.0, 50.6, 50.4, 47.7, 31.4, 28.0, 27.8 ppm.

HRMS (ESI) calcd for $[C_{28}H_{33}N_2O]^+$: 413.25874, found 413.25813.

**1-(2-Hydroxyethyl)-3,3-dimethyl-2-((1*E*,3*E*)-5-((*E*)-1,3,3-trimethyl-5-(trifluoromethyl) indolin-2-ylidene)penta-1,3-dien-1-yl)-3*H*-indol-1-ium 2,2,2-trifluoroacetate (2b)**

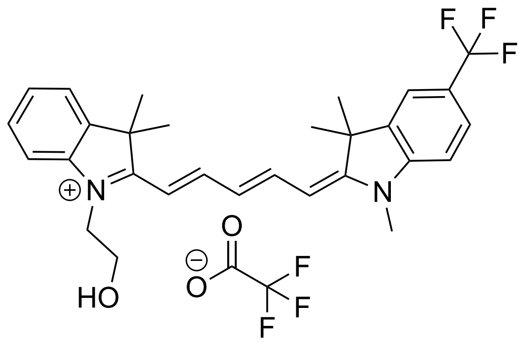

Yield: 71 mg, 79%

$^1$H NMR (400 MHz, $CD_3OD$): δ = 8.31 (t, *J* = 13.1 Hz, 1H), 8.19 (t, *J* = 13.1 Hz, 1H), 7.69 (s, 1H), 7.65 (d, *J* = 10.0 Hz, 1H), 7.54 (d, *J* = 7.4 Hz, 1H), 7.47 – 7.41 (m, 2H), 7.37 – 7.31 (m, 1H), 7.27 (d, *J* = 8.3 Hz, 1H), 6.63 (t, *J* = 12.4 Hz, 1H), 6.54 (d, *J* = 14.2 Hz, 1H), 6.14 (d, *J* = 13.3 Hz, 1H), 4.33 (t, *J* = 5.3 Hz, 2H), 3.98 (t, *J* = 5.3 Hz, 2H), 3.53 (s, 3H), 1.76 (s, 6H), 1.73 (s, 6H) ppm.

$^{13}$C{$^1$H} NMR (101 MHz, $CD_3OD$): δ = 178.8, 172.7, 157.2, 154.0, 147.7, 143.7, 143.3, 142.7, 129.8, 127.6, 127.5, 127.3 (q, *J* = 3.9 Hz), 126.7 (q, *J* = 32.6 Hz), 125.9 (q, *J* = 270.8 Hz), 123.5, 120.3 (q, *J* = 3.7 Hz), 113.5, 110.9, 107.6, 103.5, 60.2, 51.6, 51.5, 49.6, 48.3, 27.9, 27.7 ppm.

HRMS (ESI) calcd for $[C_{29}H_{32}F_3N_2O]^+$: 481.24612, found 481.24606.

**2-((1*E*,3*E*)-5-((*E*)-3,3-Dimethyl-1-(pent-4-yn-1-yl)indolin-2-ylidene)penta-1,3-dien-1-yl)-3,3-dimethyl-1-(2-(methylamino)-2-oxoethyl)-3*H*-indol-1-ium 2,2,2-trifluoroacetate (14)**

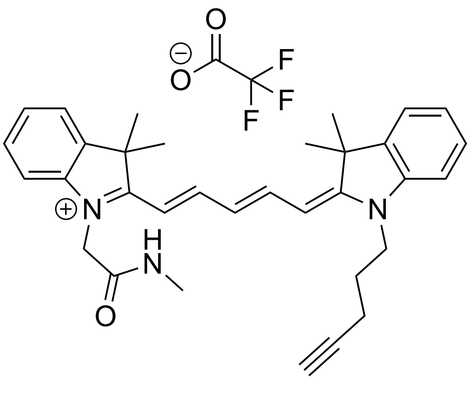

Compound **12** (100 mg, 0.28 mmol, 1.0 equiv.) and **8** (86 mg, 0.34 mmol, 1.2 equiv.) were dissolved in $Ac_2O$ (2 mL) in a microwave vial. The solution was heated to 120 °C for 1 h in the microwave. The mixture was cooled to room temperature and a solution of **7c** (108 mg, 0.31 mmol, 1.1 equiv.) in pyridine (2 mL) was added dropwise. The mixture was stirred for 30 min at 18 °C. Solvents were removed under reduced pressure and the residue was purified via reverse phase column chromatography (10 $\rightarrow$ 70% $CH_3CN$ in dd$H_2O$ + 0.1% TFA over 35 min) to afford the product as a fine blue powder (87 mg, 52%).

$^1$H NMR (400 MHz, $CD_3OD$): $\delta$ = 8.32 – 8.22 (m, 2H), 7.52 (d, *J* = 7.5 Hz, 1H), 7.48 (d, *J* = 7.5 Hz, 1H), 7.46 – 7.41 (m, 1H), 7.40 – 7.34 (m, 2H), 7.33 – 7.29 (m, 1H), 7.24 (t, *J* = 7.9 Hz, 1H), 7.15 (d, *J* = 7.5 Hz, 1H), 6.57 (t, *J* = 12.4 Hz, 1H), 6.42 (d, *J* = 13.9 Hz, 1H), 6.13 (d, *J* = 13.6 Hz, 1H), 4.79 (s, 2H), 4.26 (t, *J* = 7.4 Hz, 2H), 2.82 (s, 3H), 2.50 (t, *J* = 2.6 Hz, 1H), 2.38 (td, *J* = 6.6, 2.6 Hz, 2H), 2.04 – 1.98 (m, 2H), 1.77 (s, 6H), 1.74 (s, 6H) ppm.

$^{13}$C NMR (101 MHz, $CD_3OD$) $\delta$ = 175.9, 175.3, 168.3, 156.4, 155.5, 144.1, 143.3, 142.8, 142.1, 129.8, 129.6, 127.9, 126.8, 126.0, 123.5, 123.3, 112.3, 111.3, 105.2, 104.1, 83.7, 71.4, 50.9, 50.5, 47.2, 44.0, 28.1, 27.8, 27.3, 26.6, 16.5 ppm.

HRMS (ESI) calcd for $[C_{33}H_{38}N_3O]^+$: 492.30149, found 492.30118.

#### Synthesis of 5-exo *trig* carbocyanines

#### General procedure for the synthesis of 5-exo *trig* Cy5 methyl esters (9)

In a flame-dried flask, compound **6** (50 mg, 0.16 mmol, 1.0 equiv.) was dissolved in anhydrous $CH_3CN$ (5 mL) and oxalyl chloride (19 µL, 0.22 mmol, 1.3 equiv.) was added dropwise. The solution was stirred for 1 h at 18 °C. Anhydrous $CH_3OH$ (5 mL) was added and the solution was stirred for additional 15 min. Solvents were removed under reduced pressure and the residue was used directly without further purification. The residue was dissolved in $Ac_2O$ (1 mL) in a microwave vial. Compound **8** (51 mg, 0.20 mmol, 1.2 equiv.) was added and the solution was heated to 120 °C for 1 h in the microwave. The mixture was cooled to room temperature and a solution of **7** (0.18 mmol, 1.1 equiv.) in pyridine (1 mL) was added dropwise. The mixture was heated in the microwave at 110 °C for 30 min. Solvents were removed under reduced pressure and the residue was purified via reverse-phase column chromatography (10 → 55% $CH_3CN$ in dd$H_2O$ + 0.1% TFA over 35 min) to afford the product as a fine blue powder.

##### 3-(2-Methoxy-2-oxoethyl)-1,3-dimethyl-2-((1*E*,3*E*)-5-((*E*)-1,3,3-trimethylindolin-2-ylidene)penta-1,3-dien-1-yl)-3*H*-indol-1-ium 2,2,2-trifluoroacetate (9a)

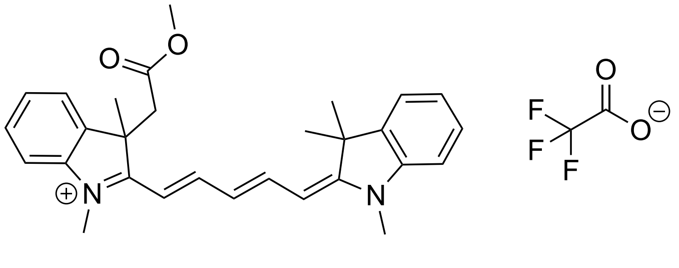

Yield: 57 mg, 61%

$^1$H NMR (400 MHz, $CD_3OD$): δ = 8.27 – 8.16 (m, 2H), 7.49 – 7.39 (m, 4H), 7.30 – 7.21 (m, 4H), 6.60 (t, *J* = 12.4 Hz, 1H), 6.27 (m, 2H), 3.63 (d, *J* = 17,4 Hz, 1H), 3.63 (s, 3H), 3,61 (s, 3H), 3,42 (d, *J* = 17.4 Hz, 1 H), 3.40 (s, 3H), 1.72 (s, 3H), 1.72 (s, 3H), 1.67 (s, 3H) ppm.

$^{13}$C{$^1$H} NMR (101 MHz, $CD_3OD$): δ = 175.3, 173.7, 171.4, 155.4, 154.7, 145.3, 144.2, 142.5, 140.1, 130.0, 129.7, 126.6, 126.2, 126.0, 123.3, 123.0, 111.8, 111.7, 104.4, 104.3, 52.1, 51.4, 50.4, 44.9, 31.5, 31.4, 27.8, 27.6 ppm.

HRMS (ESI) calcd for $[C_{29}H_{33}N_2O_2]^+$: 441.25420, found 441.25313.

**3-(2-Methoxy-2-oxoethyl)-1,3-dimethyl-2-((1*E*,3*E*)-5-((*E*)-1,3,3-trimethyl-5-(trifluoro-methyl)indolin-2-ylidene)penta-1,3-dien-1-yl)-3*H*-indol-1-ium 2,2,2-trifluoroacetate (9b)**

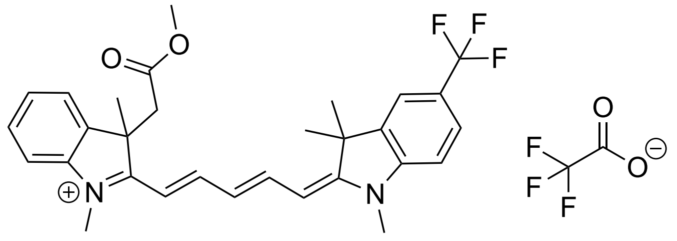

Yield: 67 mg, 64%

$^1$H NMR (400 MHz, $CD_3OD$): $\delta$ = 8.32 – 8.19 (m, 2H), 7.73 (s, 1H), 7.68 (d, *J* = 8.4 Hz, 1H), 7.55 (d, *J* = 7.5 Hz, 1H), 7.50 – 7.42 (m, 2H), 7.36 – 7.30 (m, 2H), 6.66 (t, *J* = 12.4 Hz, 1H), 6.50 (d, *J* = 14.3 Hz, 1H), 6.17 (d, *J* = 13.3 Hz, 1H), 3.76 (s, 3H), 3.70 (d, *J* = 17.5 Hz, 1H), 3.55 (s, 3H), 3.50 (d, *J* = 17.7 Hz, 1H), 3.41 (s, 3H), 1.75 (s, 3H), 1.74 (s, 3H), 1.70 (s, 3H) ppm.

$^{13}$C{$^1$H} NMR (101 MHz, $CD_3OD$): $\delta$ = 175.3, 171.4, 169.9, 155.0, 152.8, 146.3, 143.6, 141.3, 139.4, 128.8, 126.2, 125.95, 125.90 (q, *J* = 3.7 Hz), 125.2 (q, *J* = 32 Hz), 123.2, 118.9 (q, *J* = 3.7 Hz), 111.5, 109.5, 105.6, 102.1, 50.9, 50.8, 43.3, 30.8, 29.7, 26.52, 26.49, 25.8 ppm.

HRMS (ESI) calcd for $[C_{30}H_{32}F_3N_2O_2]^+$: 509.24159, found 509.24065.

**2-((1*E*,3*E*)-5-((*E*)-3,3-Dimethyl-1-(pent-4-yn-1-yl)indolin-2-ylidene)penta-1,3-dien-1-yl)-3-(2-methoxy-2-oxoethyl)-1,3-dimethyl-3*H*-indol-1-ium 2,2,2-trifluoroacetate (9c)**

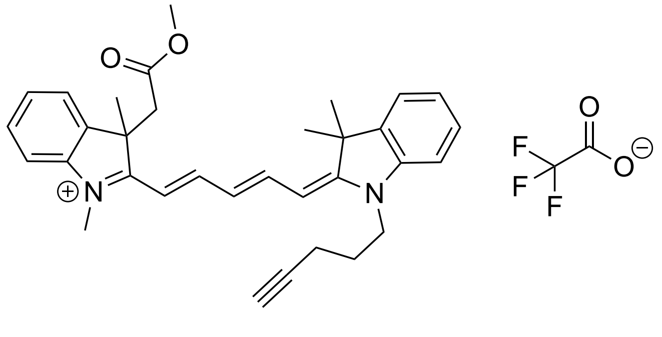

Yield: 67 mg, 66%

$^1$H NMR (400 MHz, $CD_3OD$): $\delta$ = 8.28 – 8.18 (m, 2H), 7.49 (d, *J* = 7.5 Hz, 2H), 7.45 – 7.38 (m, 2H), 7.31 (d, *J* = 7.9 Hz, 2H), 7.25 (t, *J* = 8.0 Hz, 2H), 6.59 (t, *J* = 12.4 Hz, 1H), 6.35 – 6.30 (m, 2H), 4.20 (t, *J* = 7.4 Hz, 2H), 3.65 (s, 3H), 3.64 (d, *J* = 17.3 Hz, 1H), 3.44 (d, *J* = 17.4 Hz, 1H), 3.40 (s, 3H), 2.50 (t, *J* = 2.6 Hz, 1H), 2.37 (td, *J* = 6.6, 2.6 Hz, 2H), 2.03 – 1.96 (m, 2H), 1.73 (s, 6H), 1.73 (s, 3H), 1.68 (s, 3H) ppm.

$^{13}C\{^{1}H\}$ NMR (101 MHz, $CD_3OD$): δ = 174.4, 174.2, 171.4, 155.3, 155.0, 145.3, 143.6, 142.5, 140.2, 130.1, 129.7, 126.6, 126.3, 126.1, 123.4, 123.0, 111.9, 111.7, 104.9, 104.0, 83.8, 71.3, 52.1, 51.6, 50.4, 44.9, 43.6, 31.7, 28.0, 27.5, 27.2, 16.5 ppm.
HRMS (ESI) calcd for $[C_{33}H_{37}N_2O_2]^+$: 493.28550, found 493.28501.

**2-((1*E*,3*E*)-5-((*E*)-3,3-Dimethyl-1-(pent-4-yn-1-yl)-5-(trifluoromethyl)indolin-2-ylidene) penta-1,3-dien-1-yl)-3-(2-methoxy-2-oxoethyl)-1,3-dimethyl-3*H*-indol-1-ium 2,2,2-tri-fluoroacetate (9d)**

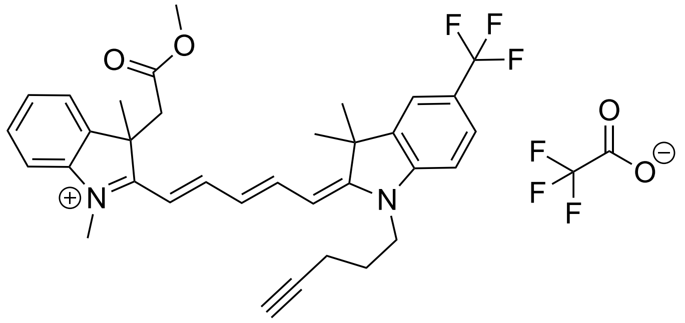

Yield: 71 mg, 63%
$^{1}H$ NMR (400 MHz, $CD_3OD$): δ = 8.33 – 8.18 (m, 2H), 7.73 (s, 1H), 7.67 (d, *J* = 8.3 Hz, 1H), 7.56 (d, *J* = 7.1 Hz, 1H), 7.51 – 7.44 (m, 2H), 7.35 (t, *J* = 7.4 Hz, 2H), 6.64 (t, *J* = 12.4 Hz, 1H), 6.53 (d, *J* = 14.4 Hz, 1H), 6.26 (d, *J* = 13.3 Hz, 1H), 4.15 (t, *J* = 7.4 Hz, 2H), 3.77 (s, 3H), 3.71 (d, *J* = 17.5 Hz, 1H), 3.51 (d, *J* = 17.5 Hz, 1H), 3.41 (s, 3H), 2.49 (t, *J* = 2.6 Hz, 1H), 2.37 (td, *J* = 6.7, 2.6 Hz, 2H), 1.98 (p, *J* = 7.3 Hz, 2H), 1.76 (s, 3H), 1.75 (s, 3H), 1.70 (s, 3H) ppm.
$^{13}C\{^{1}H\}$ NMR (101 MHz, $CD_3OD$): δ = 177.1, 171.7, 171.3, 156.4, 153.9, 147.1, 144.9, 142.7, 140.9, 130.3, 127.7, 127.5, 127.3 (q, *J* = 3.7 Hz), 126.6 (q, *J* = 32 Hz), 125.9 (q, *J* = 270.8 Hz), 123.2, 120.43 (q, *J* = 3.7 Hz), 113.0, 110.9, 107.4, 103.3, 83.8, 71.3, 52.4, 52.2, 44.7, 43.4, 32.3, 28.0, 27.1, 26.9, 16.5 ppm.
HRMS (ESI) calcd for $[C_{34}H_{36}F_3N_2O_2]^+$: 561.27289, found 561.27194.

**General procedure for the synthesis of 5-exo *trig* Cy5 alcohols (4)**

Methyl ester **9** (40 mg, 1.0 equiv.) was dissolved in anhydrous THF (3 mL) in a flame-dried flask and cooled to 0 °C. $LiAlH_4$ (4.0 equiv.) was added, and the solution was stirred for 1 h at 0 °C, followed by 16 h at 18 °C. The mixture was cooled again to 0 °C and water was added. The aqueous phase was extracted trice with $CH_2Cl_2$, and the combined organic phases were washed with brine and dried over $Na_2SO_4$. DDQ (1.0 equiv.) was added to re-oxidize the conjugated system of the carbocyanine core. Solvents were removed under reduced pressure and the residue was purified via reverse-phase column chromatography (10 $\rightarrow$ 65% $CH_3CN$ in dd$H_2O$ + 0.1% TFA over 35 min) to afford the product as a fine blue powder.

**3-(2-Hydroxyethyl)-1,3-dimethyl-2-((1*E*,3*E*)-5-((*E*)-1,3,3-trimethylindolin-2-ylidene)-penta-1,3-dien-1-yl)-3*H*-indol-1-ium 2,2,2-trifluoroacetate (4a)**

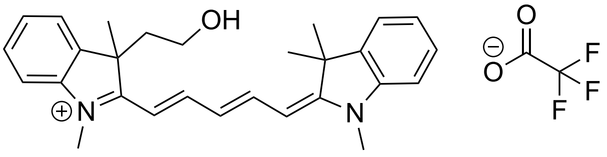

Yield: 21 mg, 55%

$^1$H NMR (400 MHz, $CD_3OD$): $\delta$ = 8.24 (t, $J$ = 13.1 Hz, 2H), 7.50 – 7.40 (m, 4H), 7.30 – 7.24 (m, 4H), 6.62 (t, $J$ = 12.4 Hz, 1H), 6.30 (d, $J$ = 4.9 Hz, 1H), 6.26 (d, $J$ = 4.9 Hz, 1H), 3.62 (s, 3H), 3.60 (s, 3H), 3.14 – 3.06 (m, 1H), 3.04 – 2.97 (m, 1H), 2.75 – 2.63 (m, 1H), 2.56 – 2.35 (m, 1H), 1.73 (s, 9H) ppm.

$^{13}$C{$^1$H} NMR (101 MHz, $CD_3OD$): $\delta$ = 175.4, 173.8, 155.4, 155.0, 145.0, 144.3, 142.5, 140.2, 129.9, 129.7, 126.6, 126.3, 126.2, 123.4, 123.3, 111.82, 111.80, 104.8, 104.4, 59.2, 57.5, 53.0, 50.5, 44.5, 31.5, 28.2, 27.8 ppm.

HRMS (ESI) calcd for $[C_{28}H_{33}N_2O]^+$: 413.25874, found 413.25819.

**3-(2-Hydroxyethyl)-1,3-dimethyl-2-((1*E*,3*E*)-5-((*E*)-1,3,3-trimethyl-5-(trifluoro-methyl)-indolin-2-ylidene)penta-1,3-dien-1-yl)-3*H*-indol-1-ium 2,2,2-trifluoro-acetate (4b)**

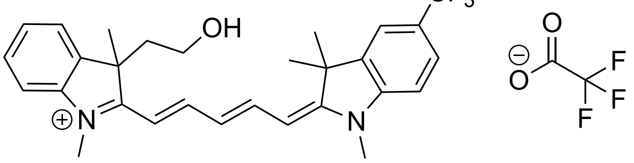

Yield: 24 mg, 63%

$^1$H NMR (400 MHz, $CD_3OD$): δ = 8.33 (t, *J* = 13.1 Hz, 1H), 8.21 (t, *J* = 13.1 Hz, 1H), 7.73 (s, 1H), 7.67 (d, *J* = 8.4 Hz, 1H), 7.55 (d, *J* = 7.6 Hz, 1H), 7.50 (t, *J* = 7.0 Hz, 1H), 7.45 – 7.36 (m, 2H), 7.30 (d, *J* = 8.4 Hz, 1H), 6.67 (t, *J* = 12.5 Hz, 1H), 6.50 (d, *J* = 14.2 Hz, 1H), 6.17 (d, *J* = 13.3 Hz, 1H), 3.72 (s, 3H), 3.55 (s, 3H), 3.12 – 3.01 (m, 2H), 2.80 – 2.72 (m, 1H), 2.51 (m, 1H), 1.75 (s, 3H), 1.74 (s, 6H) ppm.
$^{13}$C{$^1$H} NMR (101 MHz, $CD_3OD$): δ = 177.3, 172.6, 156.7, 153.9, 147.7, 144.7, 142.6, 140.9, 130.1, 127.6, 127.5, 127.3 (q, *J* = 3.7 Hz), 126.6 (q, *J* = 32.7 Hz), 124.6, 123.7, 120.3 (q, *J* = 3.7 Hz), 113.0, 110.9, 107.6, 103.4, 59.1, 57.5, 54.0, 44.5, 32.2, 31.1, 27.9 ppm.
HRMS (ESI) calcd for $[C_{29}H_{32}F_3N_2O]^+$: 481.24612, found 481.24588.

**2-((1*E*,3*E*)-5-((*E*)-3,3-Dimethyl-1-(pent-4-yn-1-yl)-5-(trifluoromethyl)indolin-2-ylidene)-penta-1,3-dien-1-yl)-3-(2-hydroxyethyl)-1,3-dimethyl-3*H*-indol-1-ium 2,2,2-trifluoro-acetate (4c)**

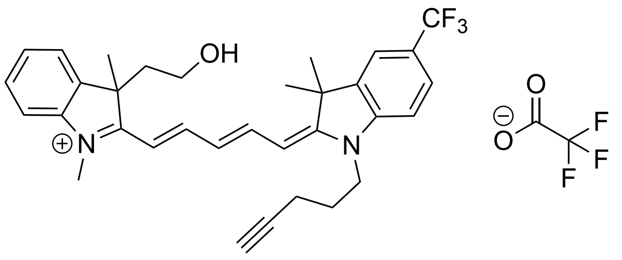

Yield: 26 mg, 69%
$^1$H NMR (400 MHz, $CD_3OD$): δ = 8.34 (t, *J* = 13.2 Hz, 1H), 8.20 (t, *J* = 13.0 Hz, 1H), 7.73 (s, 1H), 7.67 (d, *J* = 8.4 Hz, 1H), 7.56 (d, *J* = 6.7 Hz, 1H), 7.53 – 7.48 (m, 1H), 7.46 (d, *J* = 8.1 Hz, 1H), 7.40 (td, *J* = 7.3, 1.3 Hz, 1H), 7.33 (d, *J* = 8.4 Hz, 1H), 6.66 (t, *J* = 12.4 Hz, 1H), 6.53 (d, *J* = 14.3 Hz, 1H), 6.26 (d, *J* = 13.1 Hz, 1H), 4.15 (t, *J* = 7.4 Hz, 2H), 3.74 (s, 3H), 3.15 – 3.00 (m, 2H), 2.82 – 2.72 (m, 1H), 2.57 – 2.50 (m, 1H), 2.50 (t, *J* = 2.6 Hz, 1H), 2.37 (td, *J* = 6.6, 2.6 Hz, 2H), 1.98 (p, *J* = 6.8 Hz, 2H), 1.75 (s, 9H) ppm.
$^{13}$C{$^1$H} NMR (101 MHz, $CD_3OD$): δ = 177.7, 171.5, 156.8, 153.5, 147.2, 144.6, 142.6, 141.0, 130.1, 127.74, 127.65, 127.3 (q, *J* = 3.7 Hz), 126.6 (q, *J* = 32.6 Hz), 124.6, 123.7, 120.4 (q, *J* = 3.7 Hz), 113.2, 110.9, 108.0, 103.2, 83.9, 71.3, 59.2, 54.2, 44.5, 43.4, 32.3, 28.1, 27.9, 26.9, 16.5 ppm.
HRMS (ESI) calcd for $[C_{33}H_{36}F_3N_2O]^+$: 533.27742, found 533.27686.

**2-((1*E*,3*E*)-5-((*E*)-3,3-Dimethyl-1-(pent-4-yn-1-yl)indolin-2-ylidene)penta-1,3-dien-1-yl) 1,3-dimethyl-3-(2-(methylamino)-2-oxoethyl)-3*H*-indol-1-ium 2,2,2-trifluoroacetate (12)**

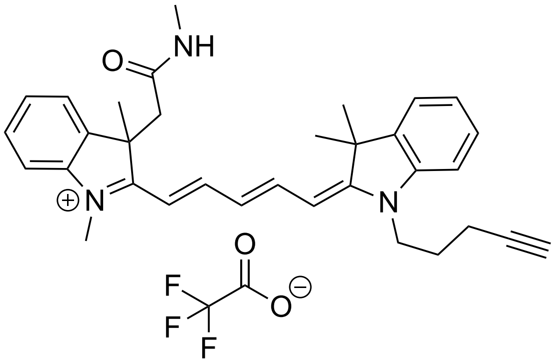

Compound **9c** (25 mg, 0.04 mmol, 1.0 equiv.) was dissolved in anhydrous THF (3 mL) and methyl amine (2.0 M solution in THF, 5 mL) was added. The solution was stirred at 70 °C for 16 h. Solvents were removed under reduced pressure and the residue was purified via reverse-phase column chromatography (10 → 55% $CH_3CN$ in dd$H_2O$ + 0.1% TFA over 35 min) to afford the product as a fine blue powder (24 mg, 96%).

$^1$H NMR (400 MHz, $CD_3OD$): $\delta$ = 8.18 (t, $J$ = 13.3 Hz, 2H), 7.48 – 7.33 (m, 4H), 7.37 – 7.21 (m, 4H), 6.58 (t, $J$ = 12.4 Hz, 1H), 6.36 (d, $J$ = 14.0 Hz, 1H), 6.28 (d, $J$ = 13.5 Hz, 1H), 4.17 (t, $J$ = 7.5 Hz, 2H), 3.67 (s, 3H), 3.40 (d, $J$ = 15.6 Hz, 1H), 3.21 (d, $J$ = 15.6 Hz, 1H), 2.50 (t, $J$ = 2.6 Hz, 1H), 2.43 (s, 3H), 2.36 (td, $J$ = 6.6, 2.6 Hz, 2H), 1.99 (p, $J$ = 6.8 Hz, 2H), 1.72 (s, 3H), 1.72 (s, 3H) 1.68 (s, 3H) ppm.
$^{13}$C{$^1$H} NMR (101 MHz, $CD_3OD$): $\delta$ = 175.7, 173.4, 171.4, 155.3, 154.5, 145.1, 143.7, 142.3, 140.40 130.0, 129.7, 126.5, 126.4, 125.8, 123.4, 123.2, 112.3, 111.4, 105.7, 103.4, 83.8, 71.3, 52.4, 50.2, 46.7, 43.5, 31.8, 28.1, 27.4, 27.2, 26.0, 16.5 ppm.
HRMS (ESI) calcd for $[C_{33}H_{38}N_3O]^+$: 492.30149, found 492.30128.

**(4*R*,7*R*,10*S*,13*S*,19*S*,*E*)-7-((1*H*-indol-3-yl)methyl)-10-(4-(4-(3-((*E*)-3,3-dimethyl-2-((2*E*,4*E*)-5-(1,3a,8-trimethyl-2-oxo-2,3,3a,8-tetrahydropyrrolo[2,3-*b*]indol-8a(1*H*)-yl)penta-2,4-dien-1-ylidene)indolin-1-yl)propyl)-1*H*-1,2,3-triazol-1-yl)butyl)-4-(4-hydroxyphenyl)-8,13,15,19-tetramethyl-1-oxa-5,8,11-triazacyclononadec-15-ene-2,6,9,12-tetraone (17-actin)**

Boc-Lys-jasplakinolide (7 mg, 0.009 mmol, 1.0 equiv.) was dissolved in TFA:$CH_2Cl_2$ (3:7 v/v, 0.3 mL) and stirred at 18 °C for 7.5 min. Toluene (0.6 mL) was added and solvents were removed under reduced pressure. Co-evaporation was carried out twice with 0.4 mL THF, and once with 0.4 mL THF and 10 μL triethylamine. The residue was used directly for the diazotizing reaction. The residue was dissolved in DMF/MTBE/$H_2O$ (94:5:1, 0.3 mL) and $FSO_2N_3$ solution (1.35 equiv., 25 μL of a 488 mM solution) and aqueous potassium hydrogen carbonate solution (3 M, 0.3 mL) were added. The mixture was stirred for 1 h at 18 °C. Ethyl acetate was added and the organic phase was washed trice with brine, once with water and again with brine. The extracted organic phase was dried over $Na_2SO_4$, filtered, and solvents were removed under reduced pressure to yield azide-modified Lys-jasplakinolide, which was used directly in the Click reaction without further purification.
In a flame-dried flask, **12** (2.7 mg, 0.0045 mmol, 0.5 equiv.) and azide-modified Lys-jasplakinolide were dissolved in degassed THF (2 mL) and the solution was degassed for an additional hour by bubbling argon through the solution while stirring vigorously. $Cu(PPh_3)_2NO_3$ (1.5 mg, 0.0023 mmol, 0.25 equiv.) and triethylamine (5 μL) were added and the solution was stirred for 16 h at room temperature. Solvents were removed under reduced pressure and the residue was purified via reverse-phase column chromatography (10 → 90% $CH_3CN$ in dd$H_2O$ over 35 min).

Yield: 4.5 mg (42 %)
HRMS (ESI) calcd for $[C_{71}H_{87}N_{10}O_7]^+$: 1191.67537, found 1191.67359.

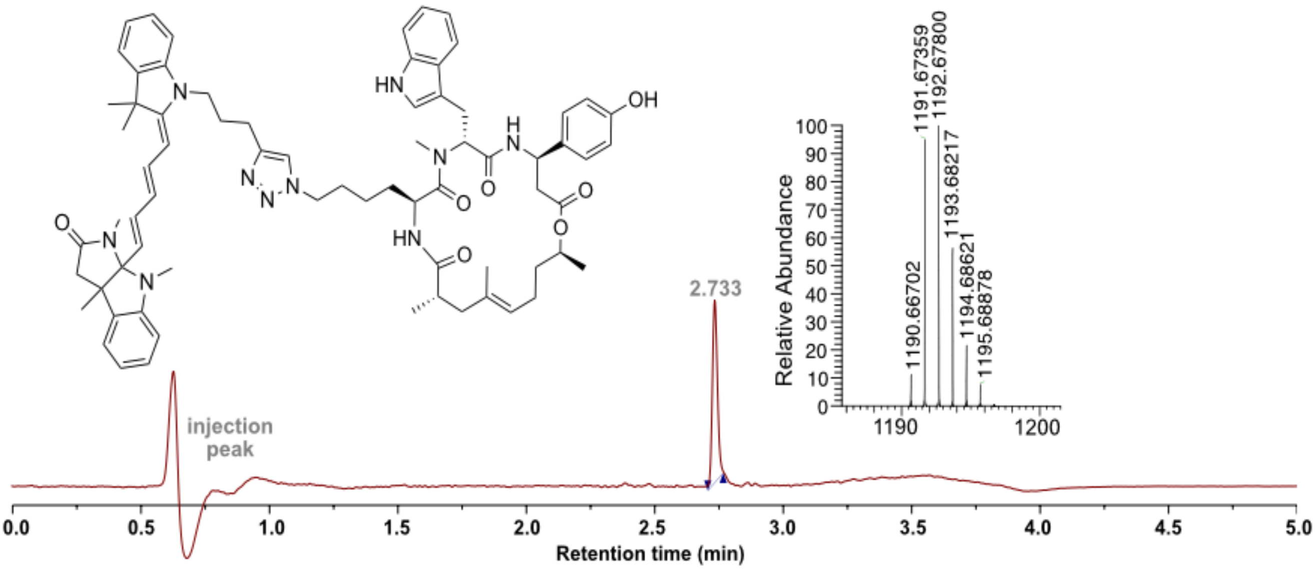

**8a-((1*E*,3*E*)-5-((*E*)-3,3-dimethyl-1-(3-(1-(4-(4-(6-(4-methylpiperazin-1-yl)-1*H*,3'*H*-[2,5'-bibenzo[*d*]imidazol]-2'-yl)phenoxy)butyl)-1*H*-1,2,3-triazol-4-yl)propyl)-indolin-2-ylidene)penta-1,3-dien-1-yl)-1,3a,8-trimethyl-3,3a,8,8a-tetrahydro-pyrrolo[2,3-*b*]indol-2(1*H*)-one (18-DNA)**

Boc-C4-Hoechst (6.3 mg, 0.011 mmol, 1.0 equiv.) was dissolved in dry $CH_2Cl_2$ (0.8 mL) and TFA (0.2 mL) was added dropwise. The mixture was stirred at 18 °C for 30 min. Solvents were removed under reduced pressure and co-evaporation with THF (2x 0.4 mL) was carried out twice. The residue was used directly for the diazotizing reaction. The residue was dissolved in DMF/MTBE/$H_2O$ (94:5:1, 0.3 mL) and $FSO_2N_3$ solution (1.35 equiv., 30 $\mu$L of a 488 mM solution) and aqueous potassium hydrogen carbonate solution (3 M, 0.3 mL) were added. The mixture was stirred for 1 h at 18 °C. Ethyl acetate was added and the organic phase was washed trice with brine, once with water and again with brine. The extracted organic phase was dried over $Na_2SO_4$, filtered, and solvents were removed under reduced pressure to yield azide-modified C4-Hoechst, which was used directly in the Click reaction without further purification.
In a flame-dried flask, **12** (3.2 mg, 0.0053 mmol, 0.5 equiv.) and azide-modified C4-Hoechst were dissolved in degassed THF (2 mL) and the solution was degassed for an additional hour by bubbling argon through the solution while stirring vigorously. $Cu(PPh_3)_2NO_3$ (1.7 mg, 0.0023 mmol, 0.25 equiv.) and triethylamine (5 $\mu$L) were added and the solution was stirred for 16 h at room temperature. Solvents were removed under reduced pressure and the residue was purified via reverse-phase column chromatography (10 $\rightarrow$ 55% $CH_3CN$ in dd$H_2O$ + 0.1% TFA over 35 min).

Yield: 2.9 mg (27 %)
HRMS (ESI) calcd for $[C_{62}H_{69}N_{12}O_2]^+$: 1013.56610, found 1013.56422.

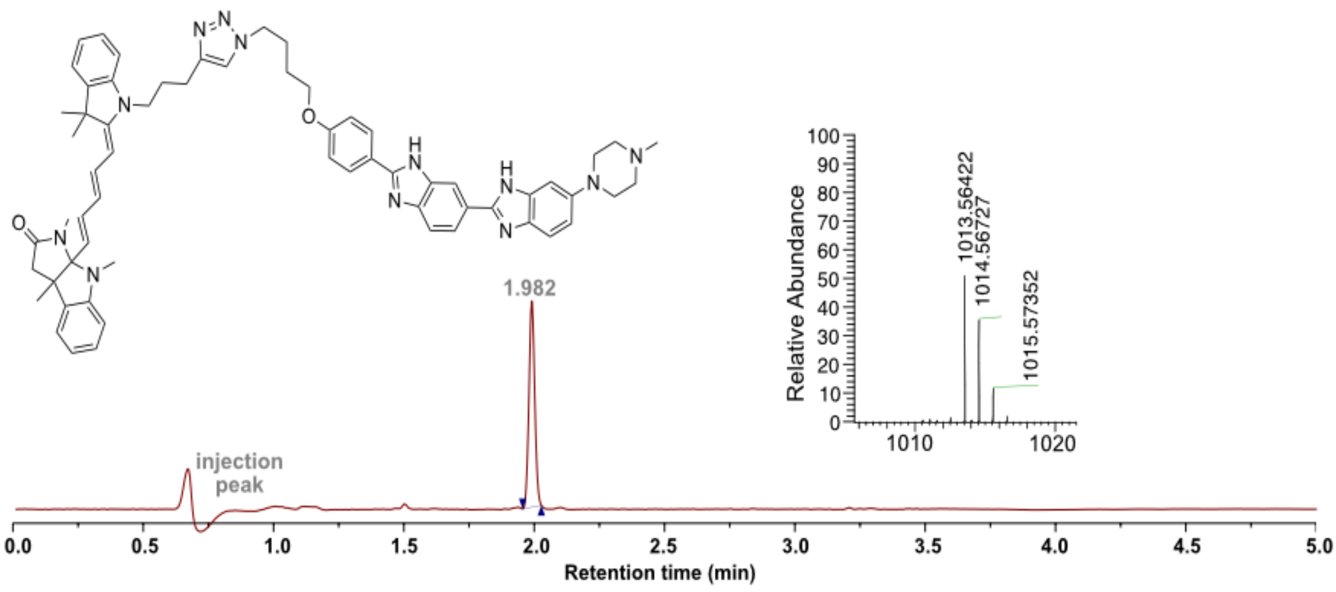

**2-((*E*)-3-((*E*)-3,3-Dimethyl-1-(pent-4-yn-1-yl)-5-(trifluoromethyl)indolin-2-ylidene)prop-1-en-1-yl)-3-(2-methoxy-2-oxoethyl)-1,3-dimethyl-3*H*-indol-1-ium 2,2,2-trifluoro-acetate (21)**

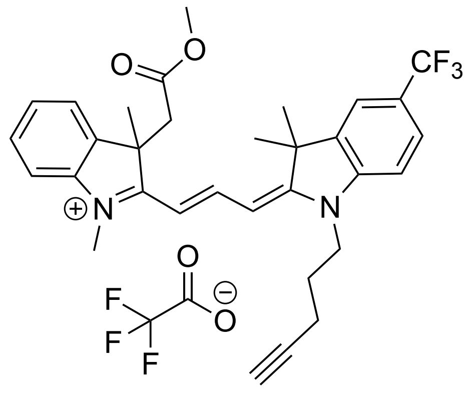

In a flame-dried flask, compound **6** (50 mg, 0.16 mmol, 1.0 equiv.) was dissolved in anhydrous $CH_3CN$ (5 mL) and oxalyl chloride (19 μL, 0.22 mmol, 1.3 equiv.) was added dropwise. The solution was stirred for 1 h at 18 °C. Anhydrous $CH_3OH$ (5 mL) was added and the solution was stirred for additional 15 min. Solvents were removed under reduced pressure and the residue was used directly without further purification. The residue was dissolved in $Ac_2O$ (1 mL) in a microwave vial. *N,N'*-Diphenylformamidine (40 mg, 0.21 mmol, 1.2 equiv.) was added and the solution was heated to 95 °C for 1 h in the microwave. The mixture was cooled to room temperature and a solution of **7d** (74 mg, 0.18 mmol, 1.1 equiv.) in pyridine (1 mL) was added dropwise. The mixture was heated in the microwave at 100 °C for 30 min. Solvents were removed under reduced pressure and the residue was purified via reverse-phase column chromatography (10 $\rightarrow$ 60% $CH_3CN$ in $ddH_2O$ + 0.1% TFA over 35 min) to afford the product as a fine red powder (72 mg, 67%).

$^1H$ NMR (400 MHz, $CD_3OD$): δ = 8.45 (t, *J* = 13.5 Hz, 1H), 7.85 (s, 1H), 7.74 (d, *J* = 9.4 Hz, 1H), 7.59 (d, *J* = 7.4 Hz, 1H), 7.53 – 7.45 (m, 3H), 7.39 – 7.35 (m, 1H), 6.56 (d, *J* = 13.9 Hz, 1H), 6.48 (d, *J* = 13.1 Hz, 1H), 4.24 (t, *J* = 7.4 Hz, 2H), 3.79 (s, 3H), 3.65 (d, *J* = 16.9 Hz, 1H), 3.42 (s, 3H), 3.36 (d, *J* = 17.1 Hz, 1H), 2.50 (t, *J* = 2.6 Hz, 1H), 2.40 (td, *J* = 6.8, 2.7 Hz, 2H), 2.04 (p, *J* = 6.8 Hz, 2H), 1.81 (s, 3H), 1.80 (s, 3H), 1.73 (s, 3H) ppm.
$^{13}C\{^1H\}$ NMR (101 MHz, $CD_3OD$): δ = 175.5, 173.8, 169.4, 150.2, 145.3, 143.4, 141.2, 139.0, 129.0, 126.4 (q, *J* = 32 Hz), 126.2, 124.4 (q, *J* = 270 Hz), 121.9, 119.4 (q, *J* = 3.7 Hz), 111.7, 110.6, 104.5, 102.2, 82.3, 70.0, 51.0, 50.9, 48.8, 44.0, 42.6, 31.1, 26.9, 26.9, 26.2, 25.7, 15.2 ppm.
HRMS (ESI) calcd for $[C_{32}H_{34}F_3N_2O_2]^+$: 535.25724, found 535.25695.

**2-((*E*)-3-((*E*)-3,3-Dimethyl-1-(pent-4-yn-1-yl)-5-(trifluoromethyl)indolin-2-ylidene)prop-1-en-1-yl)-1,3-dimethyl-3-(2-(methylamino)-2-oxoethyl)-3*H*-indol-1-ium 2,2,2-trifluoro-acetate (22)**

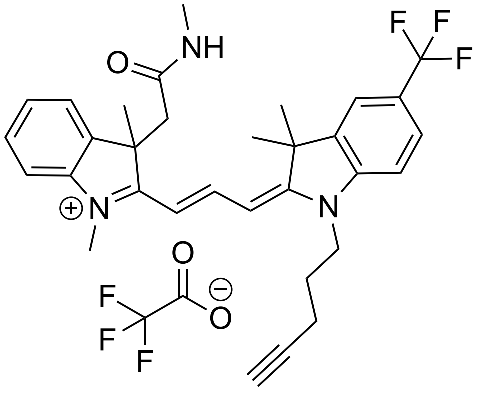

Compound **21** (25 mg, 0.04 mmol, 1.0 equiv.) was dissolved in anhydrous THF (3 mL) and methyl amine (2.0 M solution in THF, 5 mL) was added. The solution was stirred at 70 °C for 16 h. Solvents were removed under reduced pressure and the residue was purified via reverse-phase column chromatography (10 → 50% $CH_3CN$ in dd$H_2O$ + 0.1% TFA over 35 min) to afford the product as a fine red powder (24 mg, 96%).

$^1$H NMR (400 MHz, $CD_3OD$): δ = 8.45 (t, *J* = 13.5 Hz, 1H), 7.82 (s, 1H), 7.72 (d, *J* = 8.4 Hz, 1H), 7.53 (d, *J* = 7.5 Hz, 1H), 7.49 (dd, *J* = 7.1, 1.2 Hz, 1H), 7.47 – 7.43 (m, 2H), 7.37 (td, *J* = 7.3, 1.3 Hz, 1H), 6.57 (d, *J* = 13.9 Hz, 1H), 6.43 (d, *J* = 12.9 Hz, 1H), 4.22 (t, *J* = 7.4 Hz, 2H), 3.80 (s, 3H), 3.39 (d, *J* = 15.9 Hz, 1H), 3.23 (d, *J* = 15.9 Hz, 1H), 2.50 (t, *J* = 2.7 Hz, 1H), 2.43 (s, 3H), 2.39 (td, *J* = 6.8, 2.6 Hz, 2H), 2.03 (p, *J* = 6.9 Hz, 2H), 1.79 (s, 3H), 1.78 (s, 3H), 1.72 (s, 3H) ppm.

$^{13}$C{$^1$H} NMR (101 MHz, $CD_3OD$): δ = 178.4, 174.2, 170.5, 151.4, 146.8, 144.8, 142.4, 140.4, 130.3, 127.6 (q, *J* = 4.1 Hz), 127.53, 127.47 (q, *J* = 32 Hz), 125.8 (q, *J* = 270.7 Hz), 123.3, 120.7 (q, *J* = 3.5 Hz), 113.3, 111.7, 106.7, 103.1, 83.7, 71.4, 52.9, 50.0, 47.1, 43.9, 32.6, 28.5, 28.4, 27.4, 27.0, 26.0, 16.6 ppm.

HRMS (ESI) calcd for $[C_{32}H_{35}F_3N_3O]^+$: 534.27322, found 534.27268.

**3-(Carboxymethyl)-2-((1*E*,3*E*,5*E*)-7-((*E*)-3,3-dimethyl-1-(pent-4-yn-1-yl)indolin-2-ylidene)hepta-1,3,5-trien-1-yl)-1,3-dimethyl-3*H*-indol-1-ium 2,2,2-trifluoro-acetate (23)**

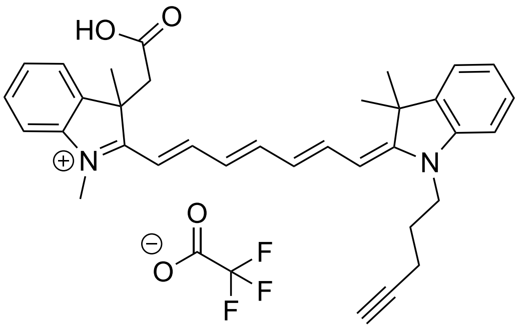

Compound **6** (100 mg, 0.34 mmol, 1.0 equiv.) and glutaconaldehydedianil hydrochloride (114 mg, 0.40 mmol, 1.2 equiv.) were dissolved in $Ac_2O$ (2 mL) in a microwave vial. The solution was heated to 120 °C for 1 h in the microwave. The mixture was cooled to room temperature and a solution of **7c** (130 mg, 0.37 mmol, 1.1 equiv.) in pyridine (2 mL) was added dropwise. The mixture was heated to 100 °C for 30 min. Solvents were removed under reduced pressure and the residue was purified via reverse phase column chromatography (10 → 65% $CH_3CN$ in dd$H_2O$ + 0.1% TFA over 35 min) to afford the product as a fine green powder (95 mg, 46%).

$^1$H NMR (400 MHz, $CD_3OD$): δ = 7.97 (t, *J* = 12.7 Hz, 1H), 7.71 (t, *J* = 12.9 Hz, 1H), 7.51 (d, *J* = 7.5 Hz, 1H), 7.43 (td, *J* = 8.2, 1.3 Hz, 1H), 7.39 – 7.35 (m, 2H), 7.33 – 7.28 (m, 2H), 7.10 (d, *J* = 7.8 Hz, 2H), 6.58 – 6.39 (m, 4H), 6.04 (d, *J* = 13.3 Hz, 1H), 4.03 (t, *J* = 7.4 Hz, 2H), 3.72 (s, 3H), 3.38 (d, *J* = 16.8 Hz, 1H), 3.28 (d, *J* = 16.8 Hz, 1H), 2.47 (t, *J* = 2.7 Hz, 1H), 2.34 (dt, *J* = 6.9, 3.4 Hz, 2H), 1.94 (p, *J* = 6.8 Hz, 2H), 1.65 (s, 6H), 1.58 (s, 3H) ppm.

$^{13}$C NMR (101 MHz, $CD_3OD$) δ = 177.9, 175.6, 168.4, 156.1, 154.4, 148.5, 145.1, 144.4, 142.7, 141.5, 129.6, 129.4, 127.3, 126.2, 124.2, 123.2, 123.0, 112.9, 110.1, 108.7, 101.6, 84.0, 73.7, 71.6, 71.1, 53.5, 42.9, 32.2, 28.4, 27.4, 26.9, 16.6 ppm.

HRMS (ESI) calcd for $[C_{34}H_{37}N_2O_2]^+$: 505.28495, found 505.28459.

**3-(2-((2,2-Difluoroethyl)amino)-2-oxoethyl)-2-((1*E*,3*E*,5*E*)-7-((*E*)-3,3-dimethyl-1-(pent-4-yn-1-yl)indolin-2-ylidene)hepta-1,3,5-trien-1-yl)-1,3-dimethyl-3*H*-indol-1-ium 2,2,2-trifluoro-acetate (24)**

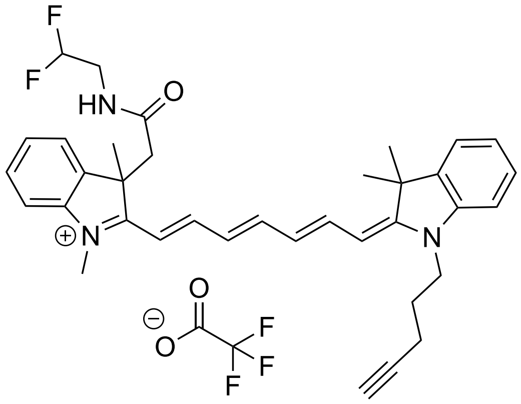

In a flame-dried flask, compound **23** (20 mg, 0.03 mmol, 1.0 equiv.) was dissolved in dry $CH_2Cl_2$ (5 mL) and 4-dimethylaminopyridine (15.8 mg, 0.13 mmol, 4.0 equiv.) and *N*-(3-dimethylamino-propyl)-3-ethyl carbodiimide hydrochloride (EDC-HCl, 24.8 mg, 0.13 mmol, 4.0 equiv.) were added. The solution was stirred at 18 °C for 30 min. 2,2-Difluoroethylamine hydrochloride (76 mg, 0.65 mmol, 20.0 equiv.) was added and the solution was stirred overnight at 18 °C. Solvents were removed under reduced pressure and the residue was purified via reverse phase column chromatography (10 → 75% $CH_3CN$ in dd$H_2O$ + 0.1% TFA over 35 min) to afford the product as a fine green powder (18 mg, 82%).

$^1$H NMR (400 MHz, $CD_3CN$): δ = 7.89 – 7.83 (m, 1H), 7.78 – 7.72 (m, 1H), 7.58 (t, *J* = 13.0 Hz, 1H), 7.47 (d, *J* = 7.9 Hz, 1H), 7.43 – 7.39 (m, 2H), 7.37 – 7.33 (m, 1H), 7.26 (t, *J* = 7.6 Hz, 2H), 7.17 (d, *J* = 7.8 Hz, 2H), 6.53 – 6.42 (m, 2H), 6.30 (d, *J* = 14.2 Hz, 1H), 6.15 (d, *J* = 13.4 Hz, 1H), 5.55 (tt, *J* = 56.0, 3.9 Hz, 1H), 4.02 (t, *J* = 7.5 Hz, 2H), 3.57 (s, 3H), 3.37 (d, *J* = 16.0 Hz, 1H), 3.32 – 3.22 (m, 3H), 2.36 – 2.31 (m, 3H), 1.64 (s, 6H), 1.60 (s, 3H) ppm.

$^{13}$C NMR (101 MHz, $CD_3CN$) δ = 174.5, 171.0, 169.9, 156.6, 152.9, 150.6, 144.8, 143.8, 141.8, 140.6, 129.7, 129.4, 126.6, 126.2, 125.0, 123.2, 117.6, 115.2, 112.8, 112.1, 111.0, 106.3, 103.1, 84.2, 71.1, 68.3, 52.0, 49.5, 46.4, 43.4, 41.8 (t, *J* = 26.7 Hz), 32.3, 28.0, 27.4, 26.7, 16.4 ppm.

HRMS (ESI) calcd for $[C_{36}H_{40}F_2N_3O]^+$: 568.31340, found 568.31309.

**Click reactions**

**General procedure for the click reaction of carbocyanines bearing an alkyne handle (4c, 13, 14, 22, 24) to azide-modified SNAPtag ligand 10**

In a flame-dried flask, alkyne-modified carbocyanine dye (15 mg, 1.0 equiv.) and **10** (1.5 equiv.) were dissolved in degassed THF (5 mL) and the solution was degassed for an additional hour by bubbling argon through the solution while stirring vigorously. $Cu(PPh_3)_2NO_3$ (0.5 equiv.) and triethylamine (10 $\mu$L) were added and the solution was stirred for 16 h at room temperature. Solvents were removed under reduced pressure and the residue was purified via reverse-phase column chromatography (10 $\rightarrow$ 45% $CH_3CN$ in dd$H_2O$ + 0.1% TFA over 35 min).

**2-((1*E*,3*E*)-5-((*E*)-1-(3-(1-(4-(((2-Amino-9*H*-purin-6-yl)oxy)methyl)benzyl)-1*H*-1,2,3-triazol-4-yl)propyl)-3,3-dimethyl-5-(trifluoromethyl)indolin-2-ylidene)penta-1,3-dien-1-yl)-3-(2-hydroxyethyl)-1,3-dimethyl-3*H*-indol-1-ium 2,2,2-trifluoroacetate (11)**

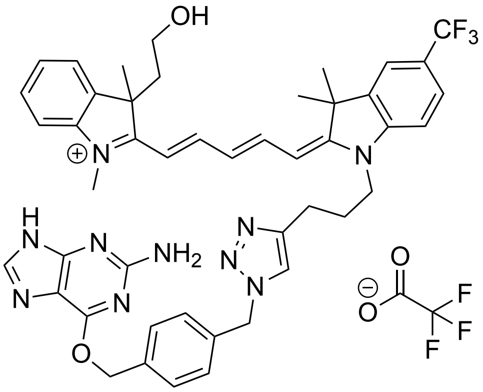

Yield: 10 mg, 45%
$^1$H NMR (400 MHz, $CD_3OD$): $\delta$ = 8.36 – 8.29 (m, 1H), 8.26 – 8.151 (m, 2H), 7.83 (s, 1H), 7.71 (s, 1H), 7.60 – 7.54 (m, 4H), 7.52 – 7.48 (m, 1H), 7.44 – 7.37 (m, 4H), 7.19 (d, *J* = 8.4 Hz, 1H), 6.60 (t, *J* = 12.4 Hz, 1H), 6.49 (d, *J* = 14.2 Hz, 1H), 6.13 (d, *J* = 13.1 Hz, 1H), 5.62 (s, 2H), 5.61 (s, 2H), 4.07 (t, *J* = 7.6 Hz, 2H), 3.73 (s, 3H), 3.13 – 3.00 (m, 2H), 2.86 (t, *J* = 7.3 Hz, 2H), 2.79 – 2.72 (m, 1H), 2.55 – 2.47 (m, 1H), 2.21 – 2.13 (m, 2H), 1.74 (s, 3H), 1.72 (s, 6H) ppm.
$^{13}$C{$^1$H} NMR (126 MHz, $CD_3OD$): $\delta$ = 177.6, 171.4, 161.1, 158.9, 156.7, 153.6, 148.1, 147.1, 144.6, 142.7, 141.0, 137.4, 137.2, 130.4, 130.1, 129.4, 129.2, 127.8, 127.6, 127.3 (q, *J* = 4.2 Hz), 125.9 (q, *J* =270.8 Hz), 123.8, 123.7, 120.4 (q, *J* = 3.3 Hz), 113.1, 110.9, 107.9, 103.4, 70.1, 59.2, 54.5, 54.1, 44.5, 43.9, 36.5, 32.3, 28.0, 27.9, 27.3, 23.4, 14.4 ppm.
HRMS (ESI) calcd for $[C_{46}H_{48}F_3N_{10}O_2]^+$: 829.39083, found 829.38980.

**2-((1*E*,3*E*)-5-((*E*)-1-(3-(1-(4-(((2-Amino-9*H*-purin-6-yl)oxy)methyl)benzyl)-1*H*-1,2,3-triazol-4-yl)propyl)-3,3-dimethylindolin-2-ylidene)penta-1,3-dien-1-yl)-1,3-dimethyl-3-(2-(methylamino)-2-oxoethyl)-3*H*-indol-1-ium 2,2,2-trifluoroacetate (15)**

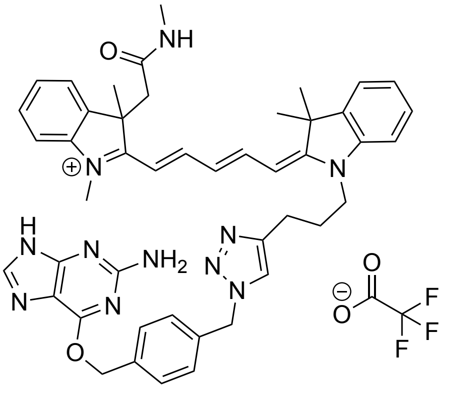

Yield: 14 mg, 64%

$^{1}$H NMR (400 MHz, $CD_3OD$): δ = 8.30 (s, 1H), 8.16 (td, *J* = 13.0, 12.6, 5.9 Hz, 2H), 7.83 (s, 1H), 7.57 (d, *J* = 8.3 Hz, 2H), 7.45 – 7.37 (m, 5H), 7.31 (dd, *J* = 8.4, 3.2 Hz, 2H), 7.26 (t, *J* = 8.0 Hz, 1H), 7.19 (t, *J* = 7.5 Hz, 1H), 7.14 (d, *J* = 7.8 Hz, 1H), 6.52 (t, *J* = 12.4 Hz, 1H), 6.32 (d, *J* = 14.0 Hz, 1H), 6.13 (d, *J* = 13.5 Hz, 1H), 5.63 (s, 2H), 5.61 (s, 2H), 4.10 (t, *J* = 7.6 Hz, 2H), 3.66 (s, 3H), 3.40 (d, *J* = 15.5 Hz, 1H), 3.20 (d, *J* = 15.8 Hz, 1H), 2.86 (t, *J* = 7.3 Hz, 2H), 2.43 (s, 3H), 2.18 (p, *J* = 7.3 Hz, 2H), 1.69 (s, 6H), 1.67 (s, 3H) ppm.

$^{13}$C{$^{1}$H} NMR (151 MHz, $CD_3OD$): δ = 175.6, 173.4, 171.4, 161.1, 158.6, 155.2, 154.5, 148.1, 145.1, 143.7, 142.3, 140.4, 137.5, 130.5, 130.0, 129.6, 129.4, 126.5, 126.3, 125.7, 123.8, 123.4, 123.2, 112.2, 111.5, 105.5, 103.5, 70.3, 54.5, 52.3, 50.19, 46.73, 44.0, 31.8, 28.04, 28.02, 27.6, 27.5, 26.0, 23.4 ppm.

HRMS (ESI) calcd for $[C_{46}H_{50}N_{11}O_2]^+$: 788.41435, found 788.41445.

**2-((1*E*,3*E*)-5-((*E*)-1-(3-(1-(4-(((2-Amino-9*H*-purin-6-yl)oxy)methyl)benzyl)-1*H*-1,2,3-triazol-4-yl)propyl)-3,3-dimethylindolin-2-ylidene)penta-1,3-dien-1-yl)-3,3-dimethyl-1-(2-(methylamino)-2-oxoethyl)-3*H*-indol-1-ium 2,2,2-trifluoroacetate (16)**

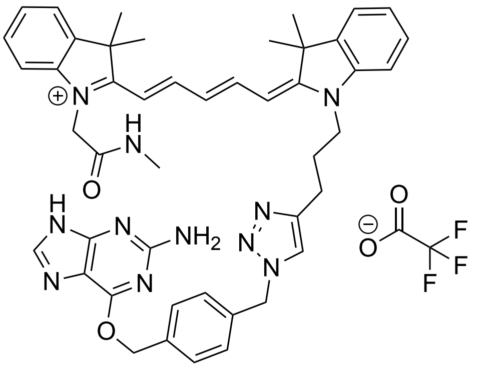

Yield: 9 mg, 42%

$^1$H NMR (400 MHz, $CD_3OD$): δ = 8.32 – 8.17 (m, 2H), 8.13 (s, 1H), 7.83 (s, 1H), 7.56 (d, *J* = 8.4 Hz, 2H), 7.48 (dd, *J* = 14.6, 6.7 Hz, 2H), 7.37 (dd, *J* = 9.5, 8.1 Hz, 4H), 7.29 – 7.23 (m, 3H), 7.14 (d, *J* = 7.9 Hz, 1H), 6.46 (t, *J* = 12.4 Hz, 1H), 6.23 (d, *J* = 13.5 Hz, 1H), 6.07 (d, *J* = 13.9 Hz, 1H), 5.61( s, 2H), 5.60 (s, 2H), 4.78 (s, 2H), 4.18 (t, *J* = 7.5 Hz, 2H), 2.87 (t, *J* = 7.1 Hz, 2H), 2.80 (s, 3H), 2.20 (t, *J* = 7.5 Hz, 2H), 1.75 (s, 6H), 1.70 (s, 6H) ppm.

$^{13}$C{$^1$H} NMR (151 MHz, $CD_3OD$): δ = 175.9, 175.1, 168.4, 161.1, 156.4, 155.2, 148.0, 144.1, 143.3, 142.9, 142.1, 137.5, 137.3, 130.3, 129.8, 129.6, 129.3, 127.0, 126.8, 126.0, 123.9, 123.5, 123.4, 112.3, 111.2, 105.4, 104.0, 69.7, 54.6, 50.9, 50.4, 49.6, 47.3, 44.4, 28.0, 27.8, 27.7, 26.6, 23.3 ppm.

HRMS (ESI) calcd for $[C_{46}H_{50}N_{11}O_2]^+$: 788.41435, found 788.41441.

**2-((*E*)-3-((*E*)-1-(3-(1-(4-(((2-Amino-9*H*-purin-6-yl)oxy)methyl)benzyl)-1*H*-1,2,3-triazol-4-yl)propyl)-3,3-dimethyl-5-(trifluoromethyl)indolin-2-ylidene)prop-1-en-1-yl)-1,3-di-methyl-3-(2-(methylamino)-2-oxoethyl)-3*H*-indol-1-ium 2,2,2-tri-fluoroacetate (19)**

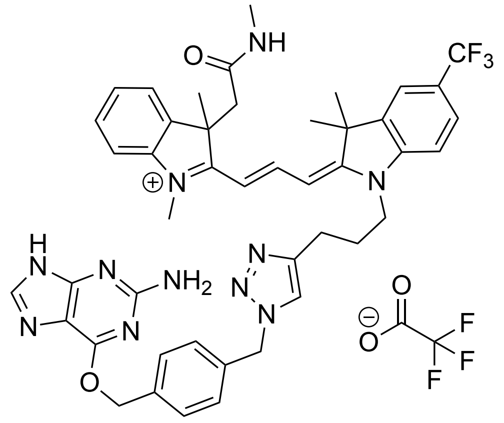

Yield: 16 mg, 73%

$^{1}$H NMR (400 MHz, $CD_3OD$): δ = 8.42 (t, *J* = 13.5 Hz, 1H), 8.31 (s, 1H), 7.84 (s, 1H), 7.80 (s, 1H), 7.66 (d, *J* = 8.4 Hz, 1H), 7.57 (d, *J* = 8.1 Hz, 2H), 7.52 (d, *J* = 7.5 Hz, 1H), 7.50 – 7.42 (m, 2H), 7.40 – 7.28 (m, 4H), 6.57 (d, *J* = 13.9 Hz, 1H), 6.37 (d, *J* = 13.0 Hz, 1H), 5.64 (s, 2H), 5.61 (s, 2H), 4.16 (t, *J* = 7.5 Hz, 2H), 3.78 (s, 3H), 3.38 (d, *J* = 15.9 Hz, 1H), 3.22 (d, *J* = 15.7 Hz, 1H), 2.88 (t, *J* = 7.3 Hz, 2H), 2.42 (s, 3H), 2.21 (p, *J* = 7.5 Hz, 2H), 1.76 (s, 3H), 1.76 (s, 3H), 1.70 (s, 3H) ppm.

$^{13}$C{$^{1}$H} NMR (151 MHz, $CD_3OD$): δ = 178.3, 174.1, 170.5, 161.0, 158.5, 153.9, 151.3, 148.0, 146.8, 144.8, 143.3, 142.5, 140.4, 137.5, 137.1, 130.5, 130.3, 129.4, 127.5, 127.3, 125.8 (q, *J* = 271.3 Hz), 123.9, 123.3, 120.7 (q, *J* = 4.1 Hz), 113.3, 111.7, 106.8, 103.4, 70.3, 54.5, 52.9, 49.9, 47.1, 44.3, 32.6, 28.5, 28.4, 27.43, 27.35, 26.0, 23.4 ppm.

HRMS (ESI) calcd for $[C_{45}H_{47}F_3N_{11}O_2]^+$: 830.38608, found 830.38619.

**2-((1*E*,3*E*,5*E*)-7-((*E*)-1-(3-(1-(4-(((2-Amino-9*H*-purin-6-yl)oxy)methyl)benzyl)-1*H*-1,2,3-triazol-4-yl)propyl)-3,3-dimethylindolin-2-ylidene)hepta-1,3,5-trien-1-yl)-3-(2-((2,2-difluoroethyl)amino)-2-oxoethyl)-1,3-dimethyl-3*H*-indol-1-ium 2,2,2-trifluoroacetate (20)**

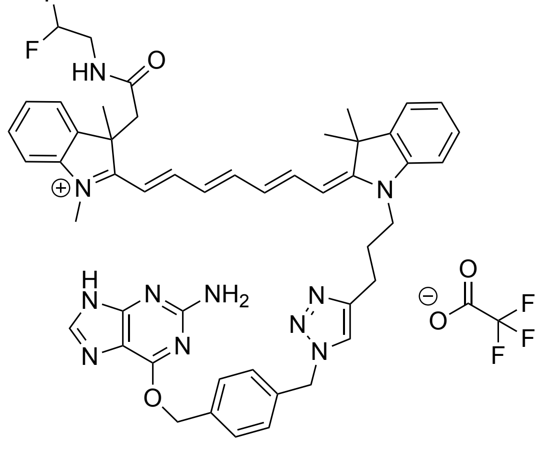

Yield: 12.5 mg, 58%

$^{1}$H NMR (400 MHz, $CD_3CN$): δ = 7.96 (s, 1H), 7.79 – 7.66 (m, 2H), 7.61 (s, 1H), 7.53 (d, *J* = 8.1 Hz, 2H), 7.45 – 7.38 (m, 3H), 7.34 – 7.30 (m, 3H), 7.26 – 7.21 (m, 2H), 7.17 (t, *J* = 7.6 Hz, 1H), 7.11 (d, *J* = 7.9 Hz, 1H), 6.76 (t, *J* = 6.0 Hz, 1H), 6.42 (t, *J* = 12.6 Hz, 1H), 6.29 (dd, *J* = 31.8, 13.3 Hz, 2H), 6.04 (d, *J* = 13.5 Hz, 1H), 5.53 (tt, *J* = 55.9, 3.8 Hz, 1H), 5.56 (s, 2H), 5.55 (s, 2H), 4.00 (t, *J* = 7.7 Hz, 2H), 3.55 (s, 3H), 3.31 – 3.17 (m, 2H), 3.13 – 3.07 (m, 2H), 2.80 (t, *J* = 7.3 Hz, 2H), 2.15 – 2.07 (m, 2H), 1.59 (s, 9H) ppm.

$^{13}$C{$^{1}$H} NMR (126 MHz, $CD_3CN$): δ = 173.5, 171.6, 170.0, 160.9, 158.0, 156.2, 154.2, 152.1, 151.0, 147.8, 144.9, 143.6, 141.9, 140.2, 137.5, 136.9, 130.1, 129.7, 129.4, 129.1, 126.4, 125.9, 125.3, 123.2, 123.1, 117.0, 115.1, 113.2, 111.9, 111.3, 105.7, 103.8, 69.6, 54.0, 51.8, 49.7, 47.2, 46.5, 44.1, 41.8 (t, *J* = 20.8 Hz), 32.3, 27.9, 27.4, 27.2, 23.2, 9.0 ppm.

HRMS (ESI) calcd for $[C_{49}H_{52}F_2N_{11}O_2]^+$: 864.42680, found 864.42562.

### NMR spectra

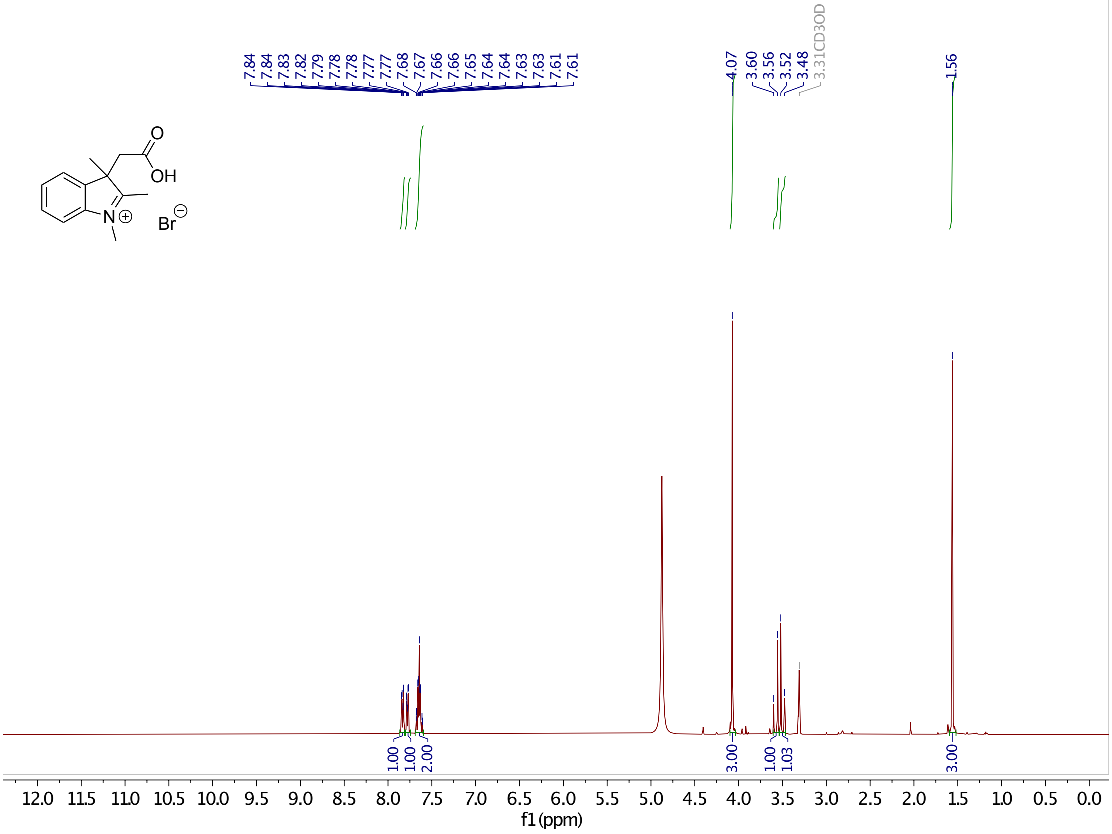

**Supplementary Figure 1.** $^{1}$H-NMR of **6.**

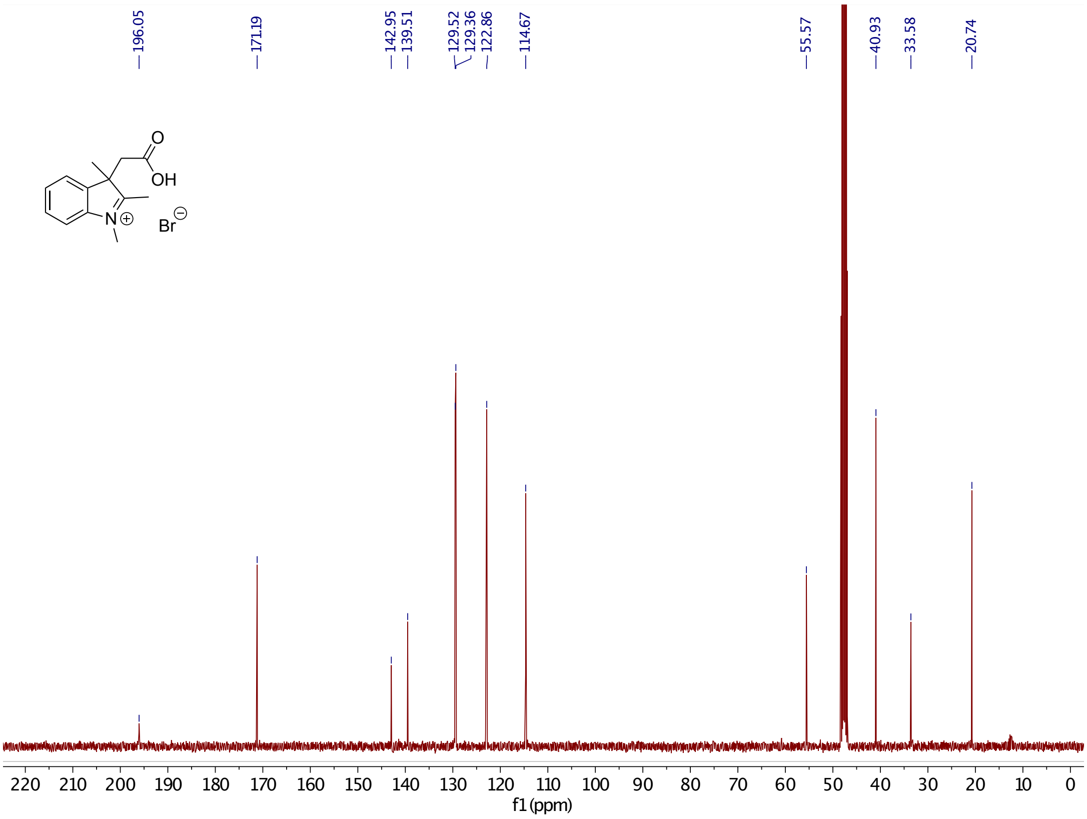

**Supplementary Figure 2.** $^{13}$C-NMR of **6**.

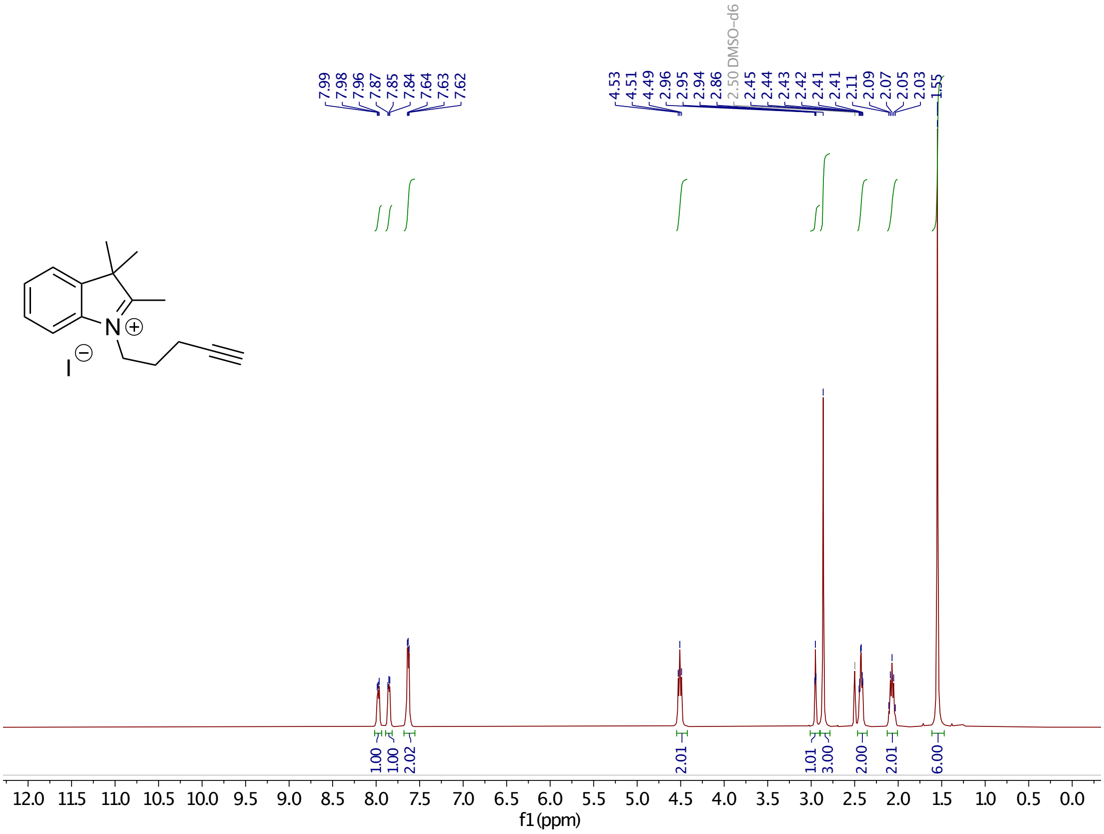

**Supplementary Figure 3.** $^{1}$H-NMR of **7c**.

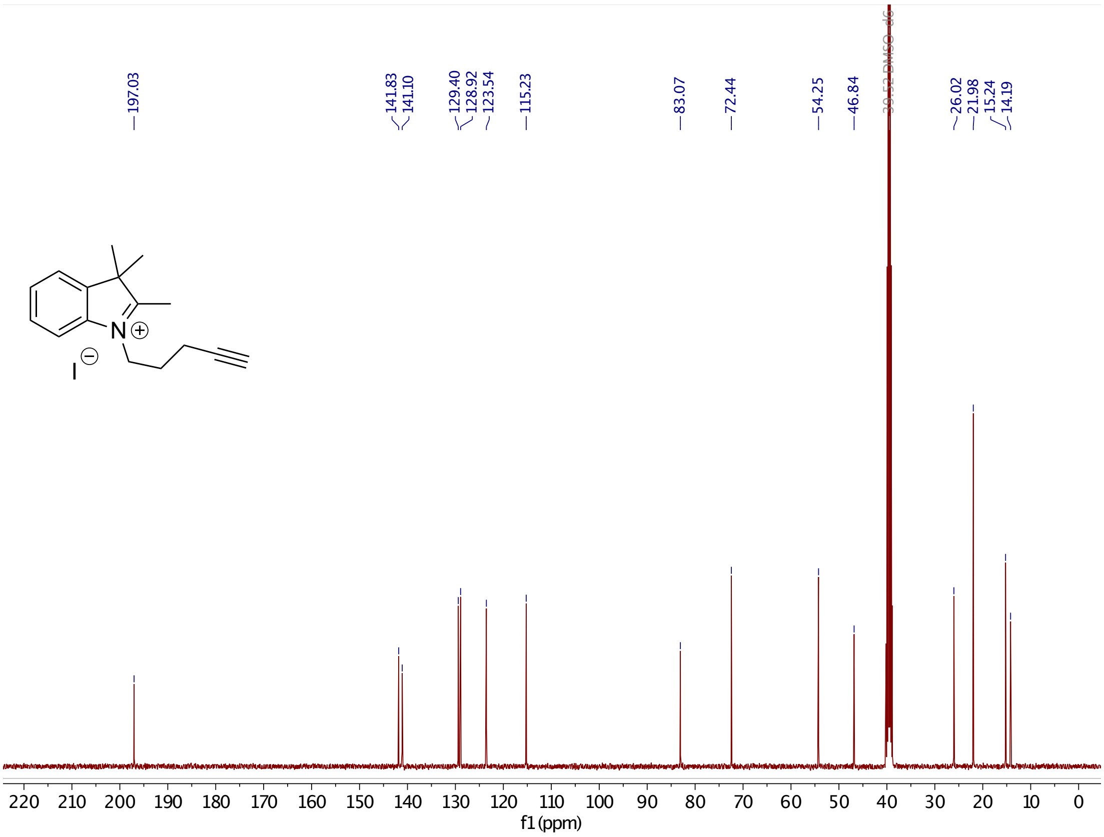

**Supplementary Figure 4.** $^{13}$C-NMR of **7c**.

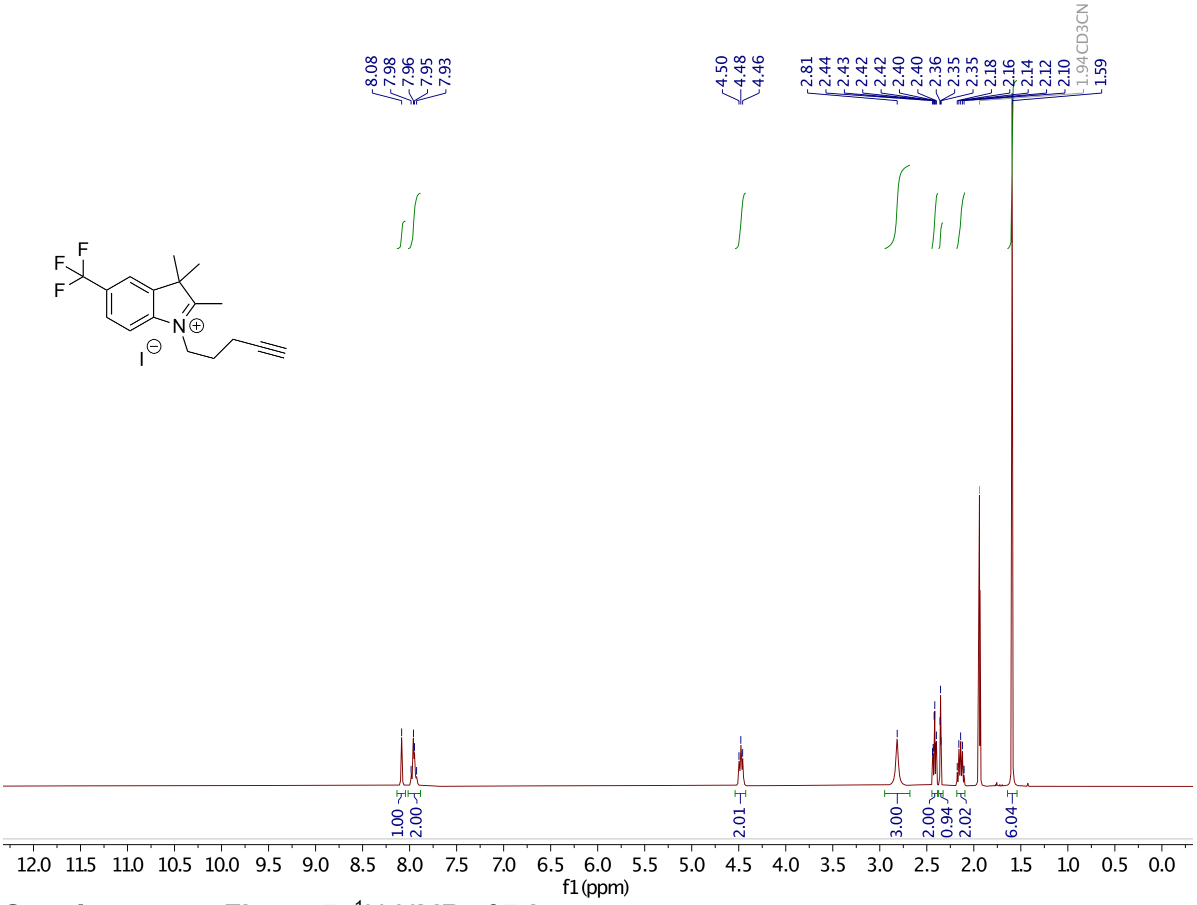

**Supplementary Figure 5.** $^{1}$H-NMR of **7d**.

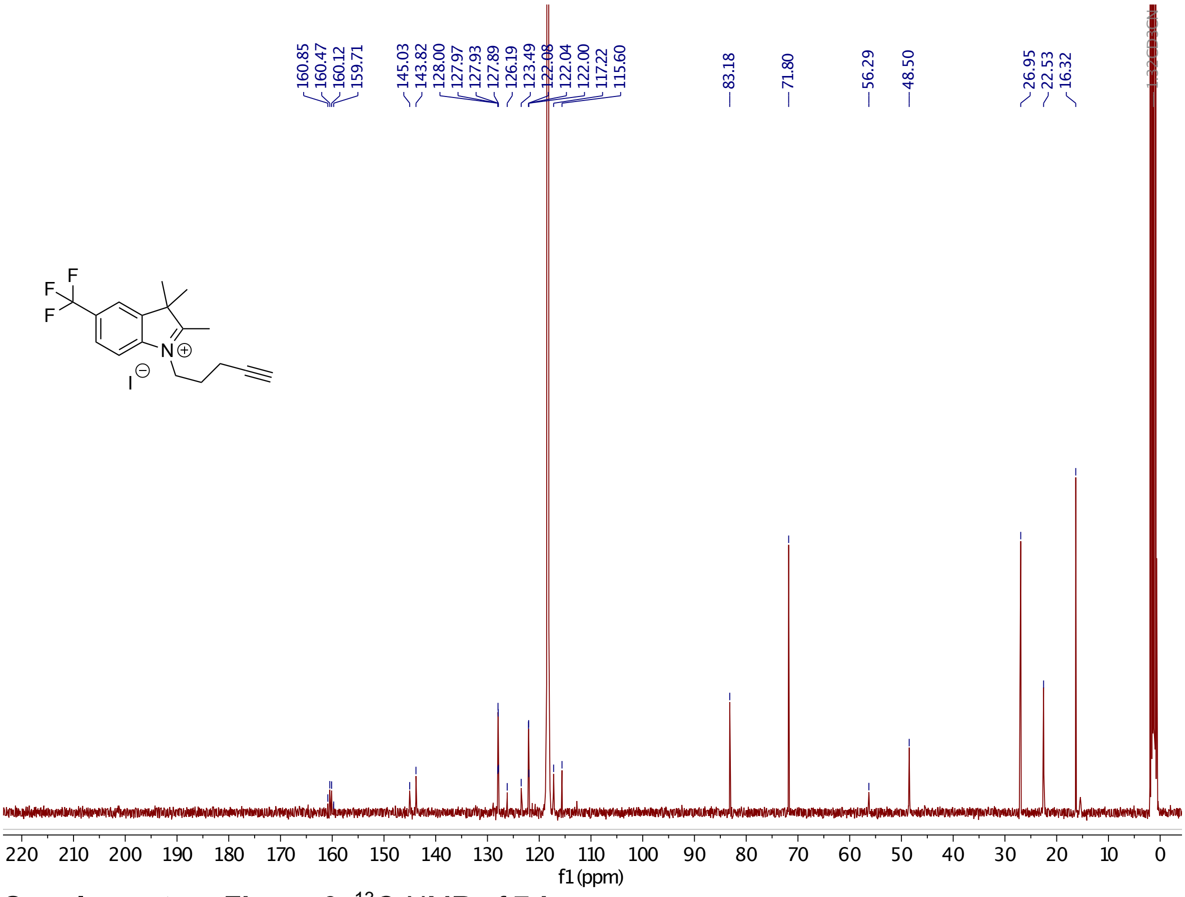

**Supplementary Figure 6.** $^{13}$C-NMR of **7d**.

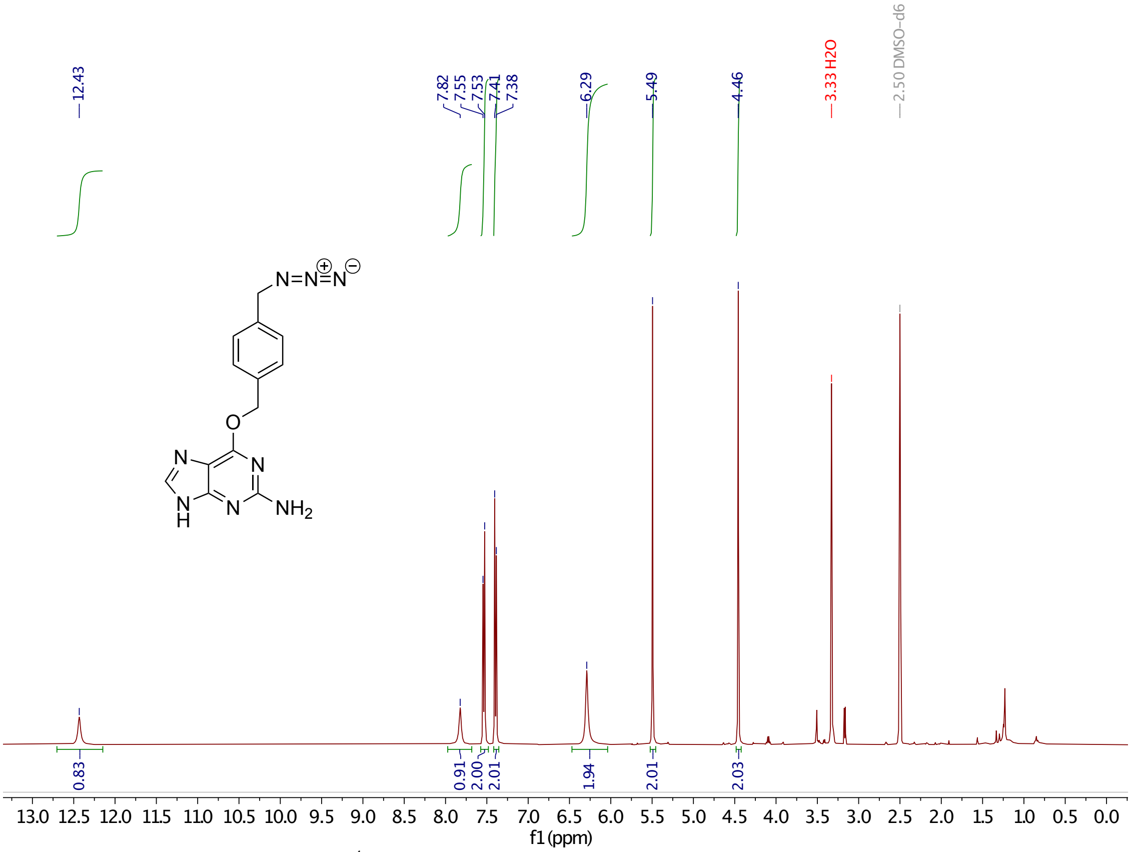

**Supplementary Figure 7.** $^{1}$H-NMR of **10**.

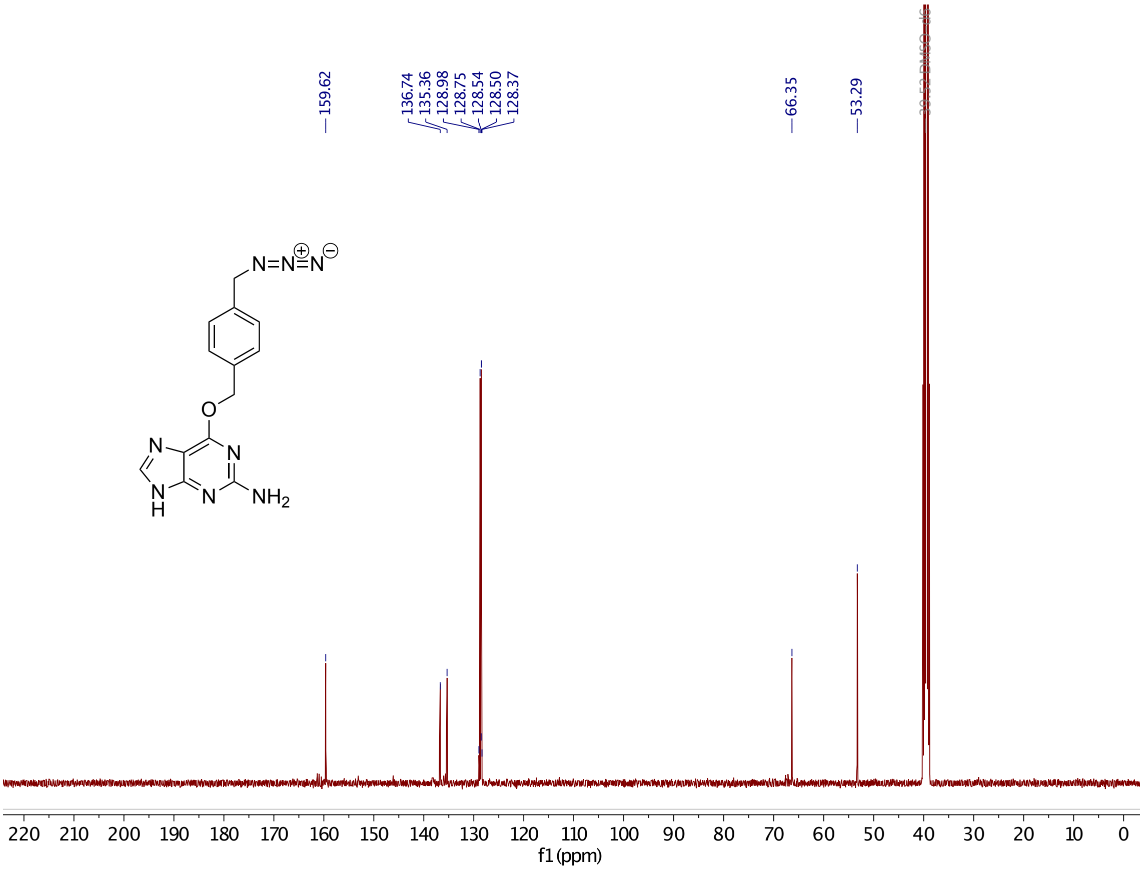

**Supplementary Figure 8.** $^{13}$C-NMR of **10**.

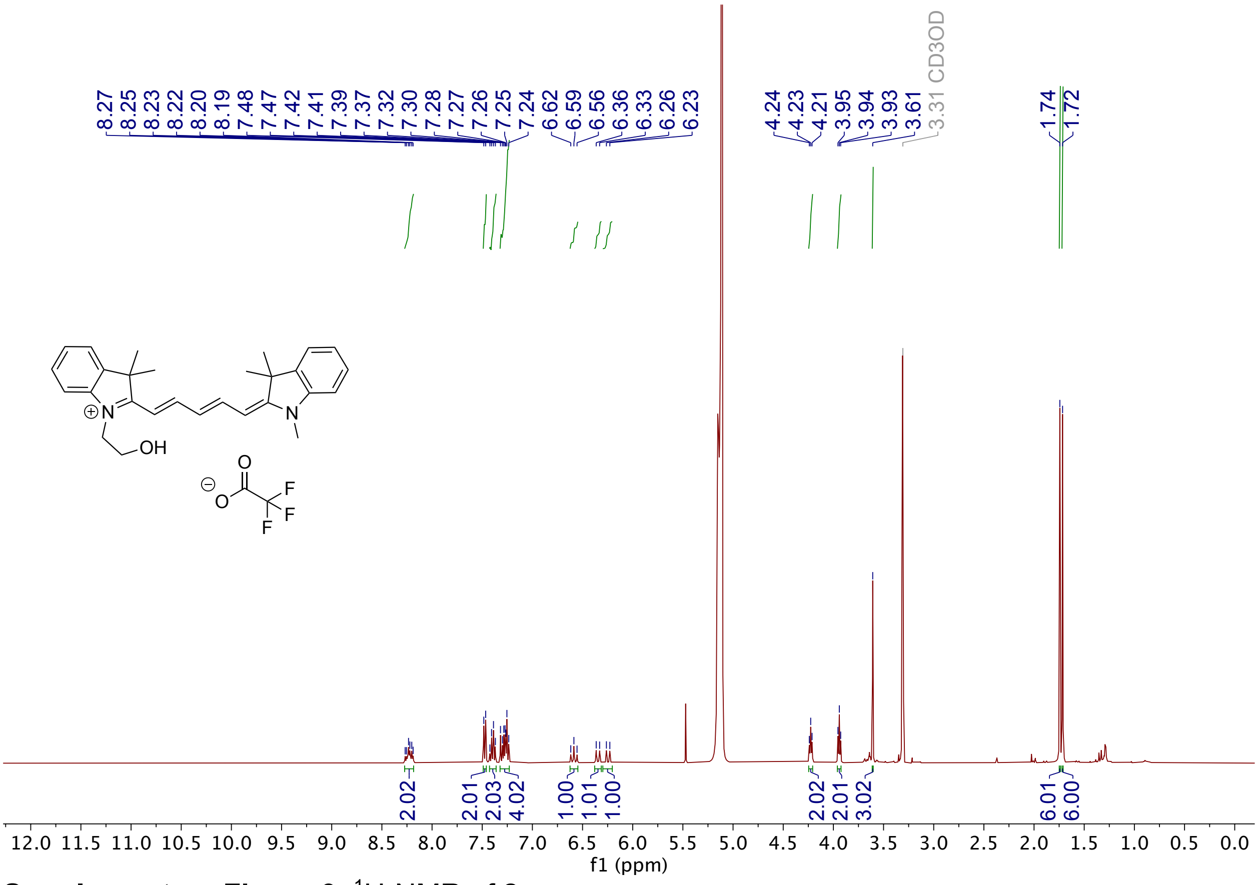

**Supplementary Figure 9.** $^{1}$H-NMR of **2a**.

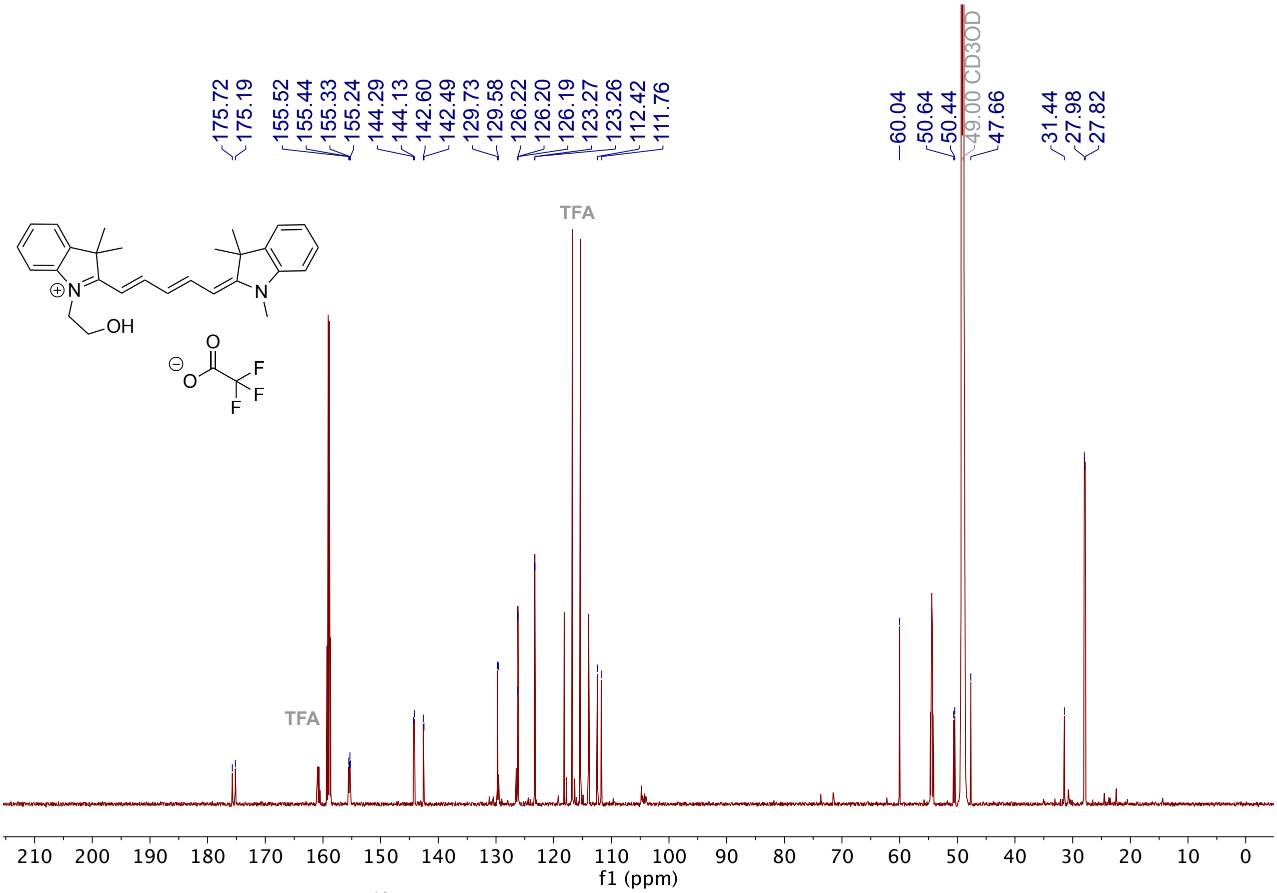

**Supplementary Figure 10.** $^{13}$C-NMR of **2a**.

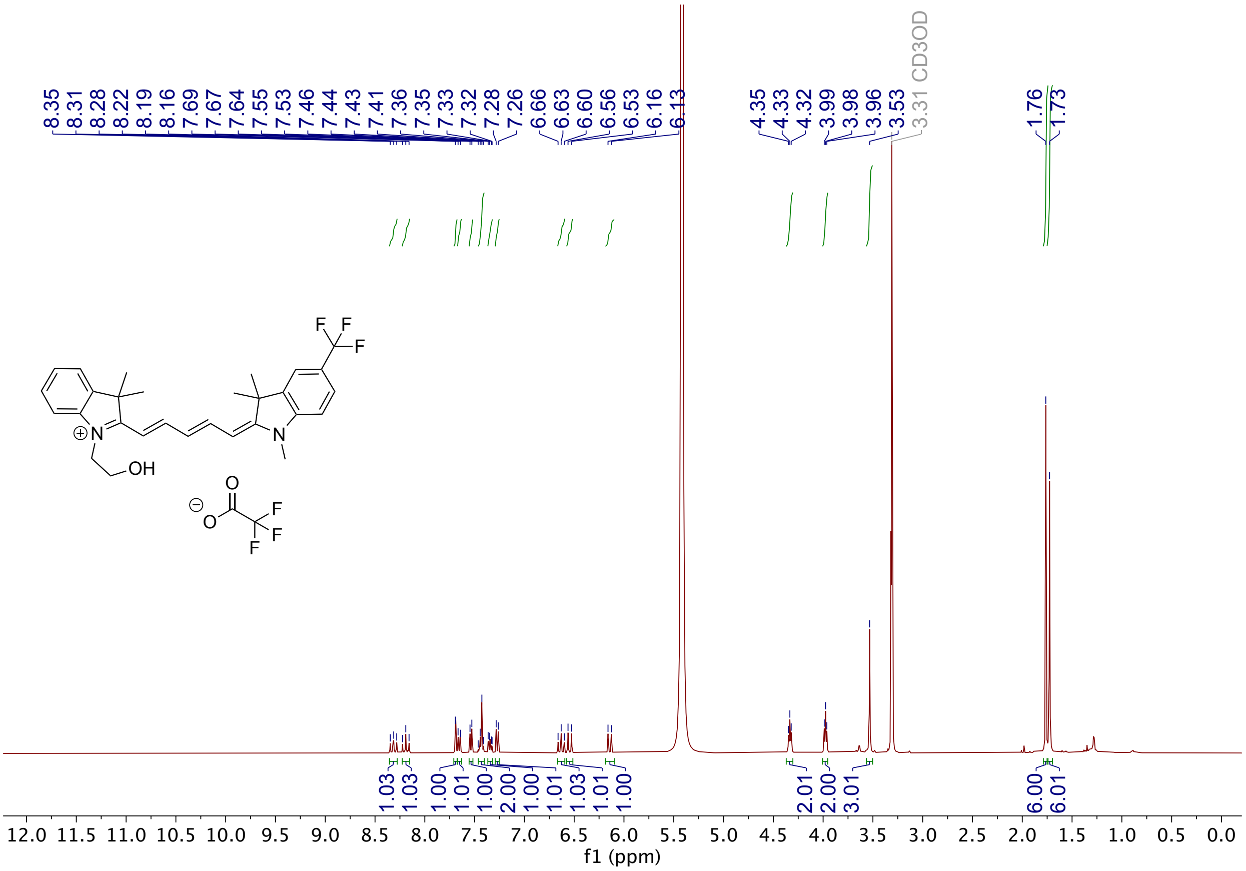

**Supplementary Figure 11.** $^{1}$H-NMR of **2b**.

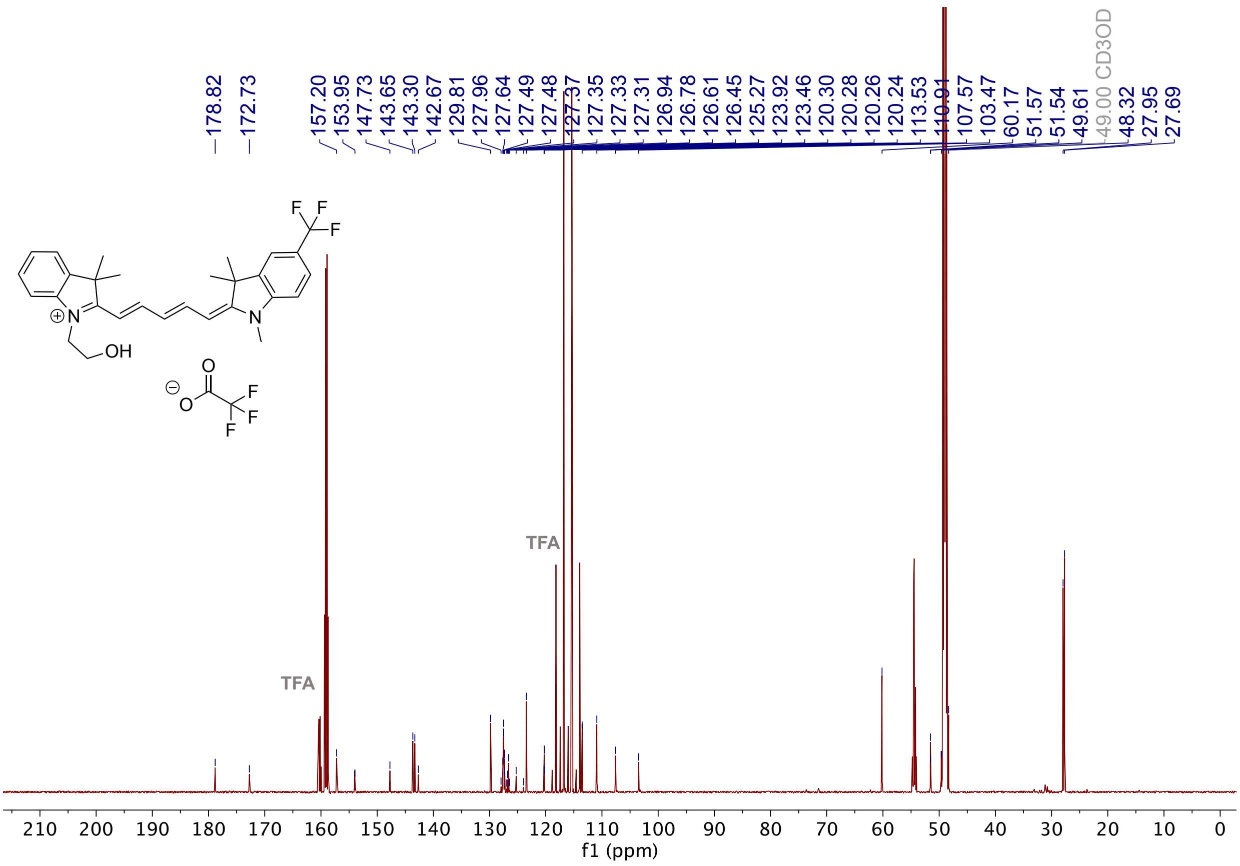

**Supplementary Figure 12.** $^{13}$C-NMR of **2b**.

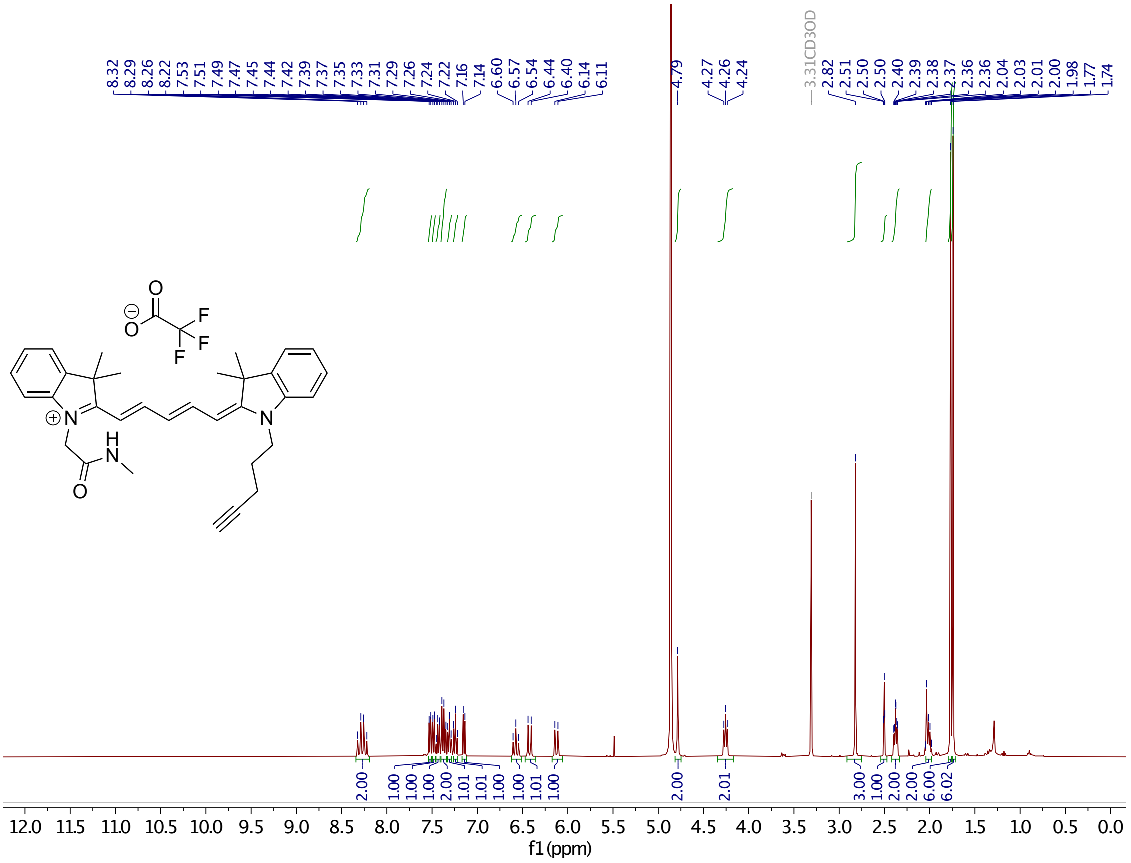

**Supplementary Figure 13.** $^{1}$H-NMR of **14**.

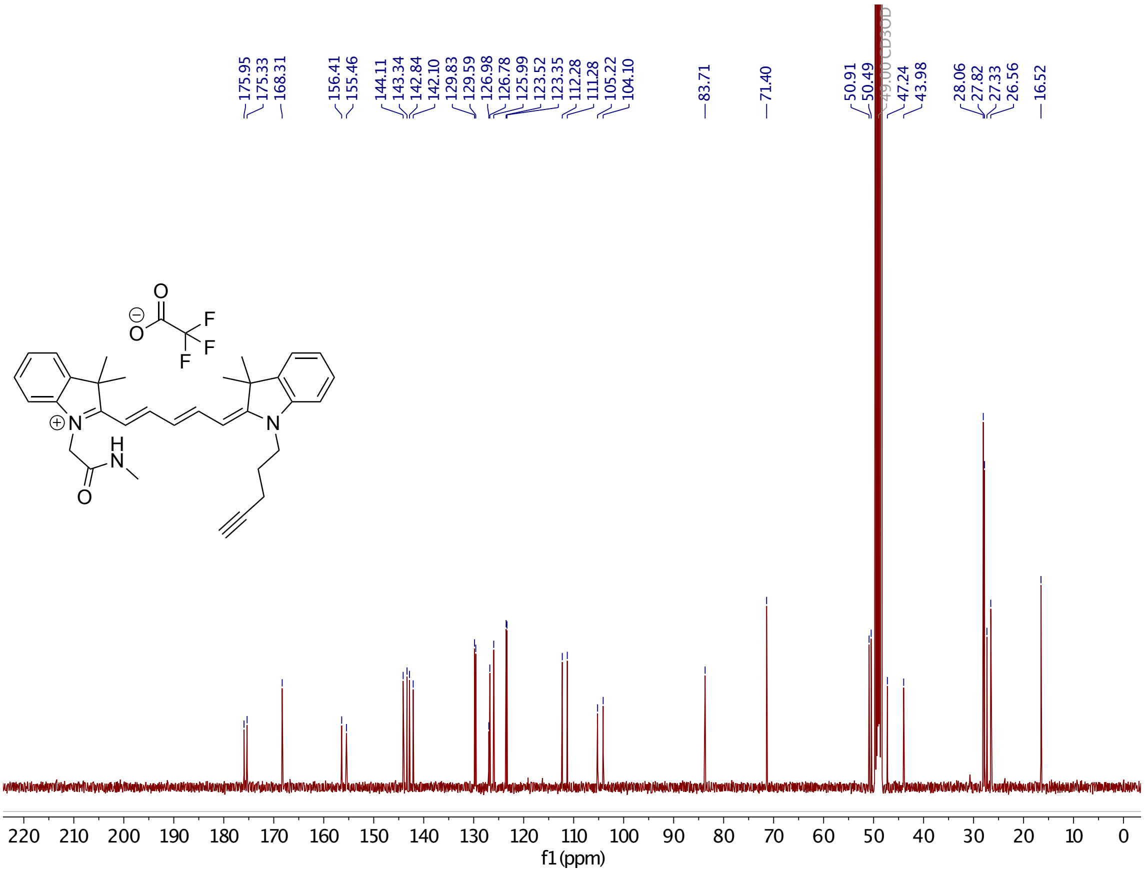

**Supplementary Figure 14**. $^{13}$C-NMR of **14**.

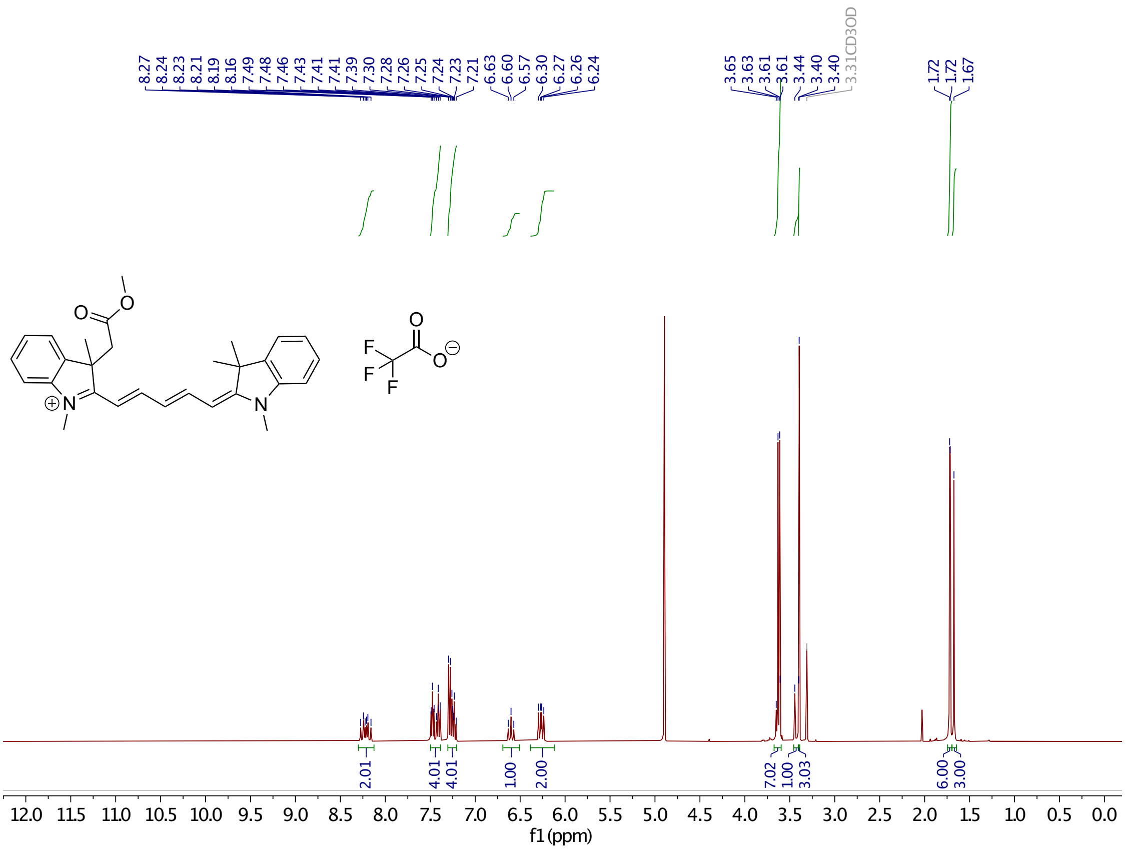

**Supplementary Figure 15.** $^{1}$H-NMR of **9a**.

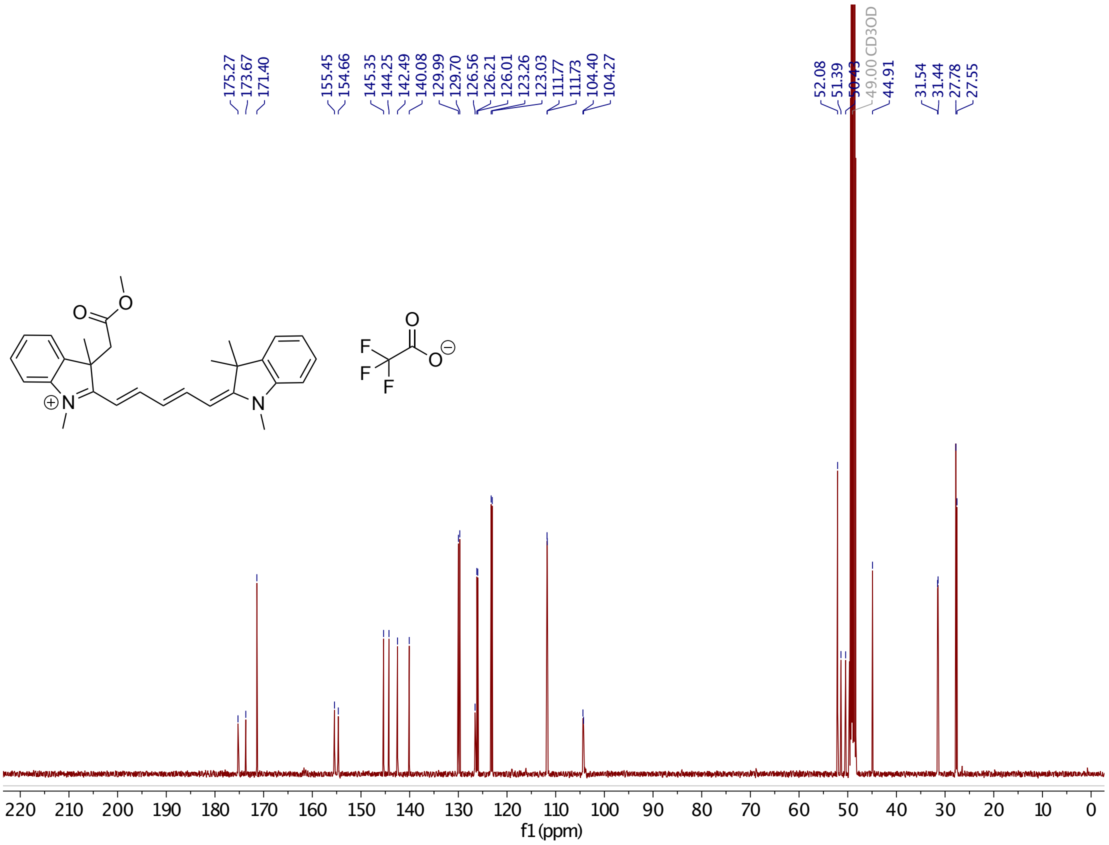

**Supplementary Figure 16.** $^{13}$C-NMR of **9a**.

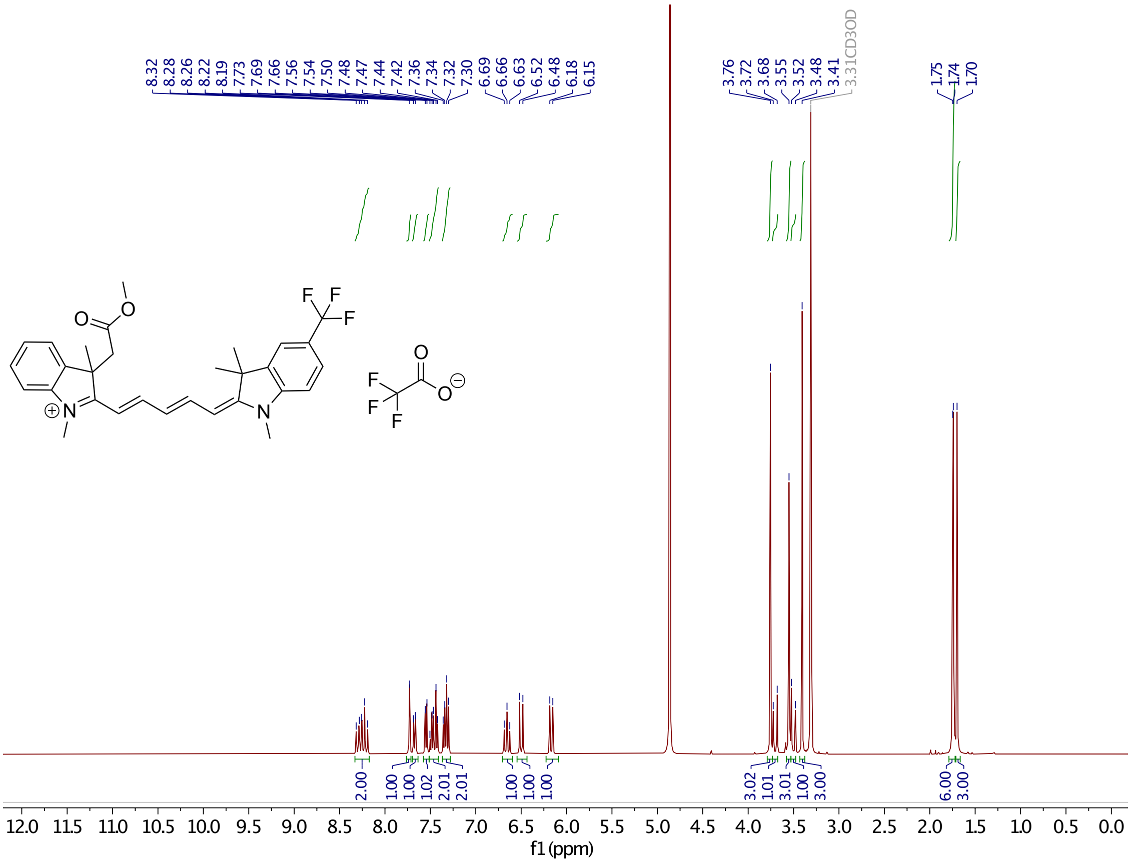

**Supplementary Figure 17.** $^{1}$H-NMR of **9b**

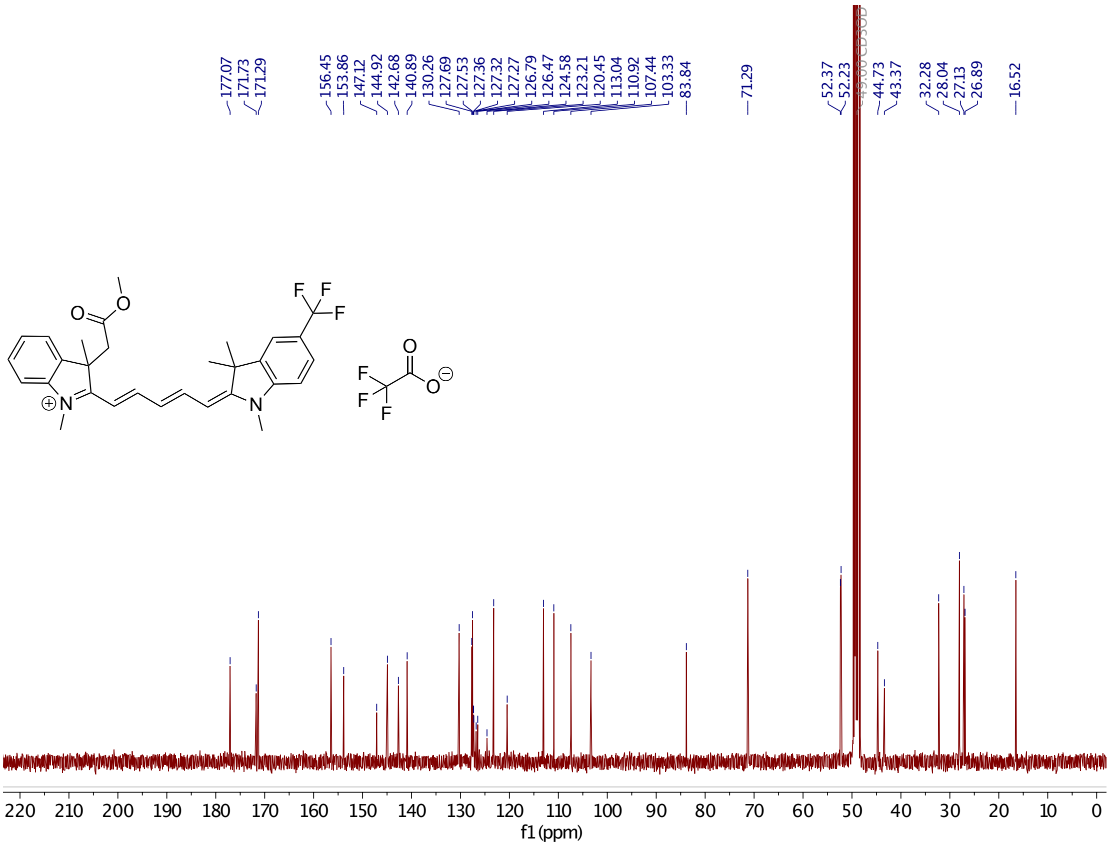

**Supplementary Figure 18.** $^{13}$C-NMR of **9b**.

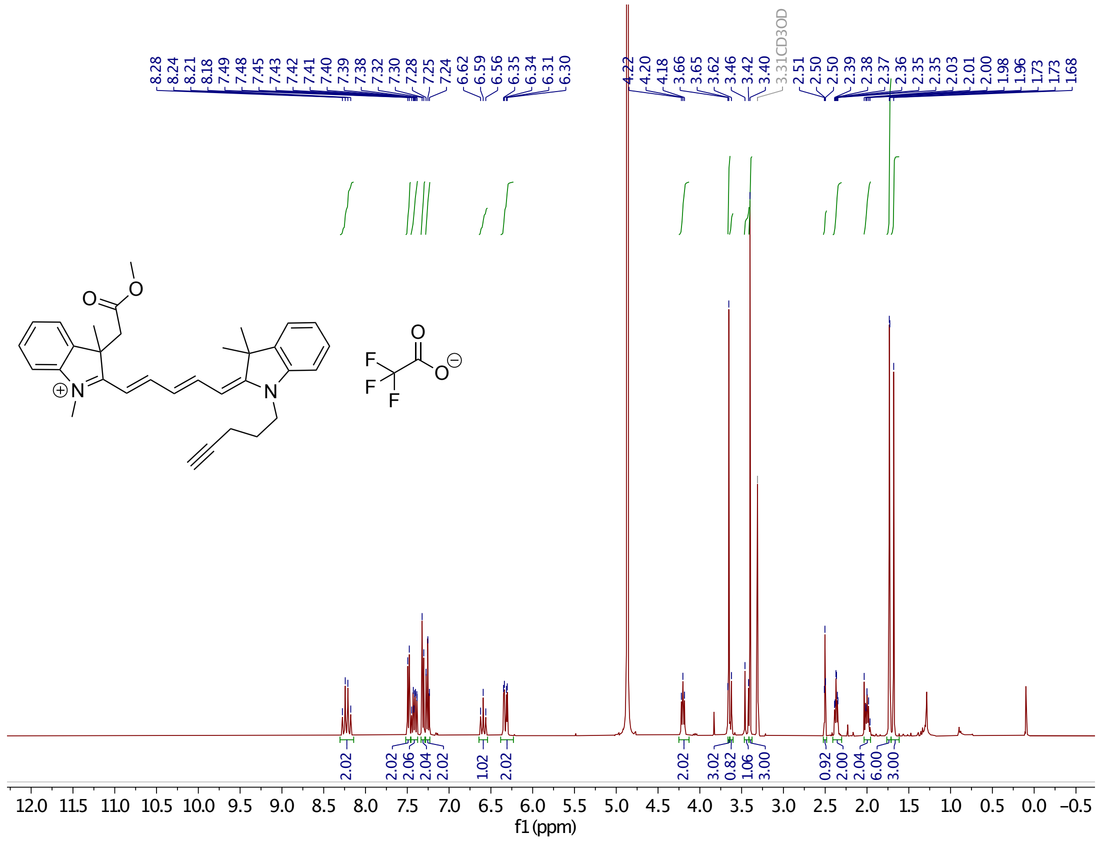

**Supplementary Figure 19.** $^{1}$H-NMR of **9c**.

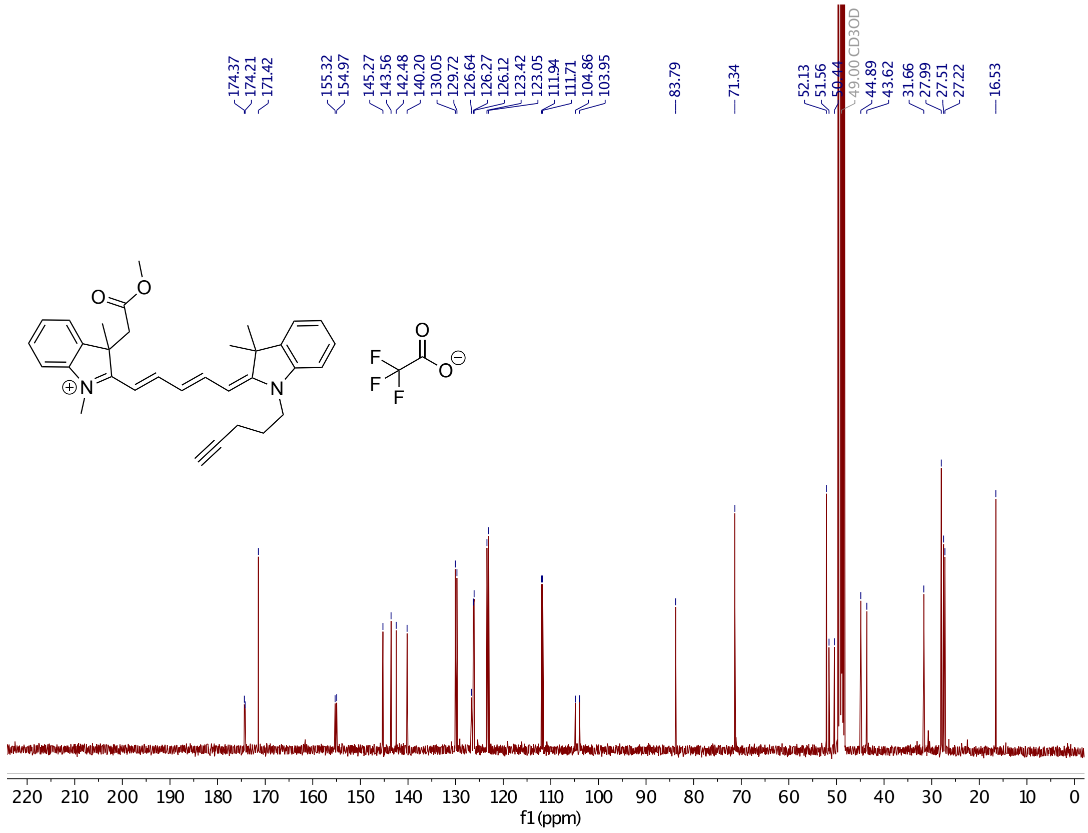

**Supplementary Figure 20.** $^{13}$C-NMR of **9c**.

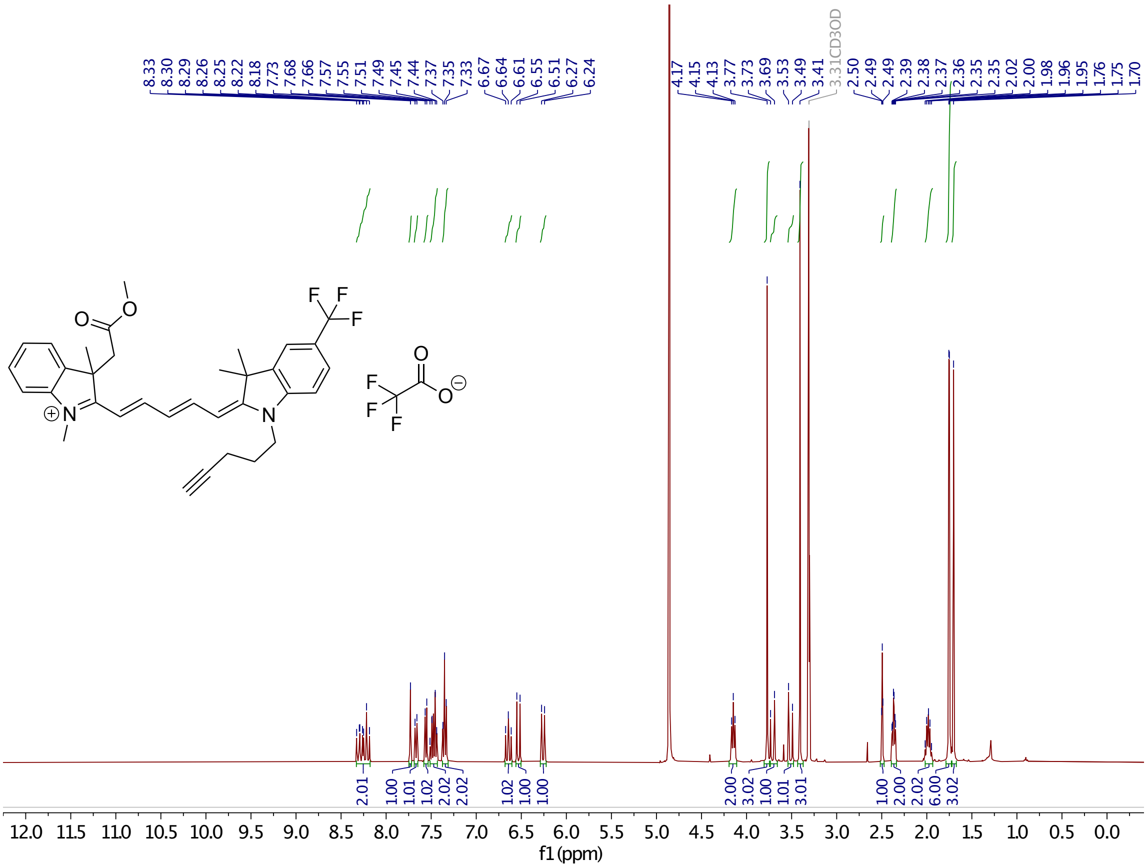

**Supplementary Figure 21.** $^{1}$H-NMR of **9d**.

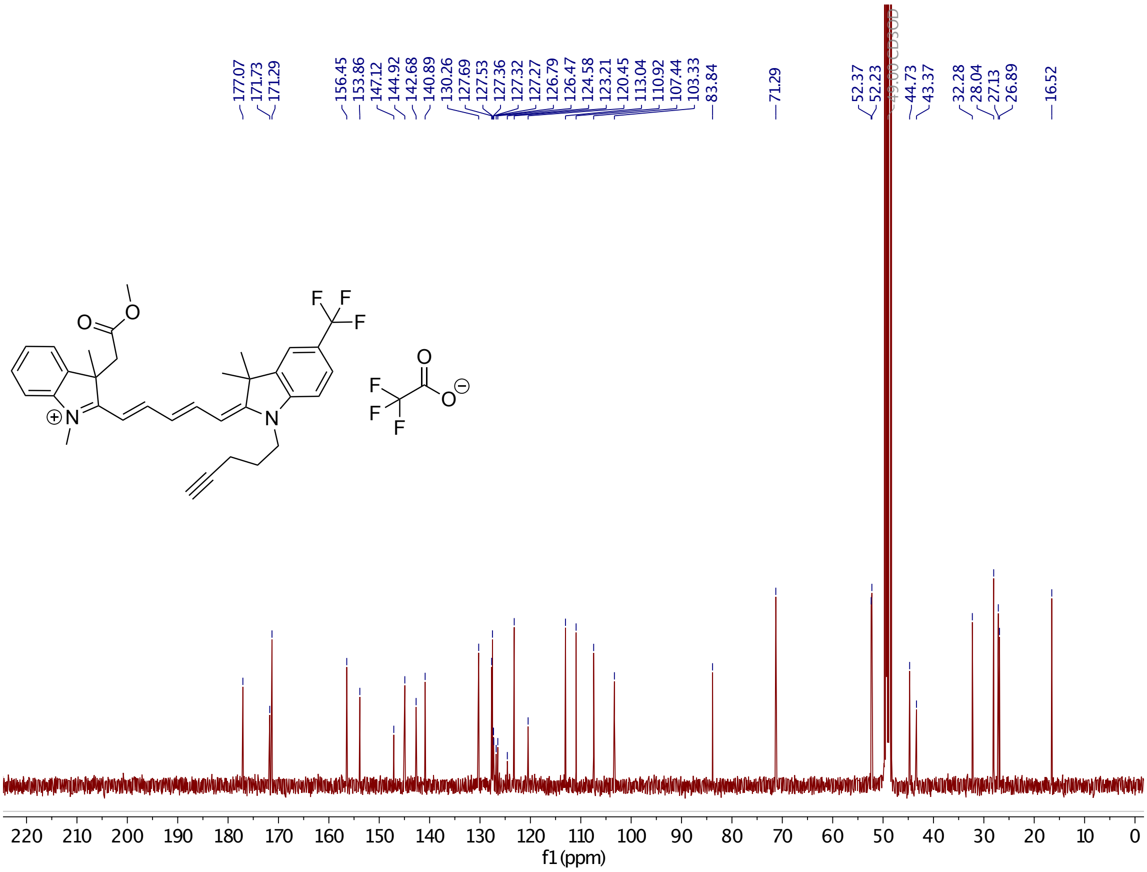

**Supplementary Figure 22.** $^{13}$C-NMR of **9d**.

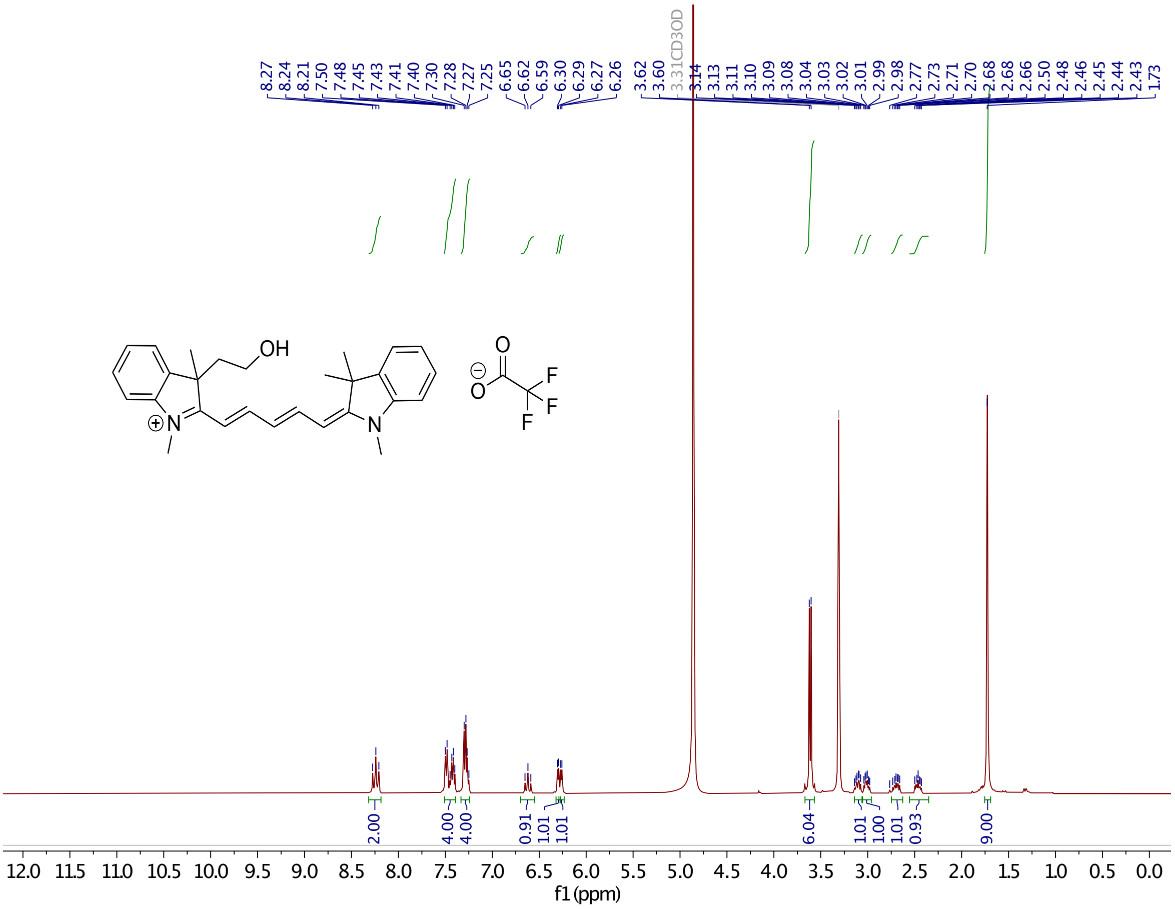

**Supplementary Figure 23.** $^{1}$H-NMR o **4a.**

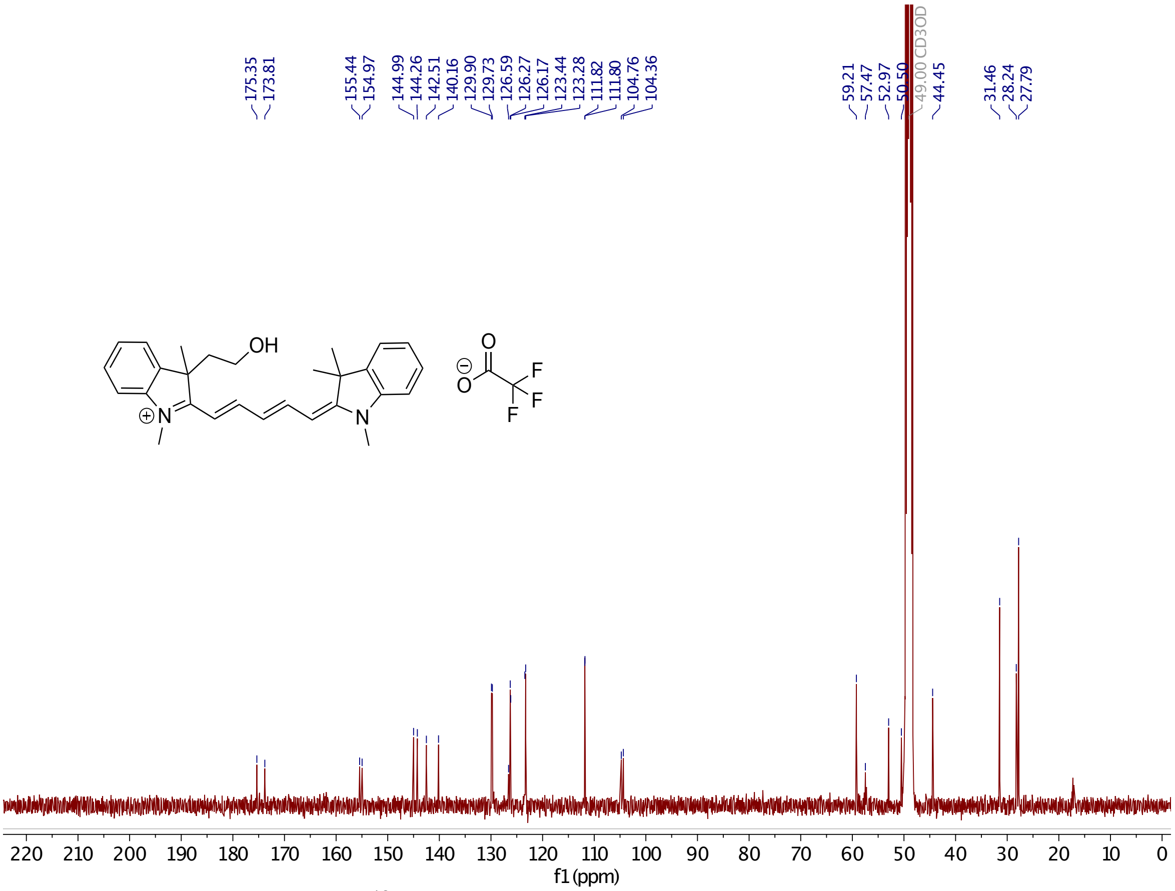

**Supplementary Figure 24.** $^{13}$C-NMR of **4a**.

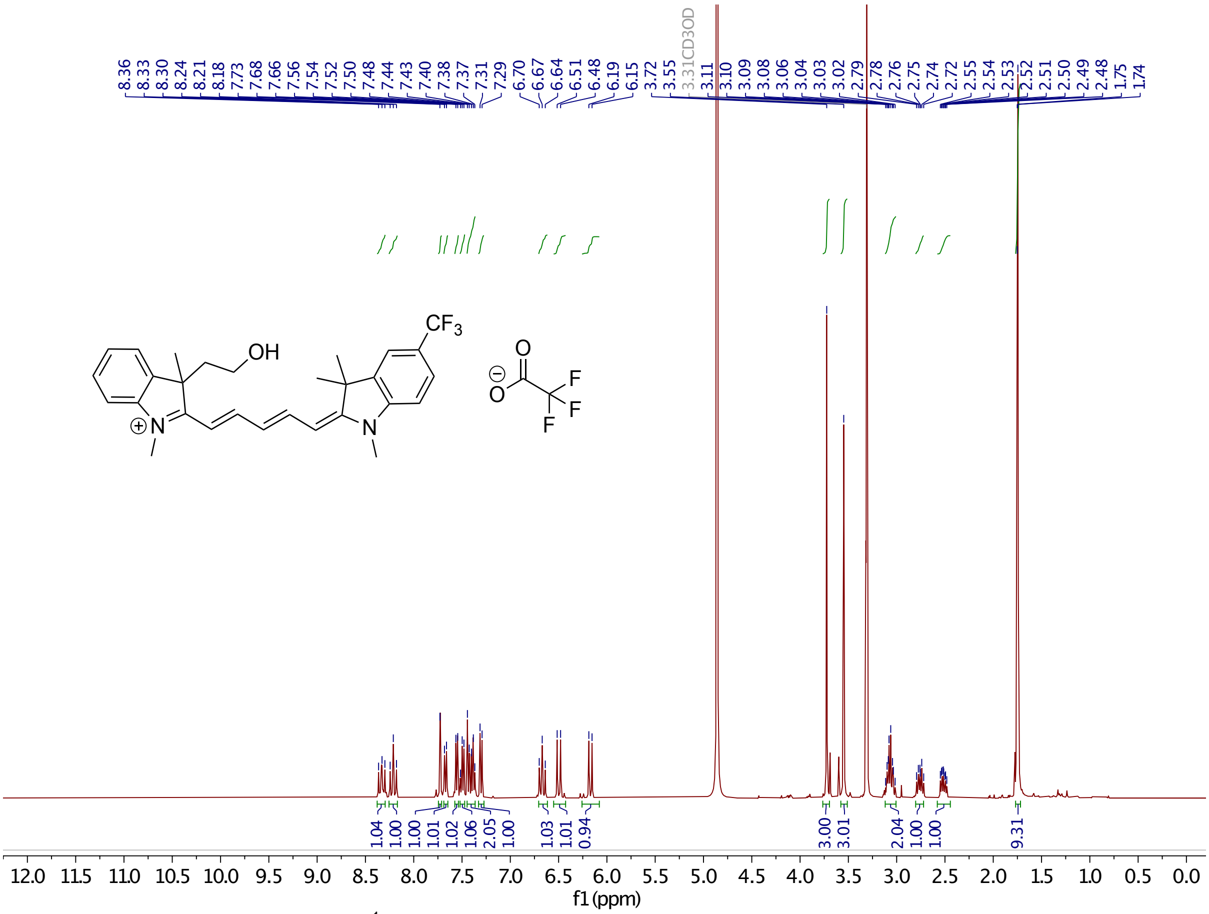

**Supplementary Figure 25.** $^{1}$H-NMR of **4b.**

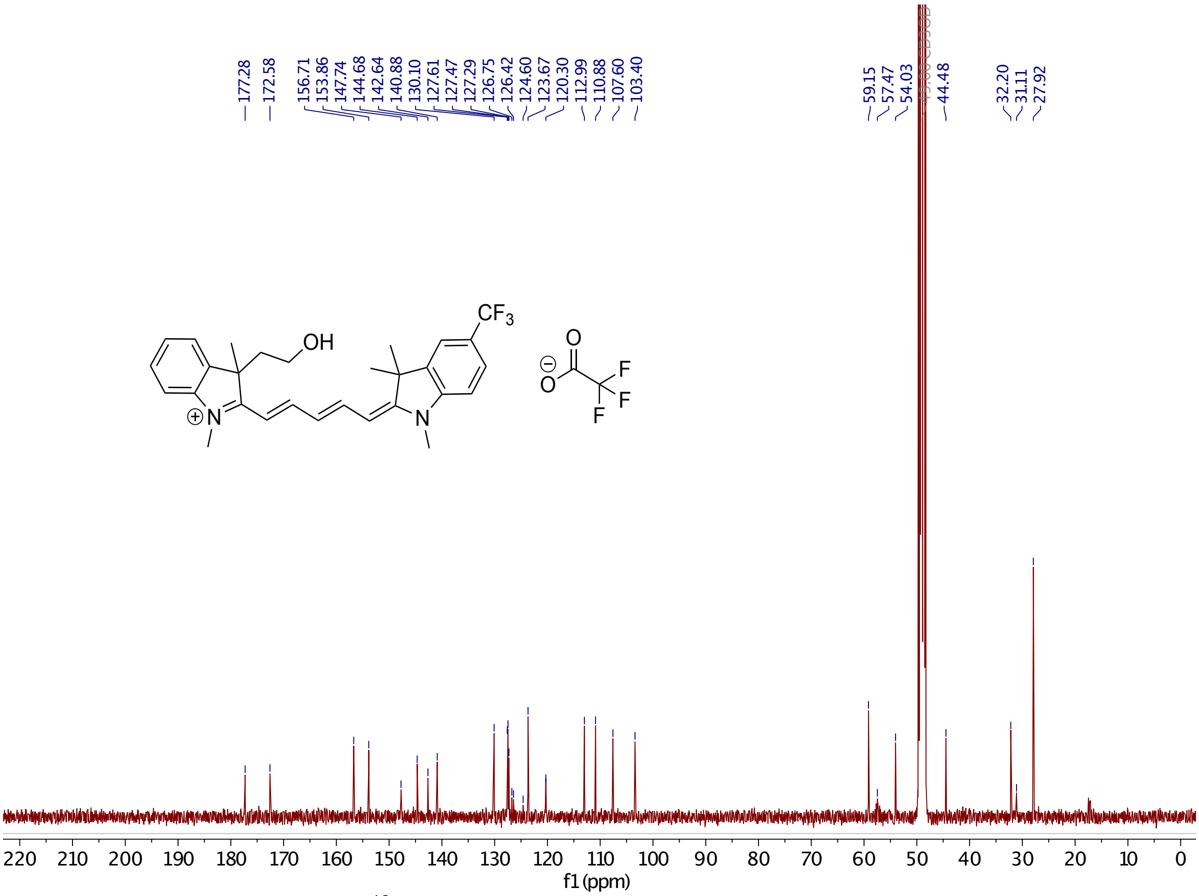

**Supplementary Figure 26.** $^{13}$C-NMR of **4b**.

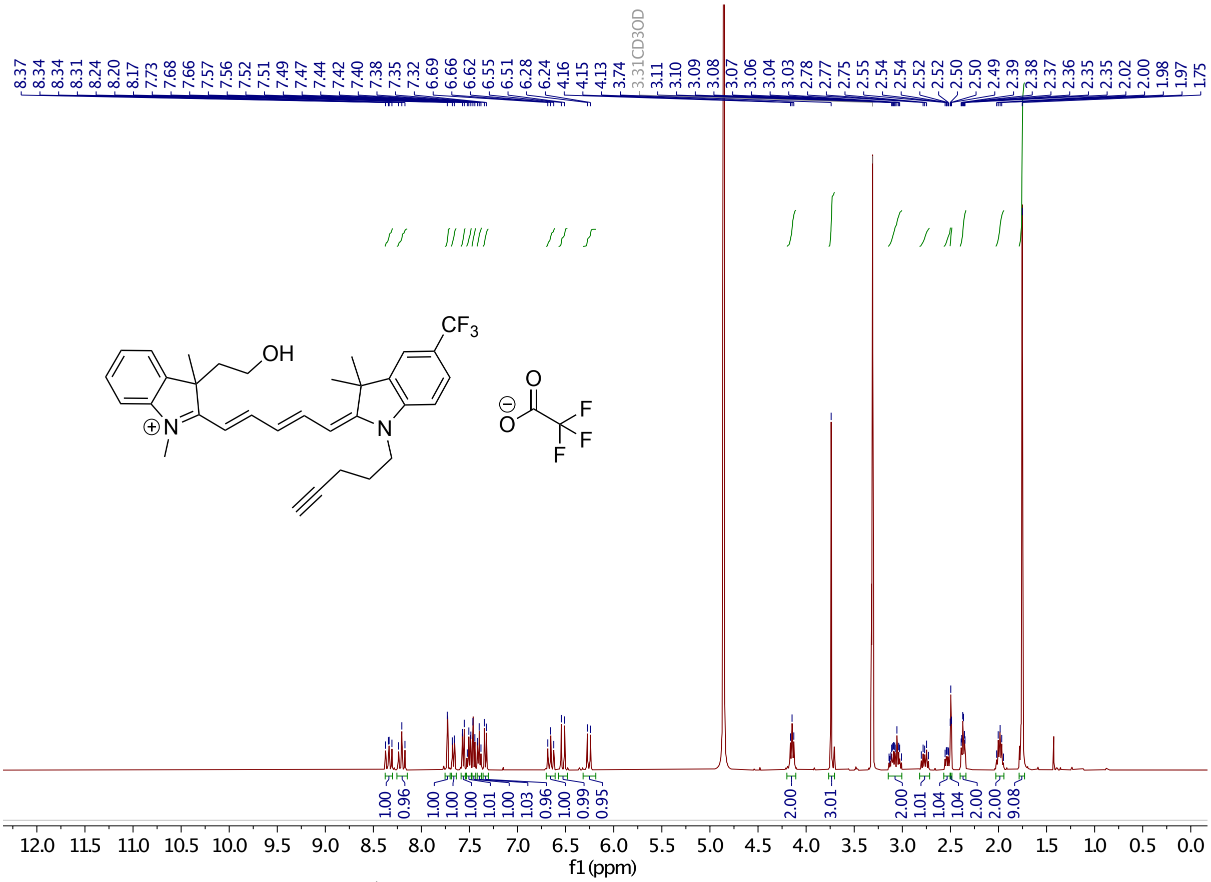

**Supplementary Figure 27.** $^{1}$H-NMR of **4c.**

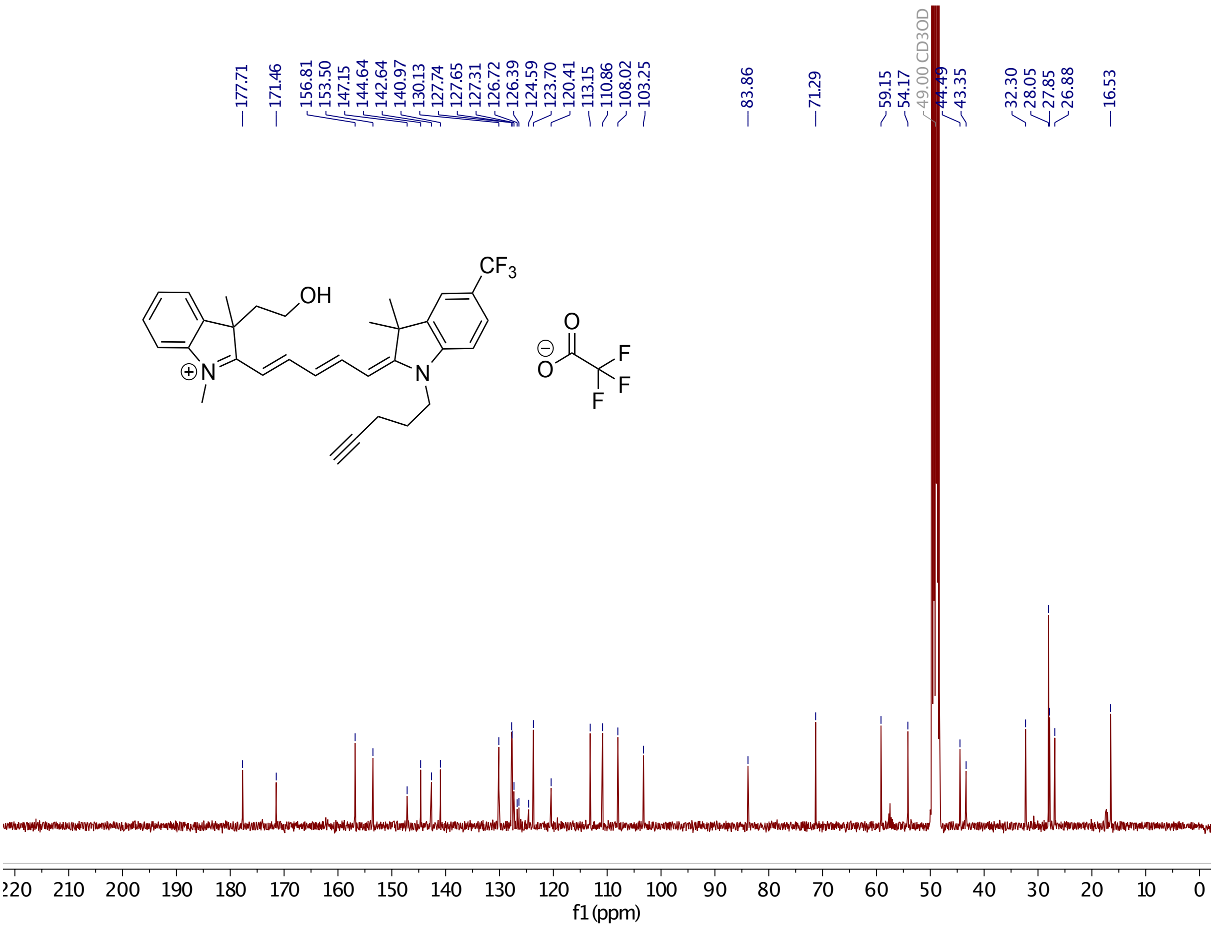

**Supplementary Figure 28.** $^{13}$C-NMR of **4c**.

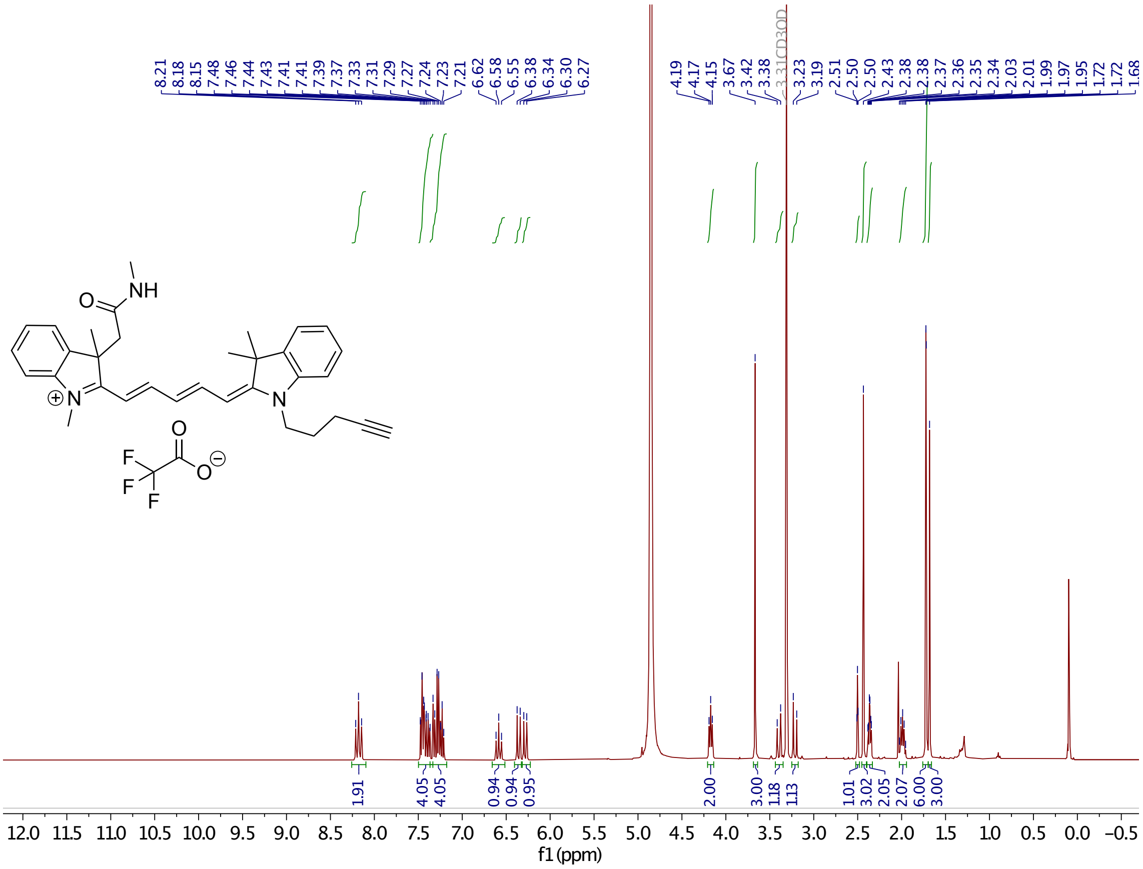

**Supplementary Figure 29.** $^1$H-NMR of **12**.

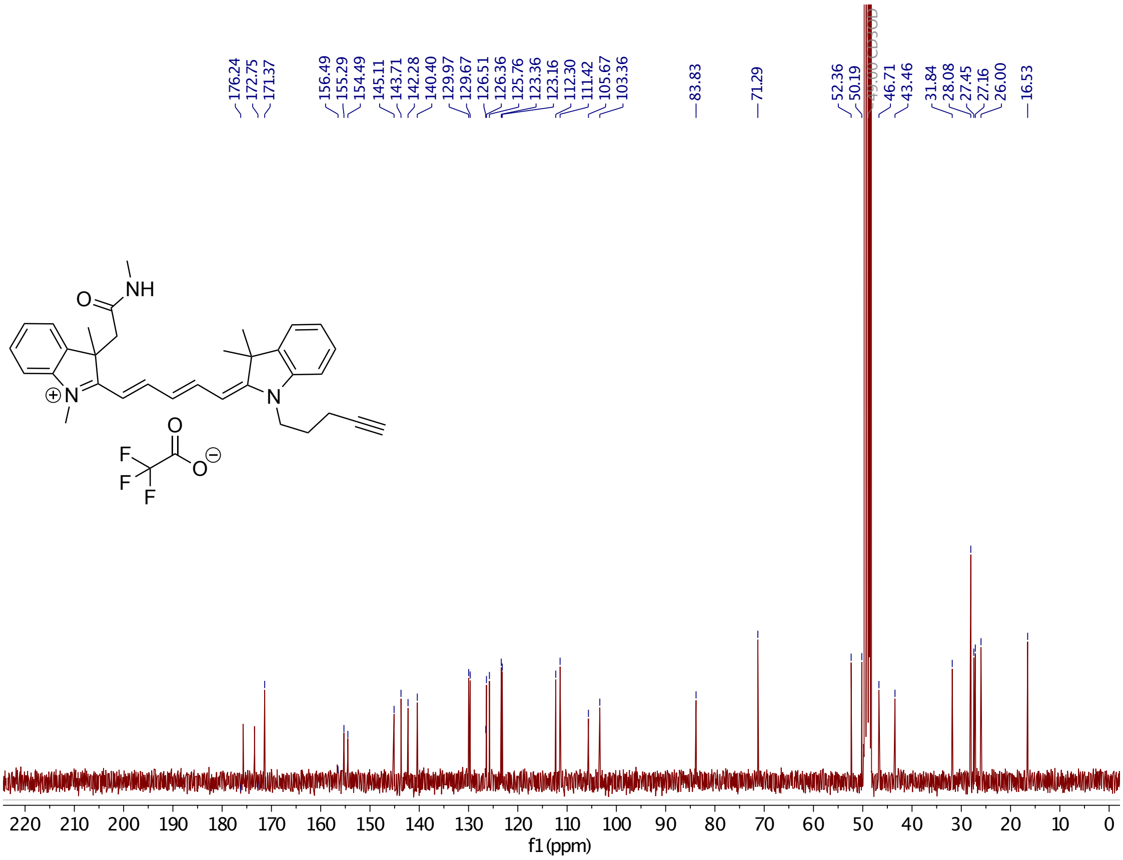

**Supplementary Figure 30.** $^{13}$C-NMR of **12**.

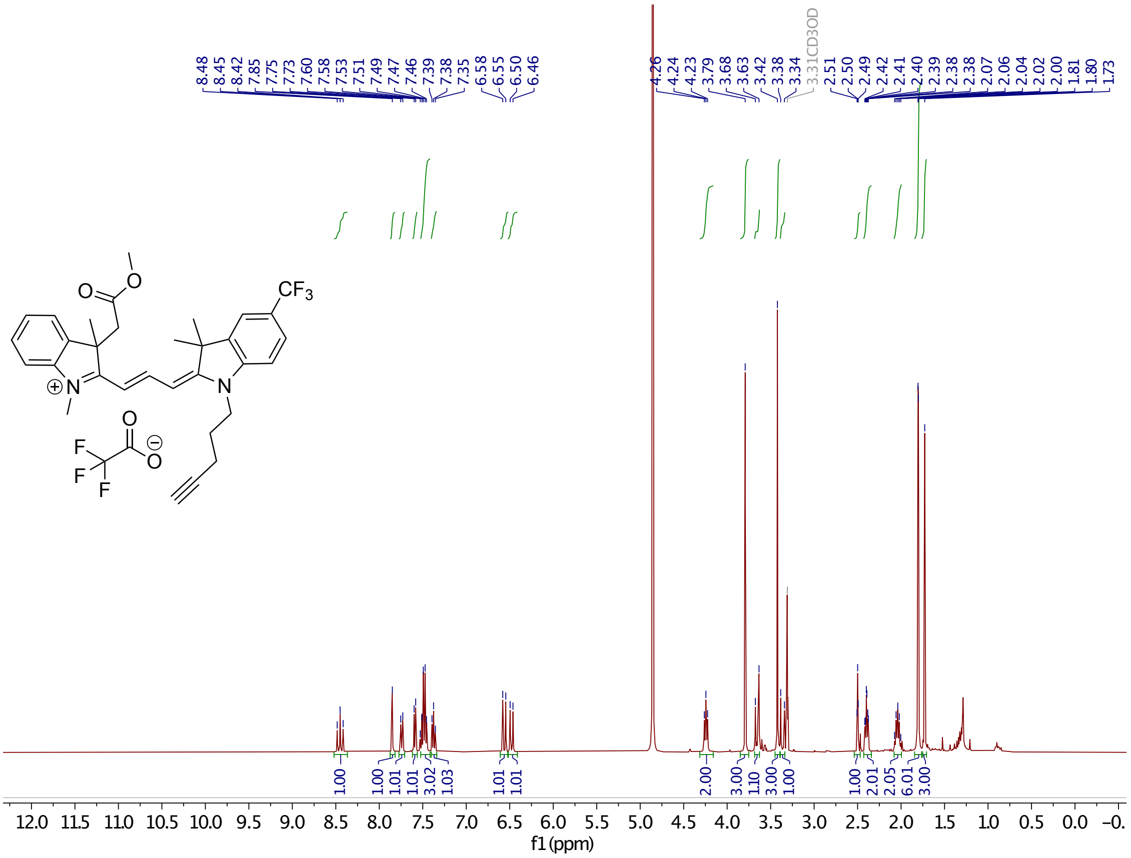

**Supplementary Figure 31.** $^{1}$H-NMR of **21**.

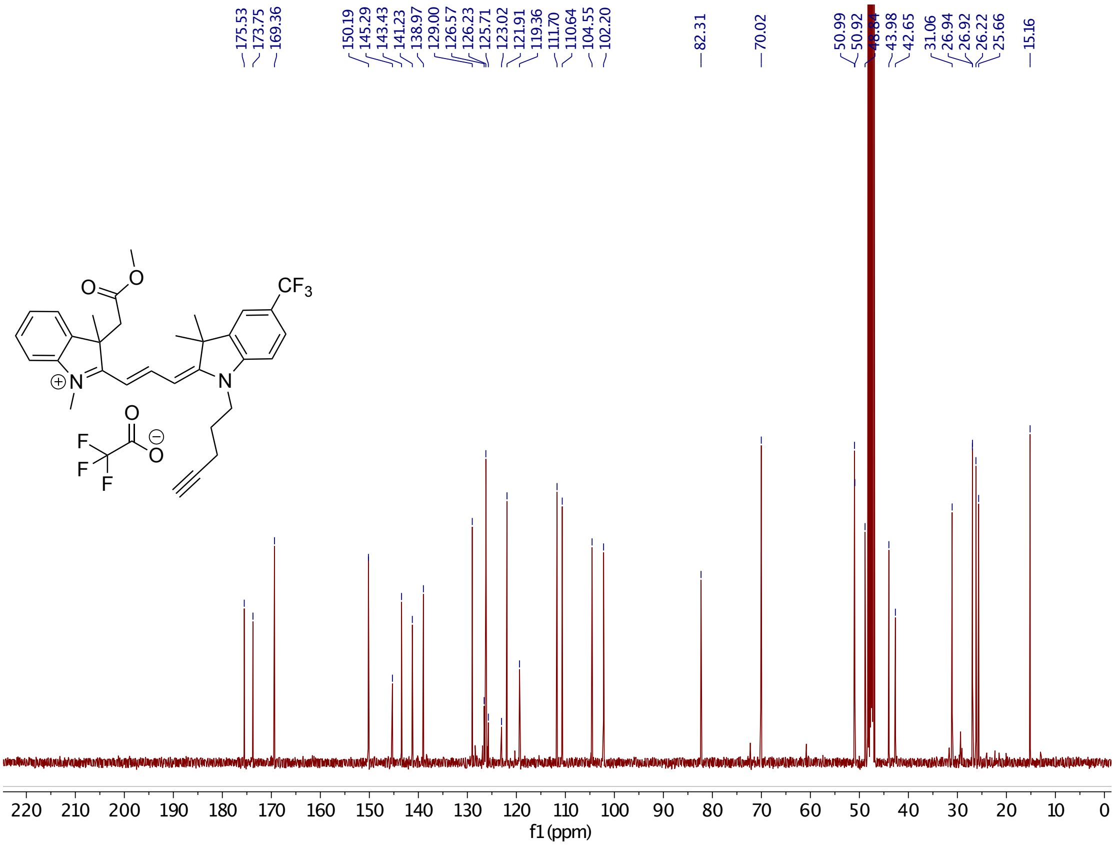

**Supplementary Figure 32.** $^{13}$C-NMR of **21**.

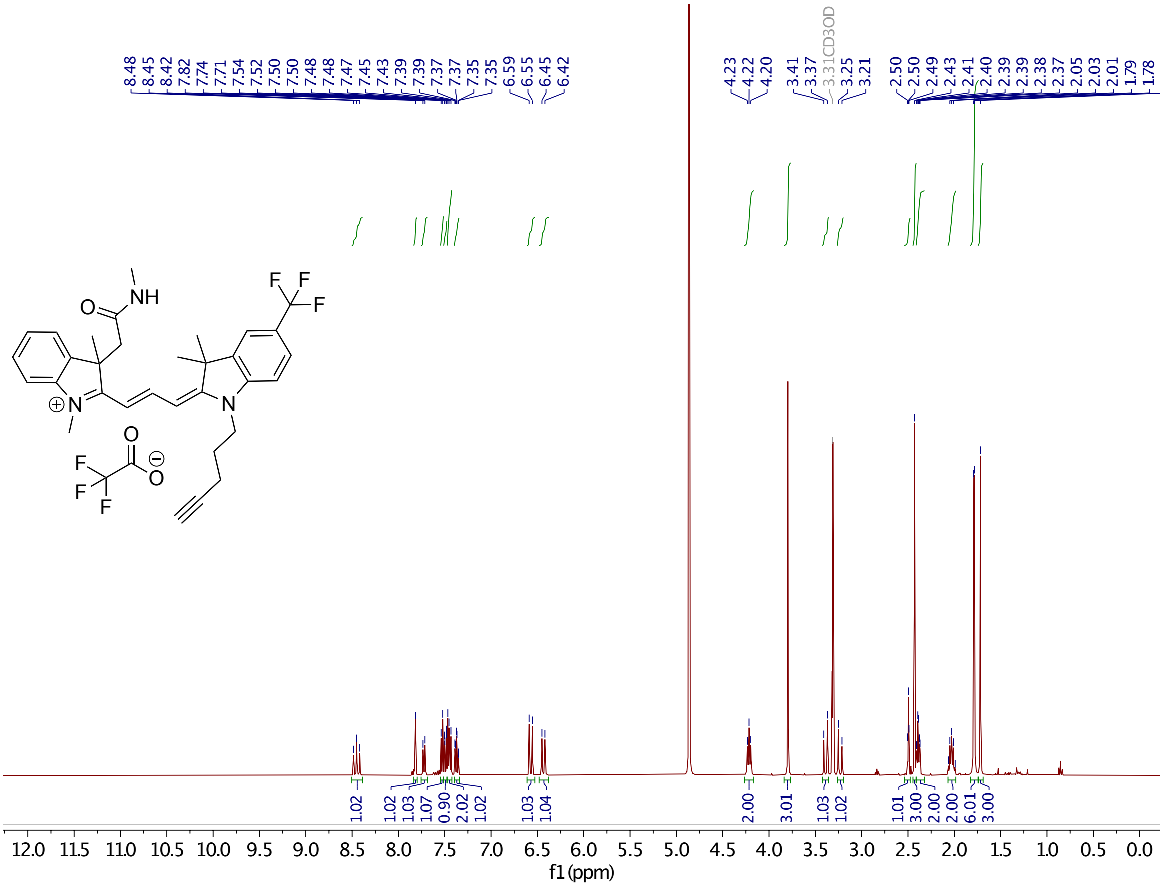

**Supplementary Figure 33.** $^{1}$H-NMR of **22**.

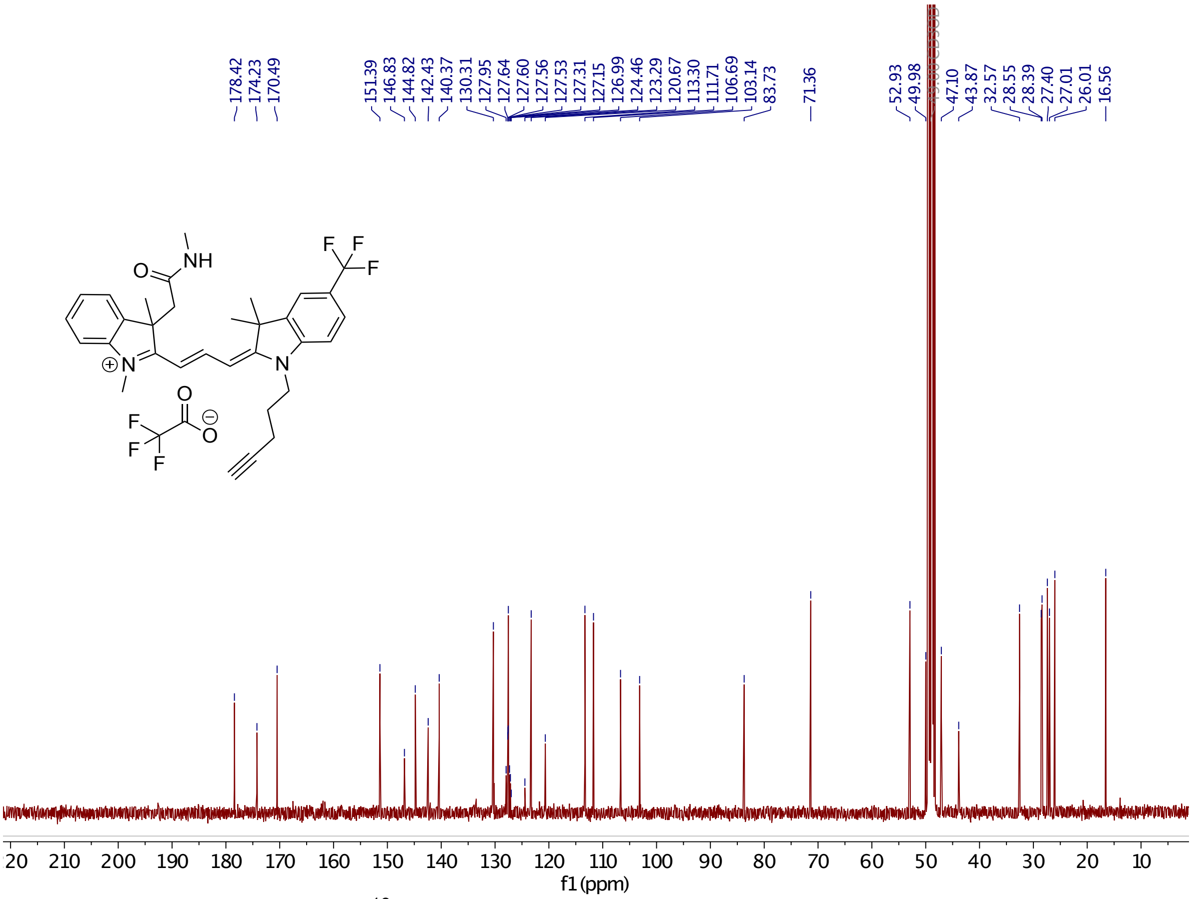

**Supplementary Figure 34.** $^{13}$C-NMR of **22**.

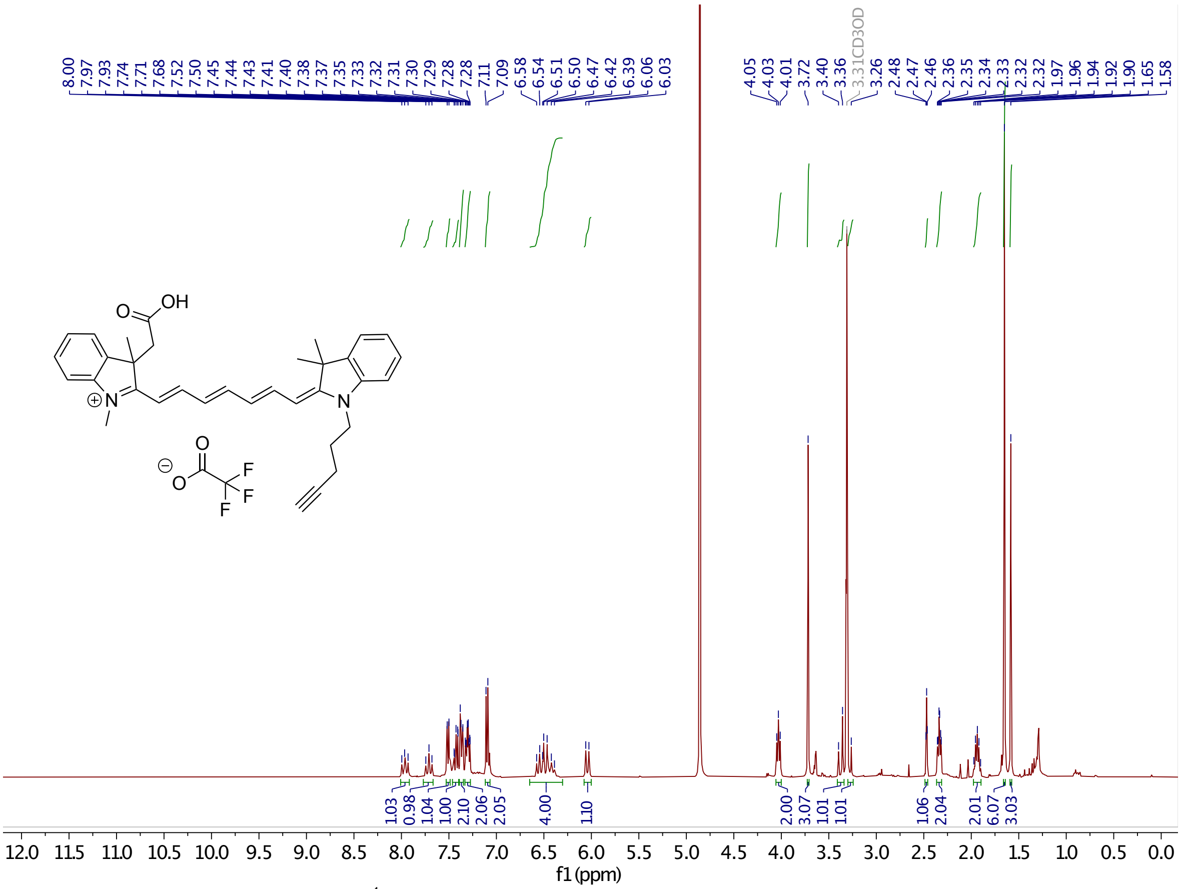

**Supplementary Figure 35.** $^{1}$H-NMR of **23**.

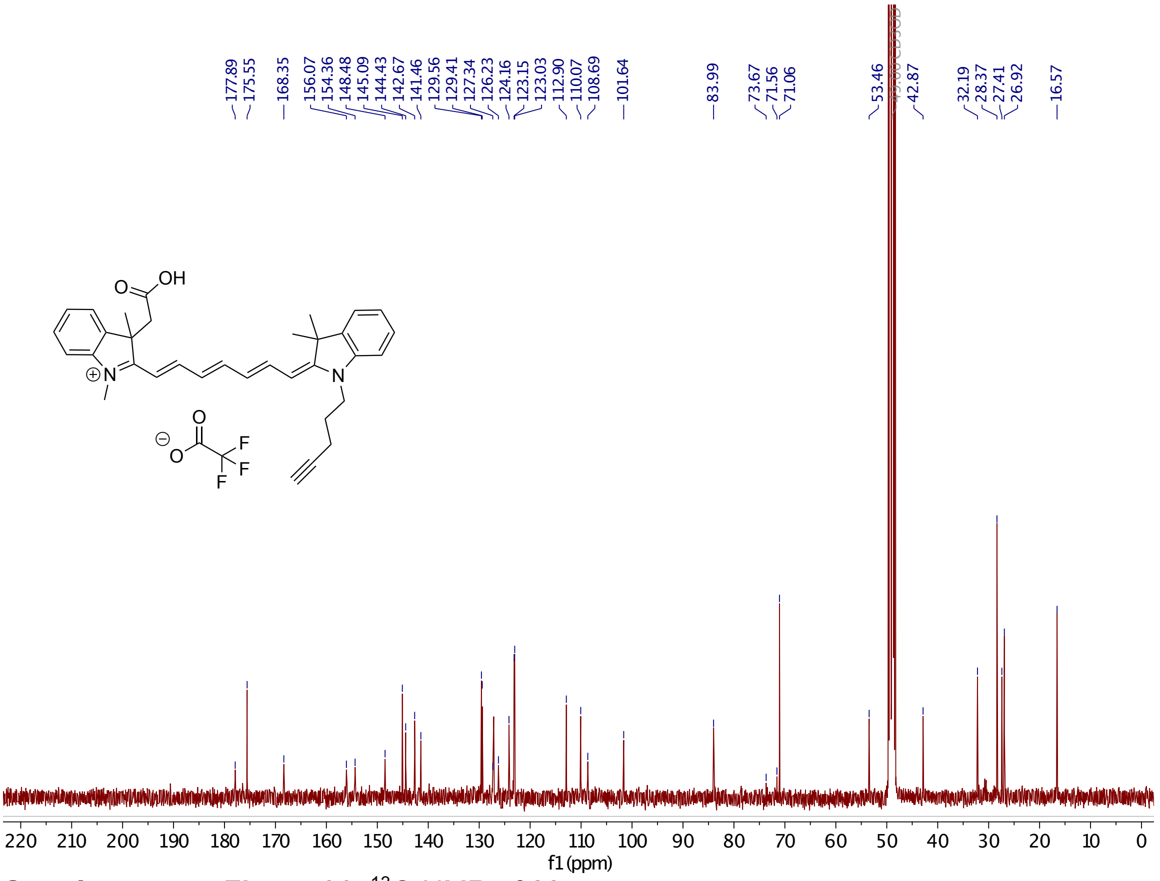

**Supplementary Figure 36.** $^{13}$C-NMR of **23**.

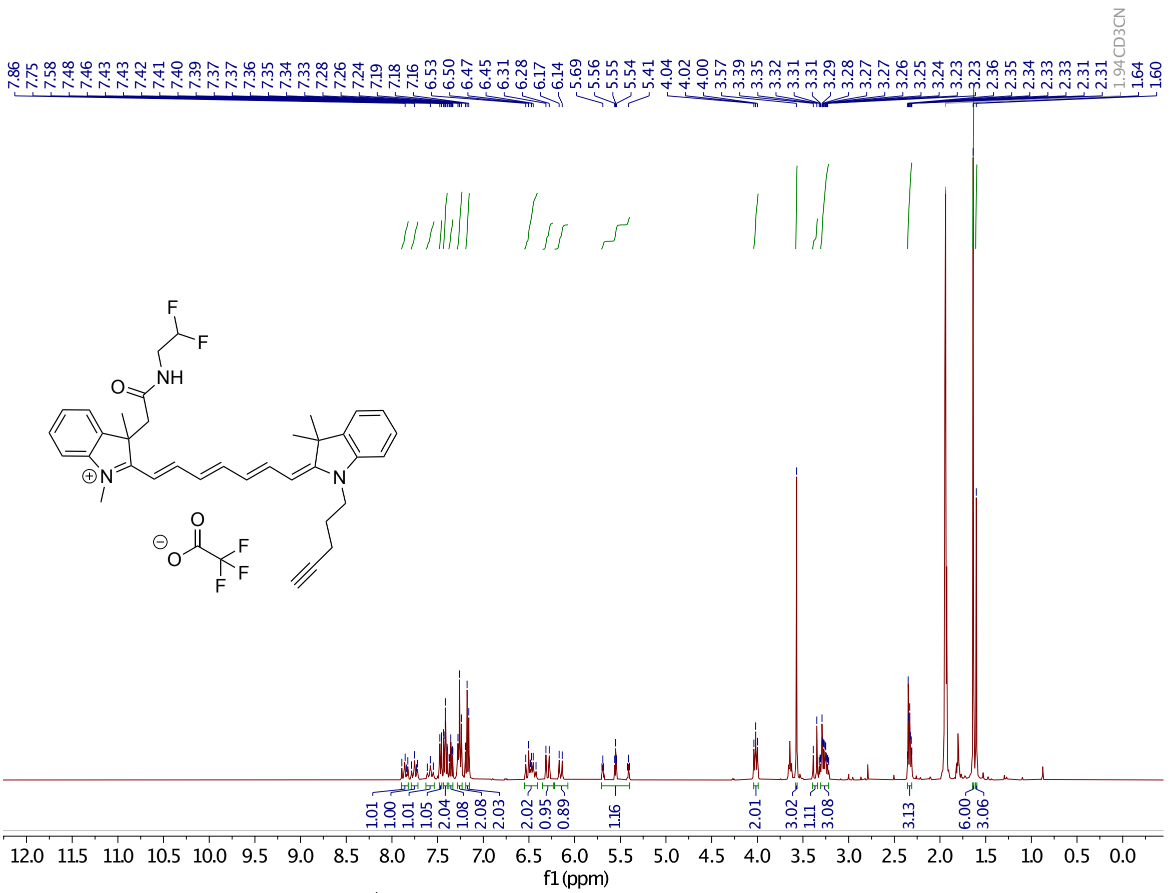

**Supplementary Figure 37.** $^{1}$H-NMR of **24**.

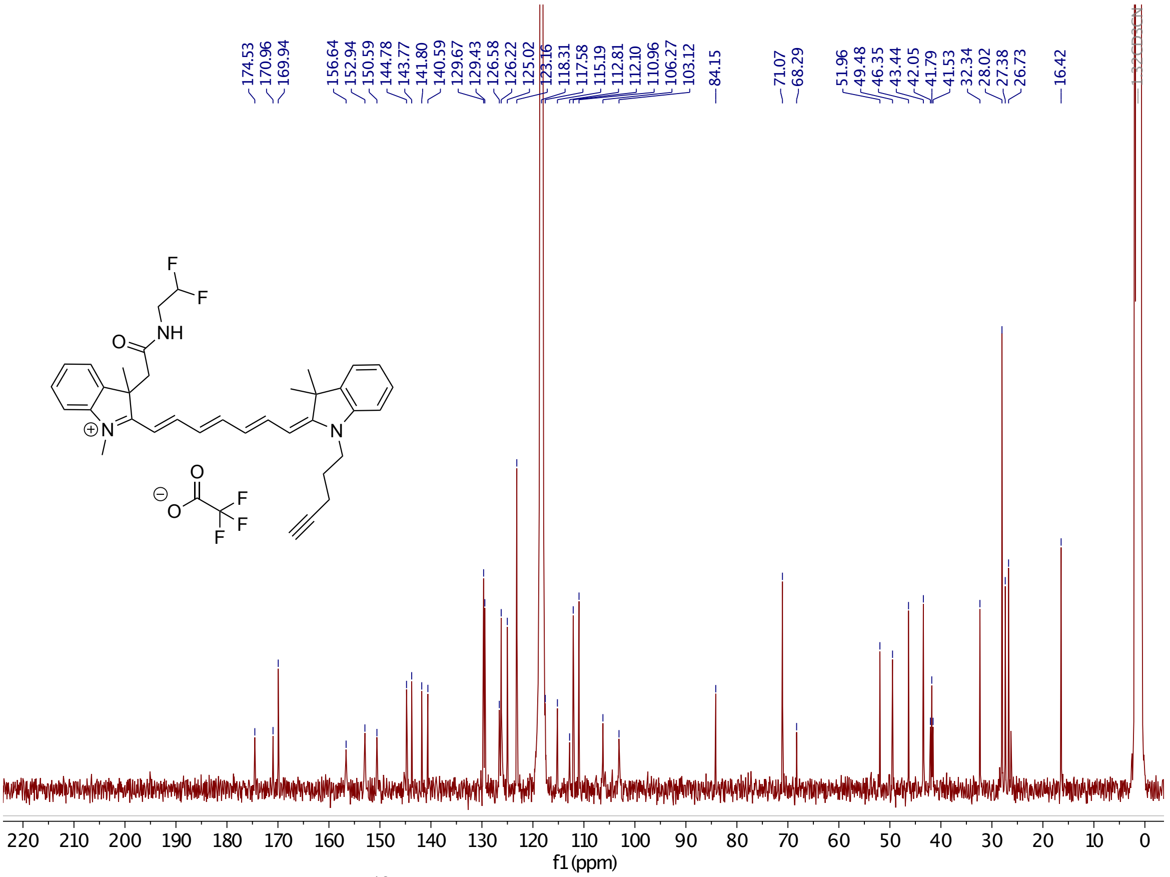

**Supplementary Figure 38.** $^{13}$C-NMR of **24**.

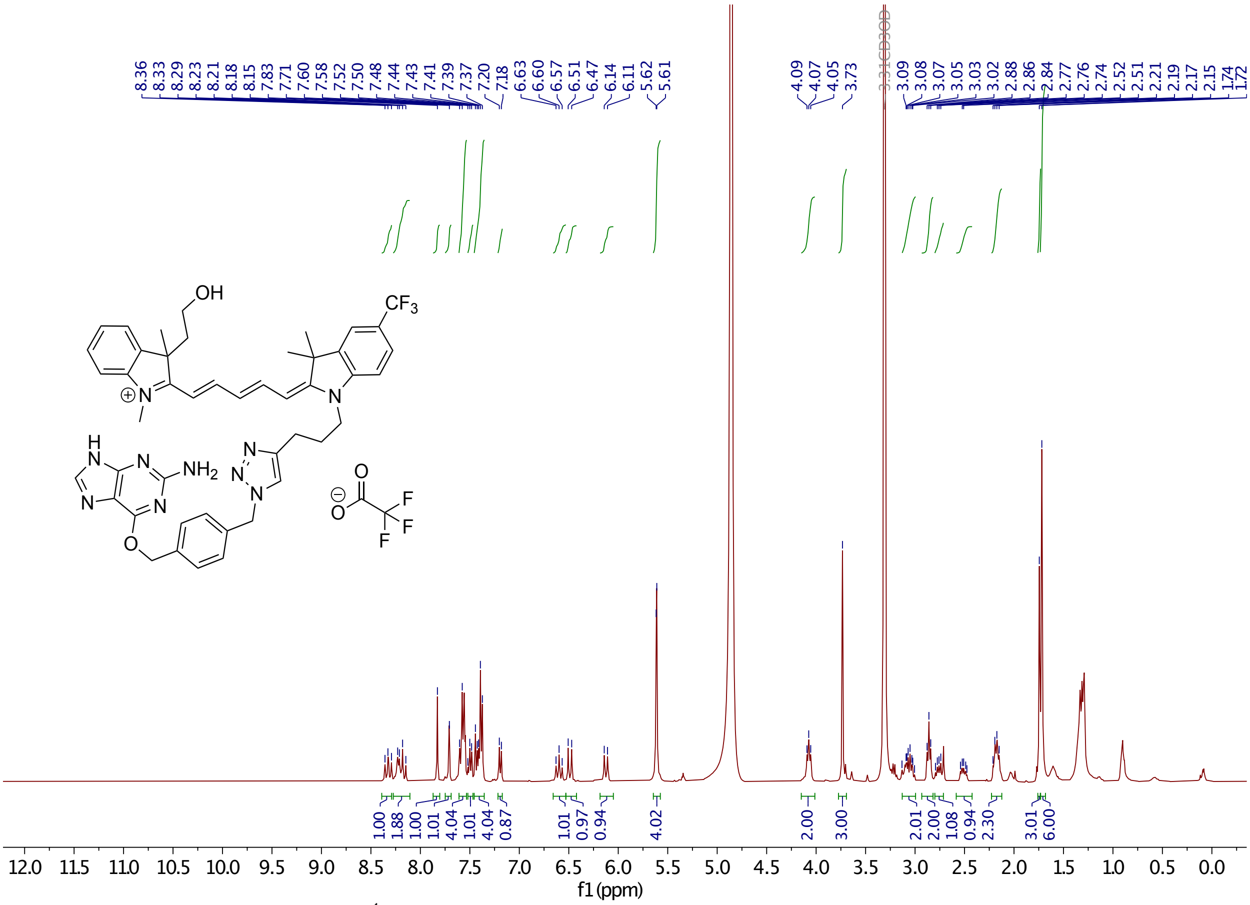

**Supplementary Figure 39.** $^{1}$H-NMR of **11**.

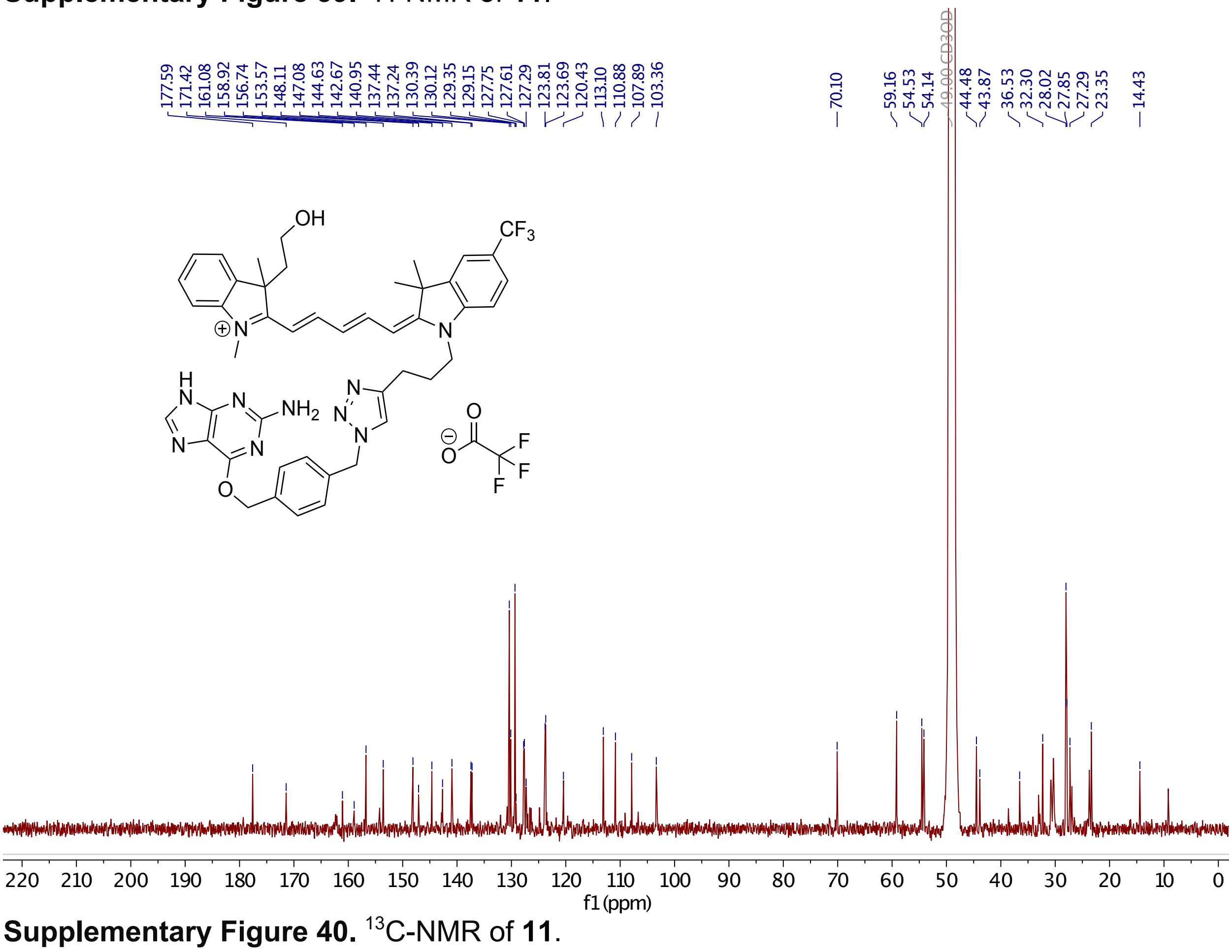

**Supplementary Figure 40.** $^{13}$C-NMR of **11**.

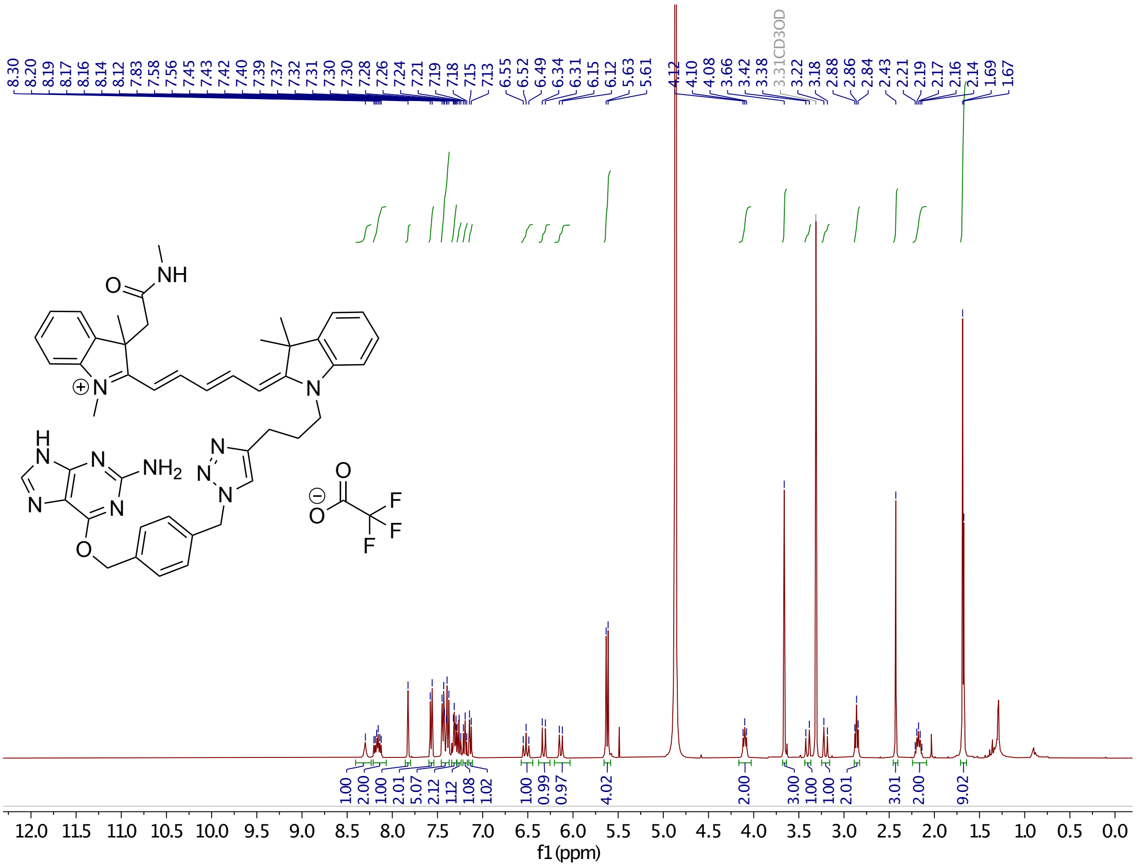

**Supplementary Figure 41.** $^{1}$H-NMR of **15**.

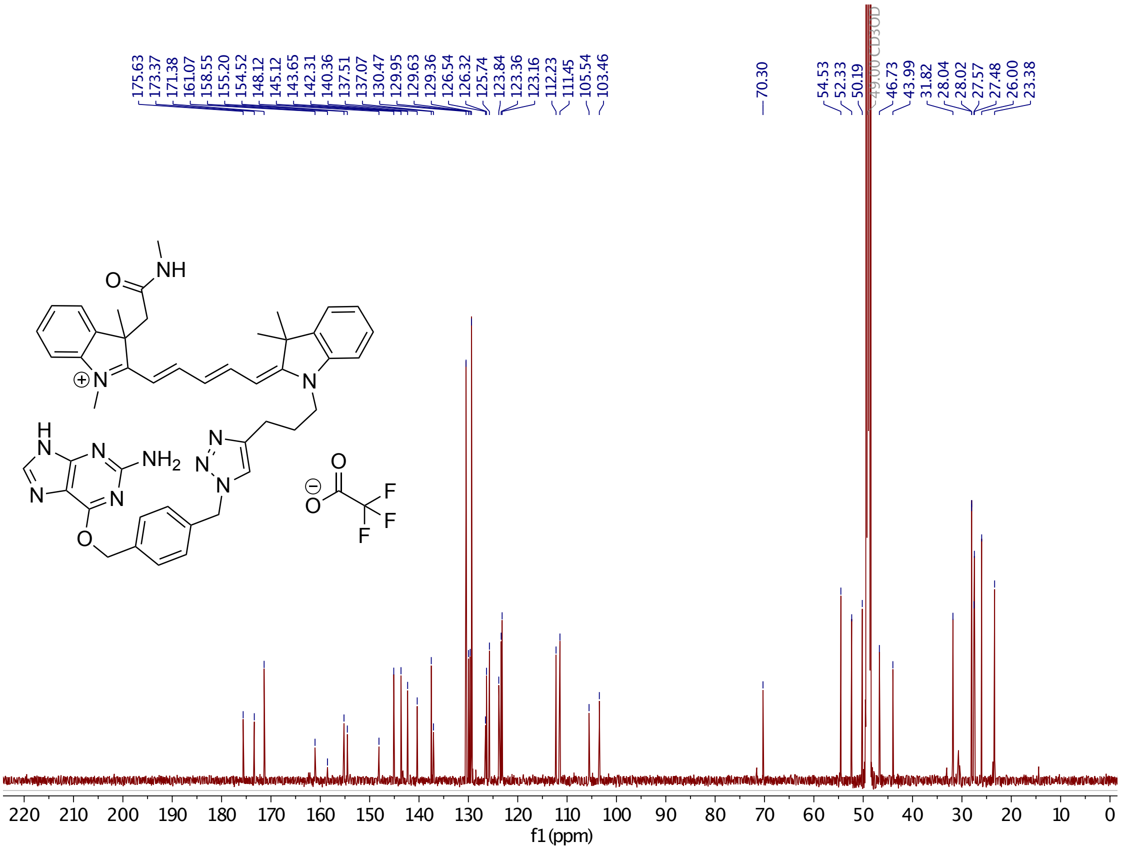

**Supplementary Figure 42.** $^{13}$C-NMR of **15**.

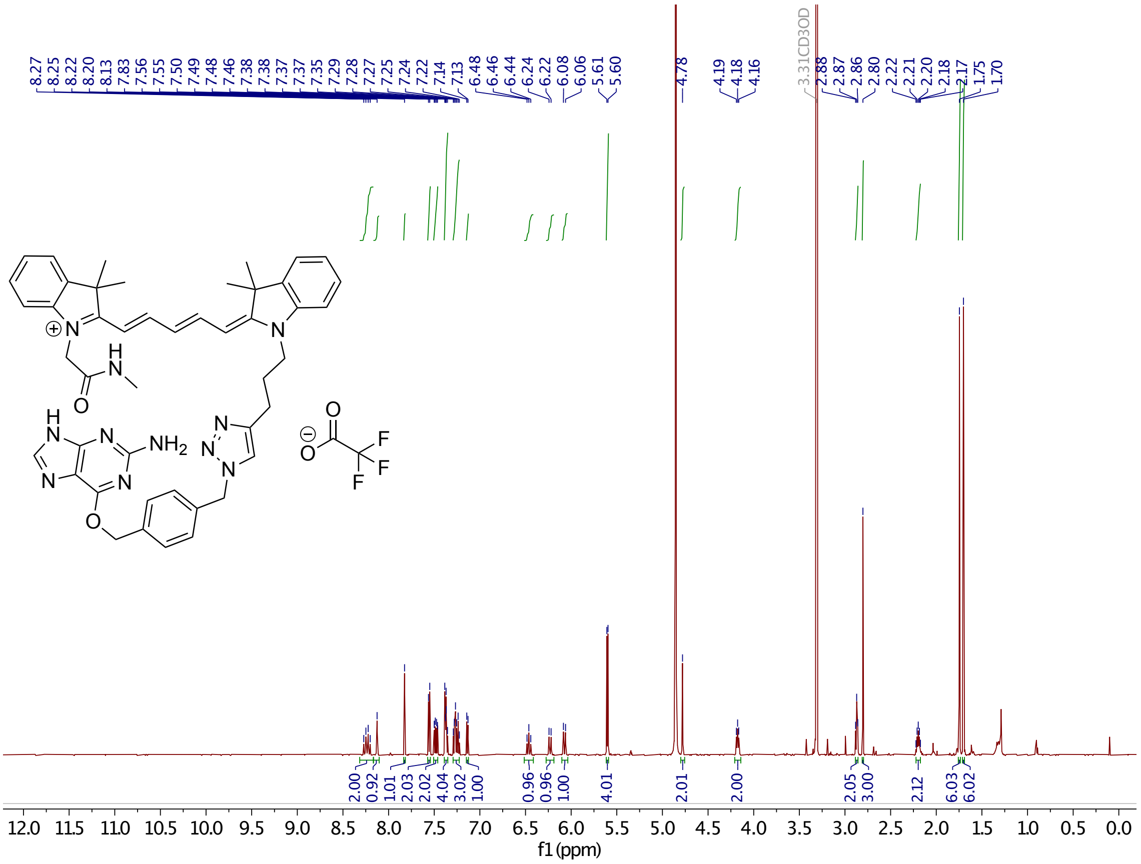

**Supplementary Figure 43.** $^{1}$H-NMR of **16**.

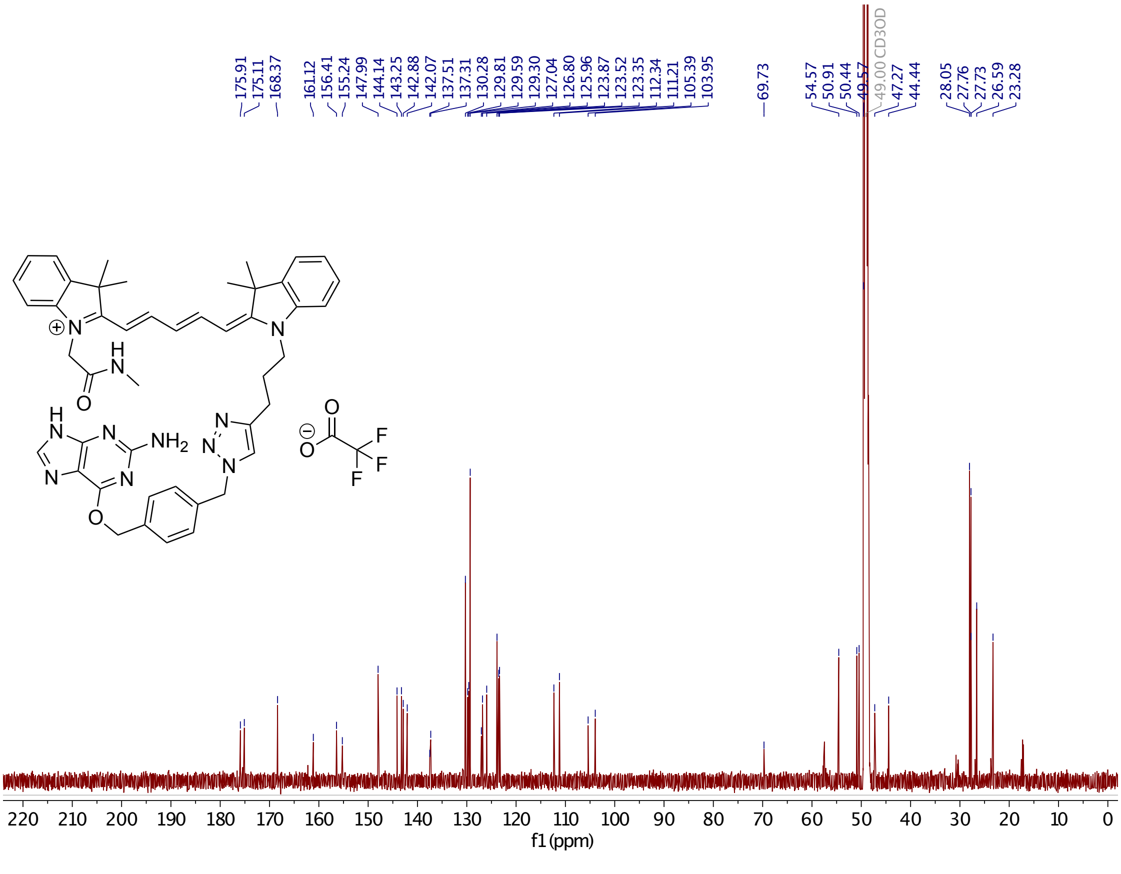

**Supplementary Figure 44.** $^{13}$C-NMR of **16**.

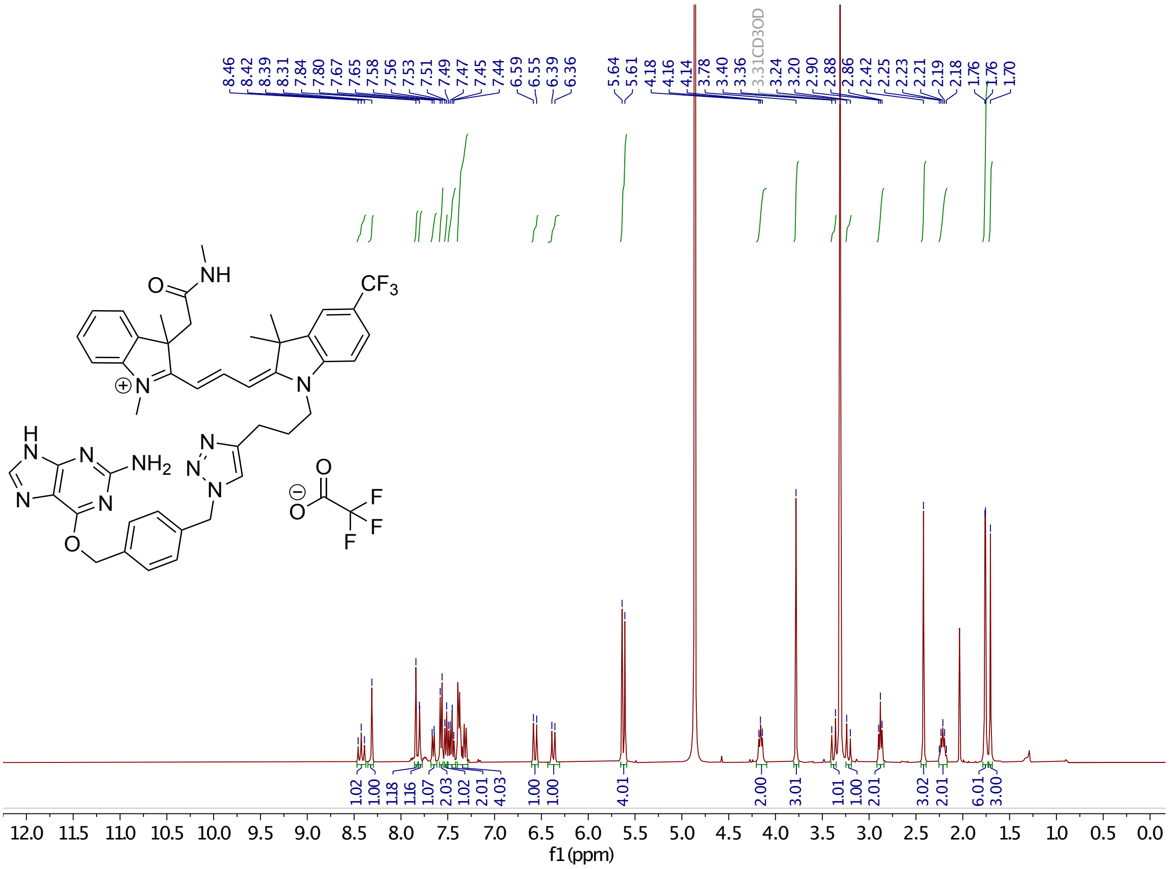

**Supplementary Figure 45.** $^{1}$H-NMR of **19**.

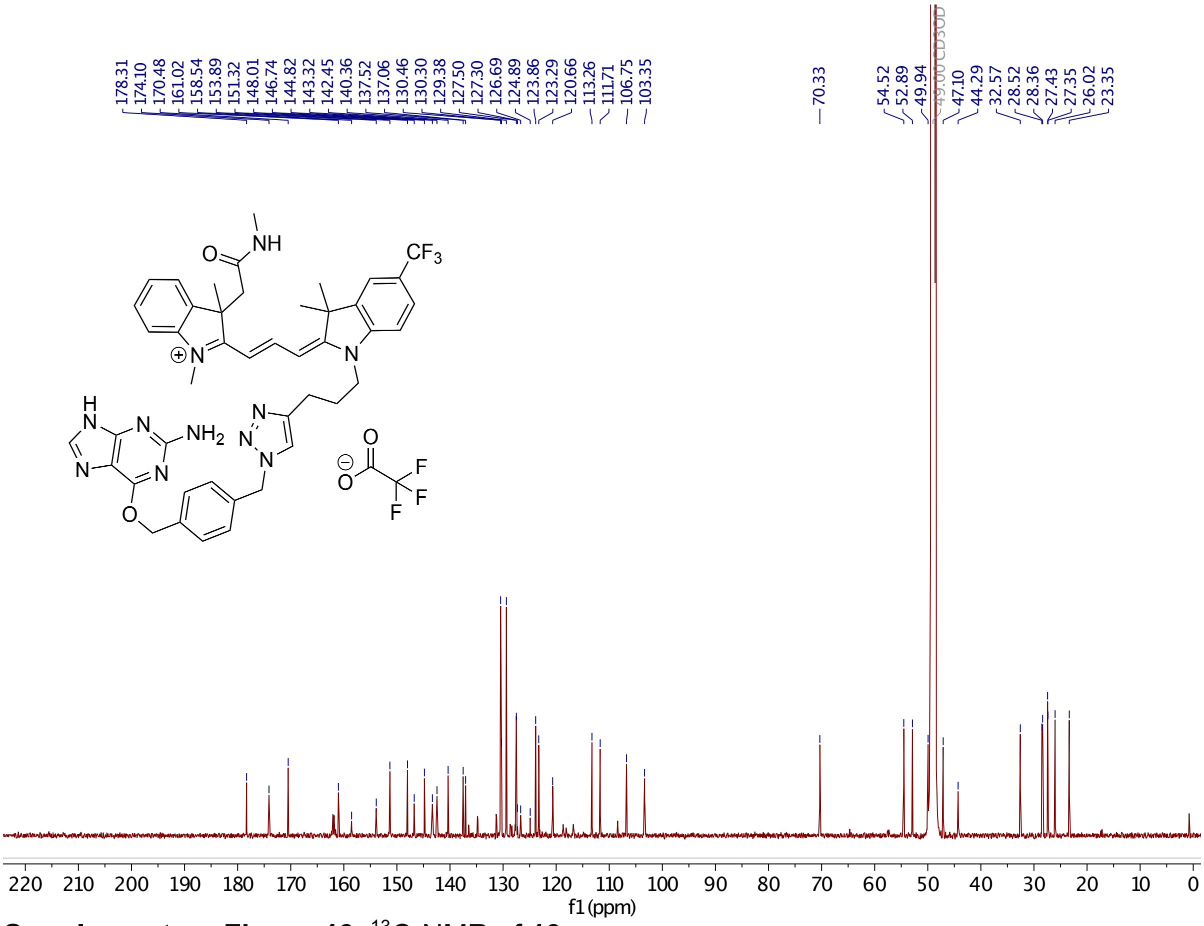

**Supplementary Figure 46.** $^{13}$C-NMR of **19**.

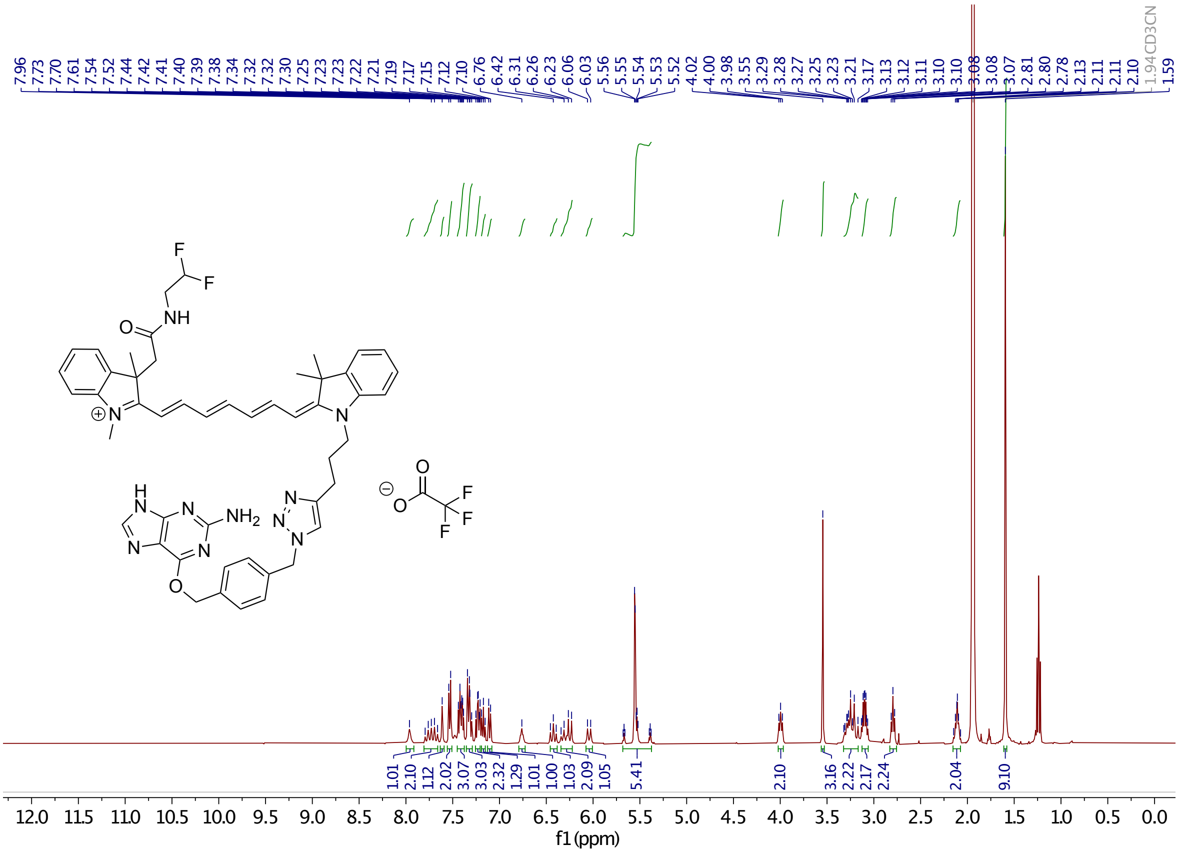

**Supplementary Figure 47.** $^1$H-NMR of **20**.

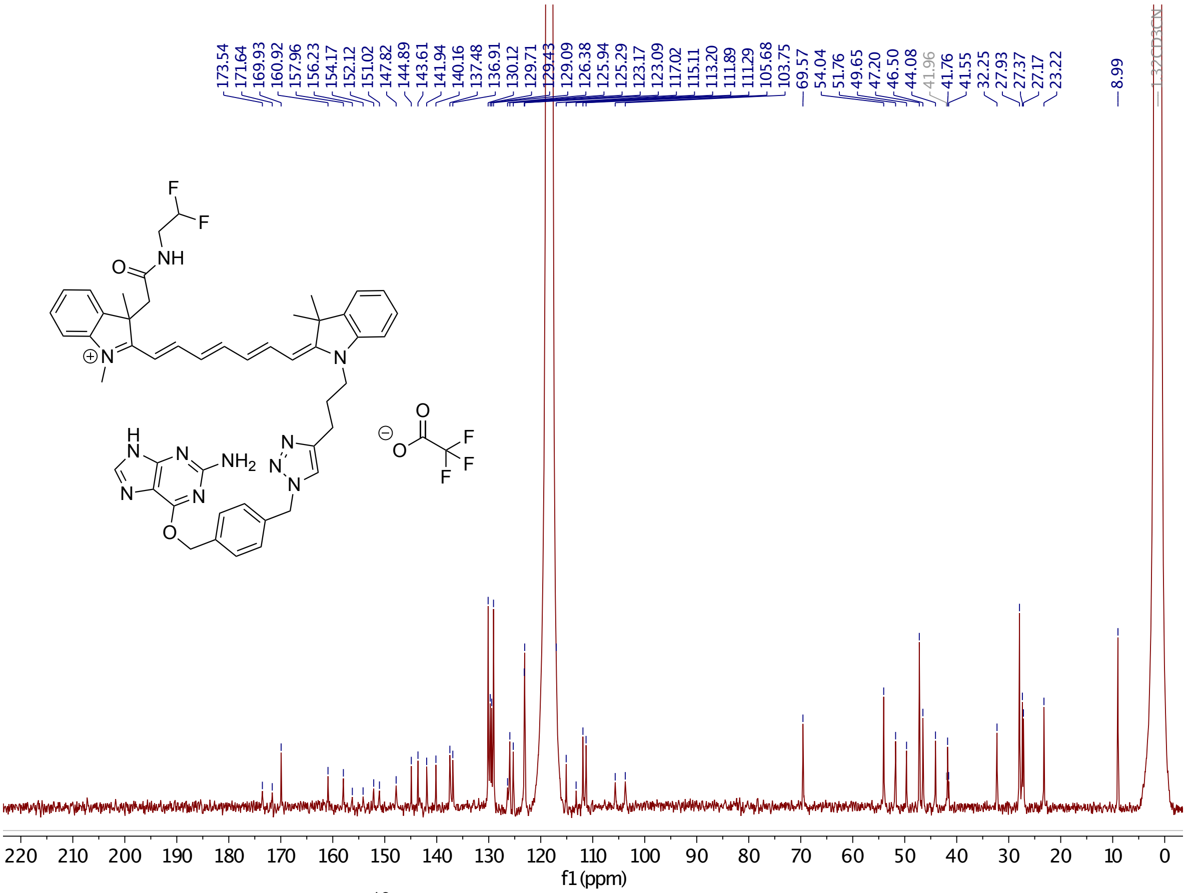

**Supplementary Figure 48.** $^{13}$C-NMR of **20**.

#### Plasmid maps

##### H2B-SNAPf-mTurquoise2

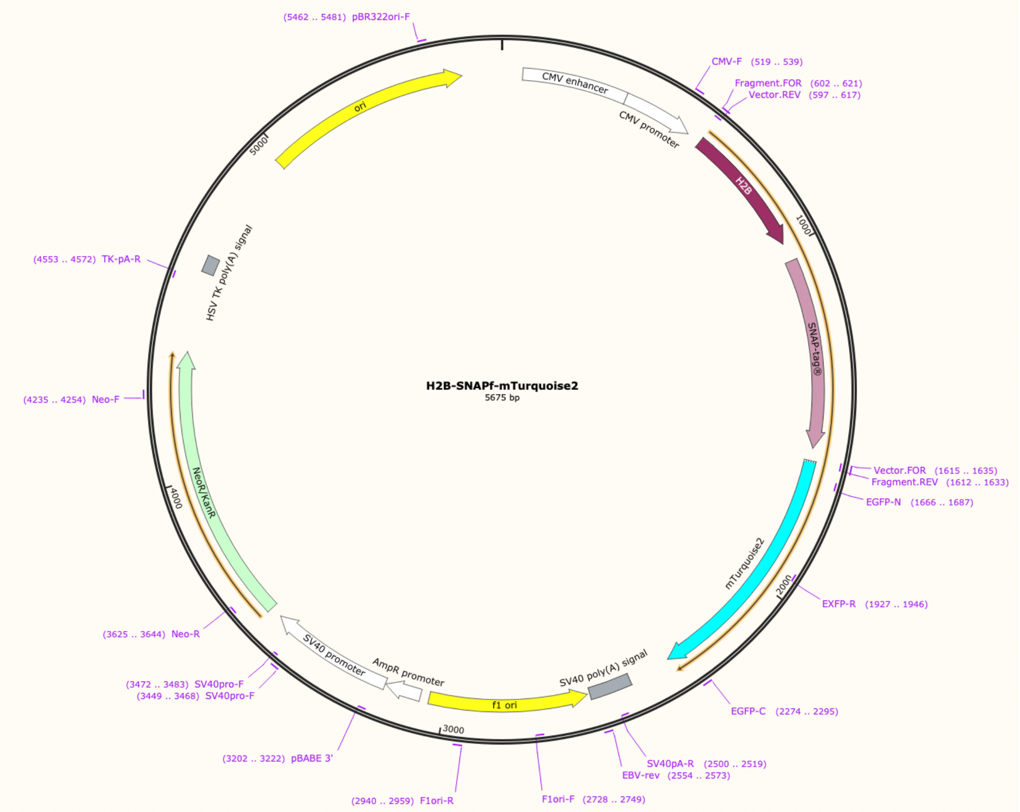

##### SNAPf-mTurquoise2-KDEL

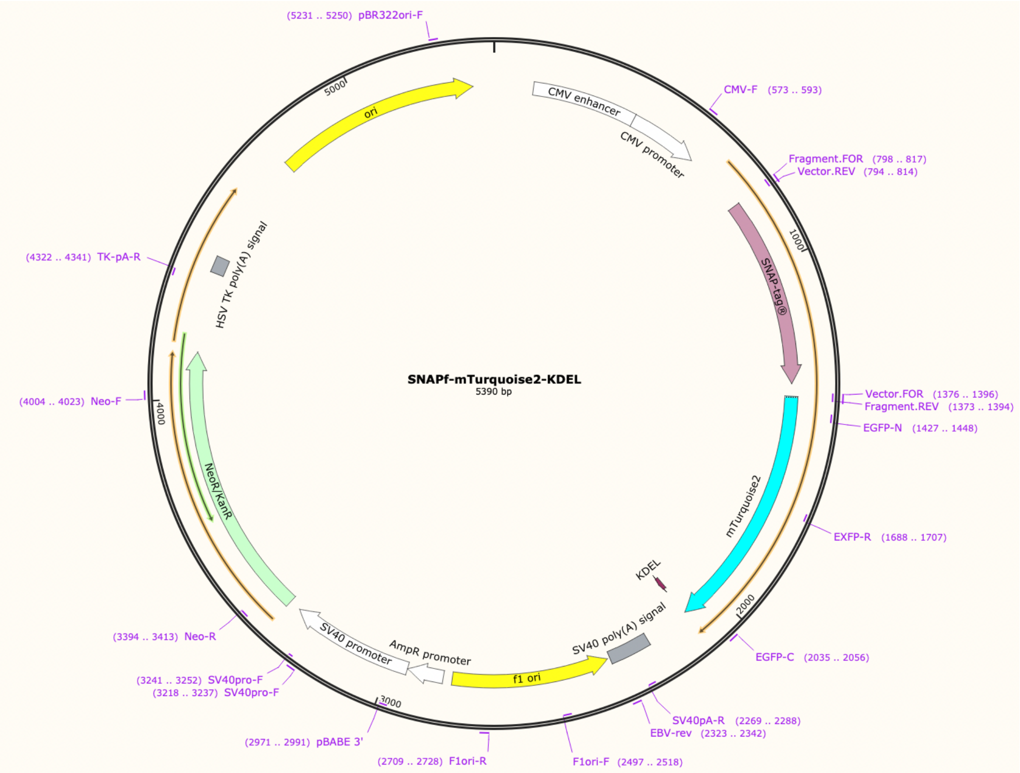

#### SNAPf-mTurquoise2

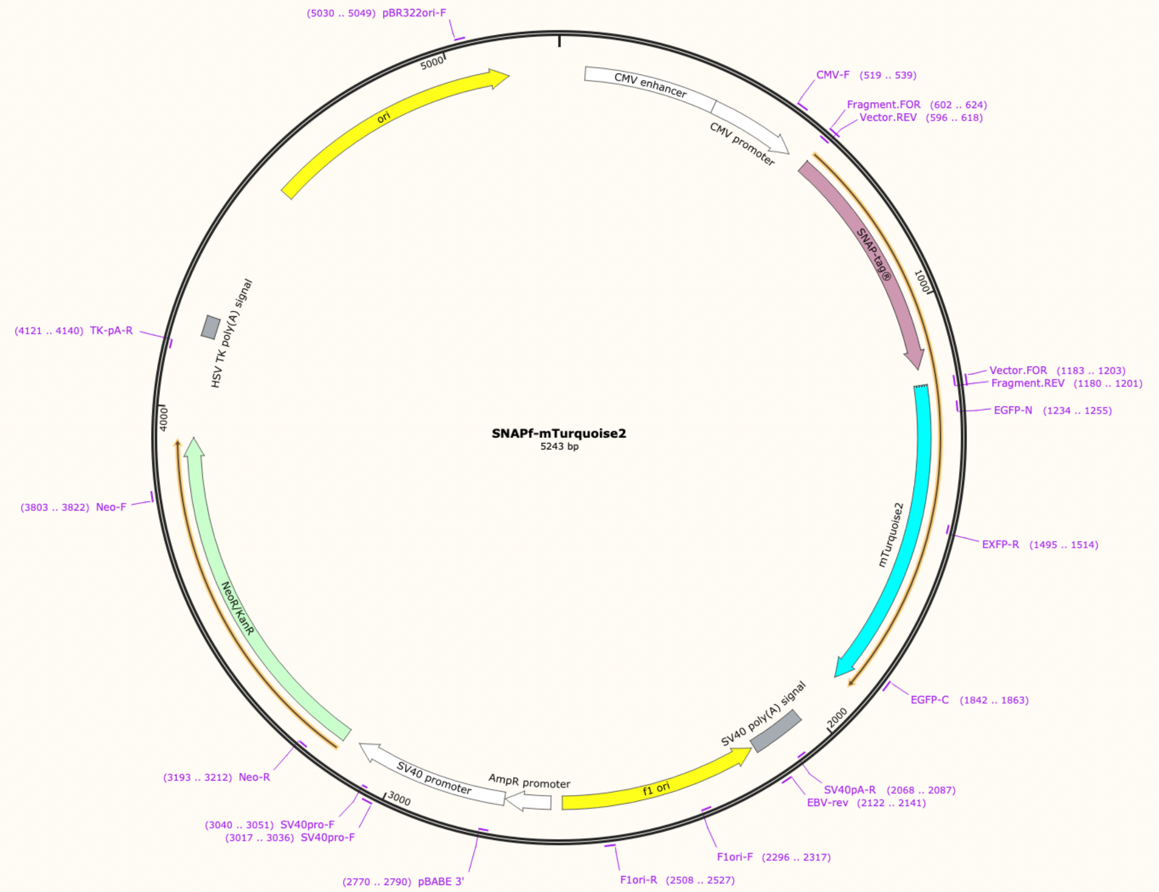

#### LifeAct-SNAPf-mTurquoise2

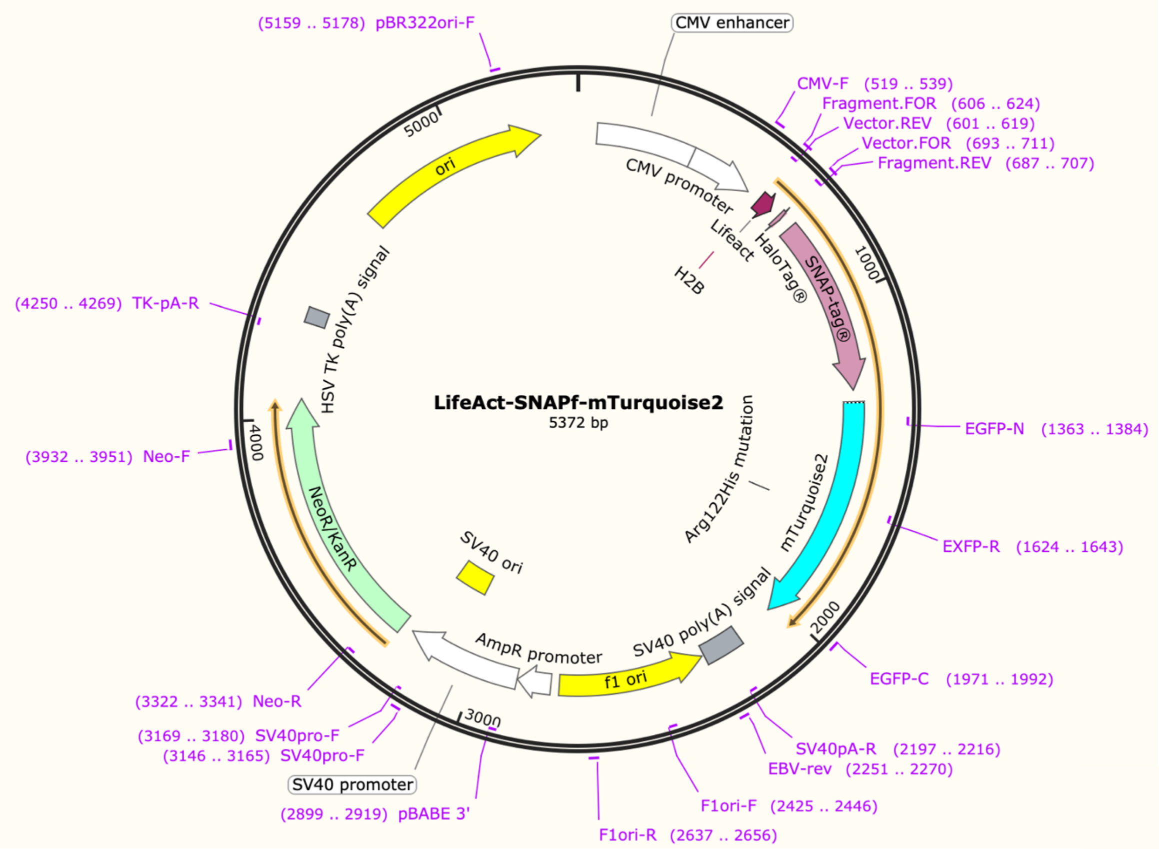

TOM20-SNAPf-mTurquoise2

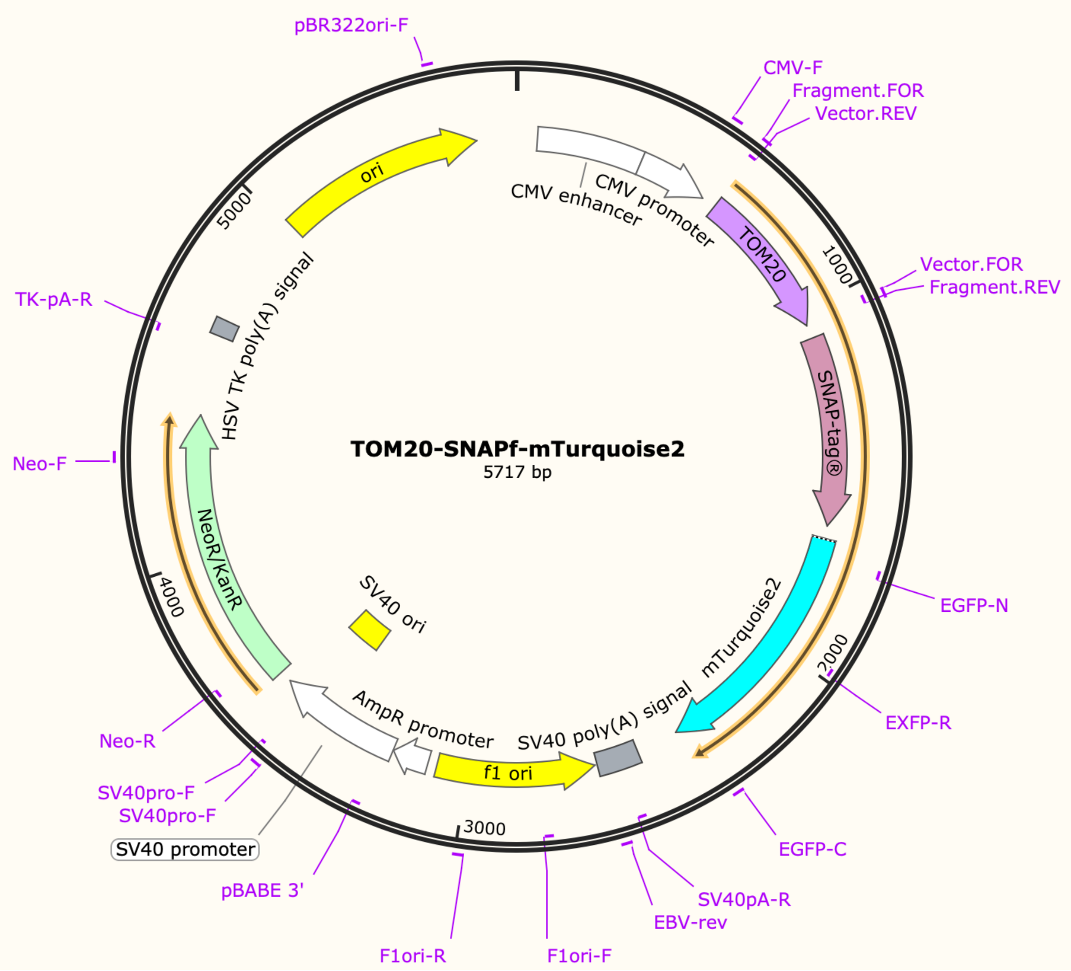
